# MiR-574-5p activates human TLR8 to promote autoimmune signaling and lupus

**DOI:** 10.1101/2023.09.06.556479

**Authors:** Tao Wang, Dan Song, Xuejuan Li, Yu Luo, Dianqiang Yang, Xiaoyan Liu, Xiaodan Kong, Yida Xing, Shulin Bi, Yan Zhang, Tao Hu, Yunyun Zhang, Shuang Dai, Zhiqiang Shao, Dahan Chen, Jinpao Hou, Esteban Ballestar, Jianchun Cai, Feng Zheng, James Y. Yang

## Abstract

Endosomal single-stranded RNA-sensing Toll-like receptor-7/8 (TLR7/8) plays a pivotal role in inflammation and immune responses and autoimmune diseases. However, the mechanisms underlying the initiation of the TLR7/8-mediated autoimmune signaling remain to be fully elucidated. Here, we demonstrate that miR-574-5p is aberrantly upregulated in tissues of lupus prone mice and in the plasma of lupus patients, with its expression levels correlating with the disease activity. miR-574-5p binds to and activates human hTLR8 or its murine ortholog mTlr7 to elicit a series of MyD88-dependent immune and inflammatory responses. These responses include the overproduction of cytokines and interferons, the activation of STAT1 signaling and B lymphocytes, and the production of autoantigens. In a transgenic mouse model, the induction of miR-574-5p overexpression is associated with increased secretion of antinuclear and anti-dsDNA antibodies, increased IgG and C3 deposit in the kidney, elevated expression of inflammatory genes in the spleen. In lupus-prone mice, lentivirus-mediated silencing of miR-574-5p significantly ameliorates major symptoms associated with lupus and lupus nephritis. Collectively, these results suggest that the miR-574-5p-hTLR8/mTlr7 signaling is an important axis of immune and inflammatory responses, contributing significantly to the development of lupus and lupus nephritis.

## INTRODUCTION

Toll-like receptors (TLRs) belong to a class of membrane-spanning pattern recognition receptors implicated in host defense and the development of a variety of human disorders ^1,2^. TLRs function by sensing danger signals such as the exogenous pathogen-associated molecular patterns from various microorganisms or the endogenous damage-associated molecular patterns, thereby initiating signaling pathways critical for the induction of innate and/or adaptive immune responses. More than a dozen TLRs have been found, and they appear to exhibit functional diversity while significantly differing in terms of cell type expression, cellular localization, ligand recognition and signal transduction ^1–3^. In humans, the cellular surface TLRs (TLR1/2/4/5/6/10) are responsible for the detection of specific microbial components expressed by bacteria, fungi, protozoa, or viruses. The endosomal TLRs (TLR3/7/8/9), on the other hand, serve primarily as sensors for viruses and nucleic acids from exogenous or endogenous sources, with TLR3 for double-stranded RNAs (dsRNAs), TLR7/8 for single-stranded RNAs (ssRNAs) and TLR9 for unmethylated CpG DNAs ^3^.

Human TLR7/8 (hTLR7/8) and mouse Tlr7/8 (mTlr7/8) are closely related phylogenetically, with their encoding genes located on the X chromosomes ^4^. However, despite the sequence similarity, these TLRs (hTLR7 versus hTLR8, mTlr7 versus mTlr8, hTLR7 versus mTlr7 and hTLR8 versus mTlr8) all differ significantly in tissue/cell expression, ligand binding, signaling and functions. In human, hTLR7 is primarily expressed by B cells and plasmacytoid dendritic cells (pDCs), whereas hTLR8 is found more in monocytes, myeloid DCs and neutrophils ^5^. Studies have shown that exogenous ssRNA sequences derived from viruses ^6,7^, fungi ^8^ or bacteria ^9^ or endogenous or self-ssRNAs present in living cells or released from dead or dying cells ^10–12^, synthetic oligoribonucleotides ^13^ and certain nucleoside analogs ^1,14,15^, and endogenous miRNAs can all serve as biological agonists for TLR7/8. However, the ligand recognition by TLR7/8 is often species-specific and receptor-specific. For example, nucleoside analog imiquimod (R837) only activates hTLR7 but not hTLR8, whereas resiquimod (R838) is a dual agonist for both hTLR7 and hTLR8 ^1,14^. Moreover, a GU-rich sequence from the U5 region of HIV-1 RNA (RNA40) is recognized by hTLR8 and mTlr7 but not hTLR7 ^6^ whereas mTlr8 is not responsive to the known RNA ligands for hTLR7/8 and mTlr7. Stimulation of TLR7/8 by exogenous or endogenous ligands triggers myeloid differentiation primary response gene-88 (MyD88)-dependent or -independent signaling pathways, eventually leading to the synthesis and secretion of interferons and pro-inflammatory cytokines or chemokines and the activation of immune cells^2^. However, hTLR7 activation predominantly induces interferons (IFNs) and IFN-induced cytokine expression in pDCs, whereas hTLR8 activation mainly elicits high level of proinflammatory cytokines in monocytes, monocyte-derived DCs, and myeloid DCs ^16^.

Dysregulation in TLR7/8 signaling is believed to be closely associated with viral and microbial infections as well as the development of neurodegeneration, cancers and autoimmune diseases including systemic lupus erythematosus (SLE) and rheumatoid arthritis, asthma, psoriasis and Type I diabetes mellitus ^1,11,17–19^. Endogenous ssRNA ligands for TLR7/8 were previously proposed to have a role in autoantibody production and autoimmunity ^12,20–22^, yet the identity and nature of these ssRNAs were not well-defined. Recent studies suggest that exosome-derived miR-574-5p might play important roles in immune and inflammation responses, autoimmune diseases and cancer through activating TLR7/8 ^23–26^. Fabbri et al. pioneered the elucidation that miRNAs are able to serve as ligands for TLR7/8 to induce prometastatic inflammatory response in cancer cells. While they were demonstrating miR-21 and miR-19-induced activation of hTLR8 and mTlr7 but not hTLR7 leading to induction of CD69 and NFκB activity and the increased production of TNFα and Il6, they also observed increased production of TNFα and Il6 following transfection of miR-574-5p in human PBMC cells, although at that moment it was not clear which TLR miR-574-5p could interact with ^23^. Surprisingly, by using synthetic microRNAs, Salvi et al. identified exosome-derived miR-574-5p as a hTLR7 endogenous ligands able to induce pDC activation ^26^. Nevertheless, their transfection studies indicated that miR-574-5p activated both hTLR7 and hTLR8 in the NFκB-luciferase reporter HEK293 cells, suggesting that hTLR7 is not the only one activated by miR-574-5p. Indeed, Hegewald et al. clearly demonstrated the physical binding between miR-574-5p and hTLR8 ^24^. In addition, they showed that synovial fluid-derived small extracellular vesicles contained high levels of miR-574-5p and that these miR-574-5p containing vesicles induce osteoclastogenesis. Since both hTLR7 and hTLR8 are believed to be implicated in the autoimmunity ^5,27–29^, it is essential to clarify whether and how miR-574-5p-induced hTLR8 activation contributes to the pathogenesis or progression of SLE.

Herein, we found that miR-574-5p, which was upregulated in both human and mouse lupus, acted as a potent ligand for mouse mTlr7 and human hTLR8 but not hTLR7. Aberrant activation of miR-574-5p-hTLR8/mTlr7 signaling was found to be associated with significant dysregulation in immune and inflammatory responses, especially the activation of the B cells. Meanwhile, transgenic expression of miR-574-5p was found to induce serum secretion of anti-nuclear antibody, to increase the renal deposit of complement C3 and IgG and the spleen expression of a few inflammation-related genes, whereas *in vivo* silencing of miR-574-5p greatly alleviated SLE and lupus nephritis in the lupus-prone B6.MRL-*Fas^lpr^*/J mice.

## RESULTS

### miR-574-5p is upregulated in SLE patients and lupus-prone mice and correlates with disease severity

Previous microarray studies indicated that miR-574-5p is among the miRNAs that was significantly upregulated in the plasma from SLE patients and rheumatoid arthritis patients ^30^. To ascertain the aberrant upregulation of miR-574-5p in SLE, we assessed the levels of serum and tissues miR-574-5p in lupus patients and lupus-prone mice. Using qPCR analyses, we showed that in comparison with that of 18 normal subjects, serum miR-574-5p was significantly elevated in a cohort of 47 SLE patients (Figure 1***a***). Furthermore, the serum levels of miR-574-5p in the SLE patients were found to correlate positively with the clinical SLE disease activity index (SLEDAI) (Spearman r = 0.3986, *P* < 0.01, Figure 1***b*** & supplemental Table S1). miR-574-5p expression was highly elevated in the serum and kidney of 90-d old lupus-prone B6.*Fas^lpr^* mice and persisted up to 180-d (Figure 1***c***-***d***), although no significant differences were observed in 60-d. Meanwhile, the levels of miR-574-5p in spleen, liver, heart, lung, brain and lymph node were also sharply elevated in the 180-d old lupus-prone mice (Figures 1***e***). The elevation of miR-574-5p in various tissues also correlates well with the onset of lupus nephritis in the mice. Taken together, these results indicate a strong link between miR-574-5p and the pathogenesis and development of SLE.

**Figure 1.**
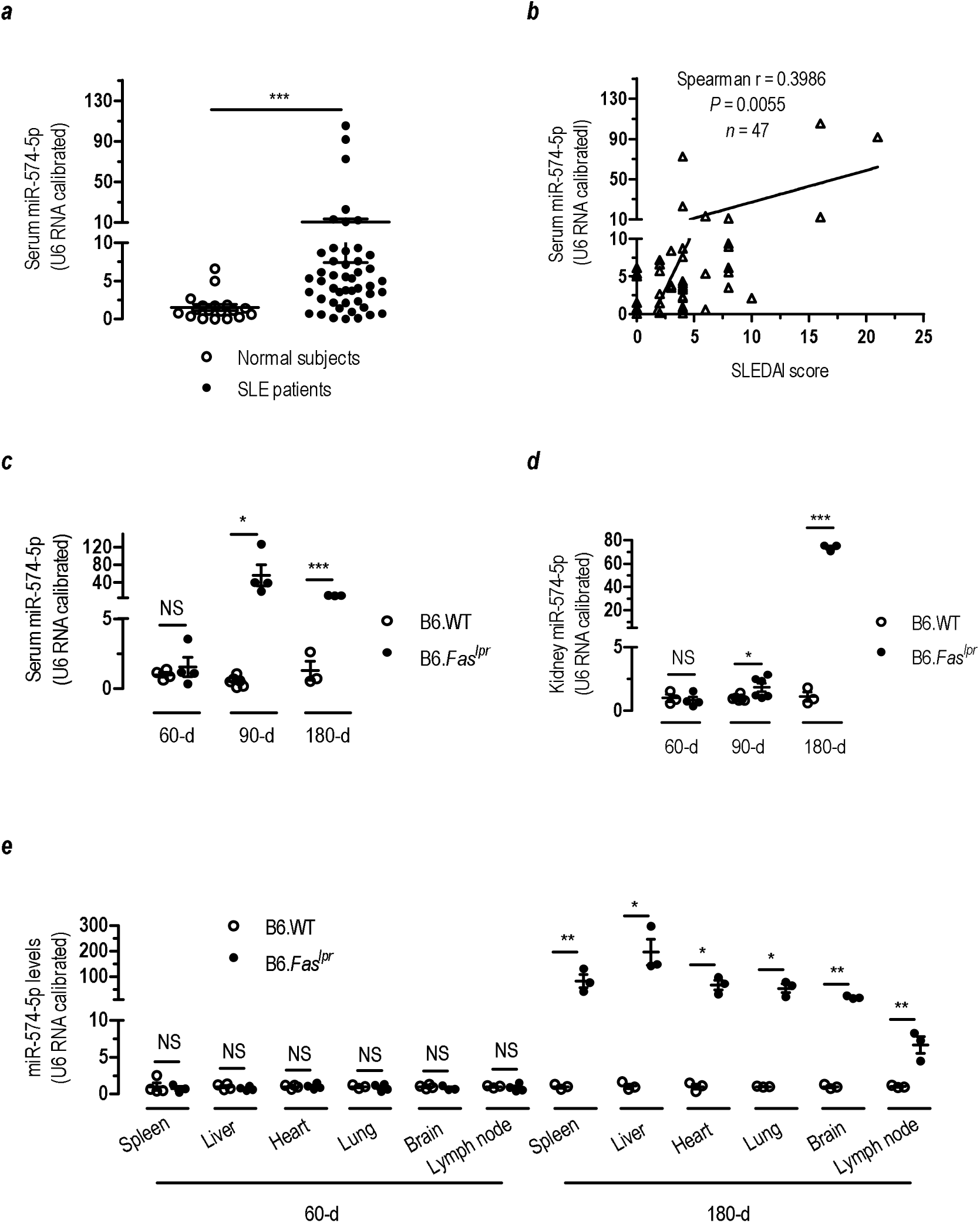
miR-574-5p is significantly up-regulated in the serum samples from human SLE patients and the serum and other tissues of lupus-prone B6.Fas^lpr^ mice. miR-574-5p levels were determined by qPCR as described in the Extended Materials and Methods in the Supplementary Information. NS, not significant. (*a*) Serum levels of miR-574-5p in SLE patients and healthy normal controls and as determined by qPCR. *** *P* < 0.001, normal controls (*n* = 18) versus SLE patients (*n* = 47), by *t*-test with Mann Whitney post-test. (*b*) Correlation between the serum miR-574-5p levels and the clinical SLE disease activity index (SLEDAI) scores in 47 SLE patients. (*c*) Serum levels of miR-574-5p in female B6.WT or B6.*Fas^lpr^* mice at ages of 60-d, 90-d and 180-d as determined by qPCR. * *P* < 0.05, *** *P* < 0.001, B6.WT versus B6.*Fas^lpr^*; *n* = 3-6. (*d*) Kidney levels of miR-574-5p in female B6.WT or B6.*Fas^lpr^* mice at ages of 60-d, 90-d and 180-d as determined by qPCR. * *P* < 0.05, *** *P* < 0.001, B6.WT versus B6.*Fas^lpr^*; *n* = 3-6.). (*e*) miR-574-5p levels in the brain, heart, liver, lung, lymph node and spleen tissues of female B6.WT or B6.*Fas^lpr^* mice at the age of 60-d and 180-d. * *P* < 0.05, ** *P* < 0.01, B6.WT versus B6.*Fas^lpr^*; *n* = 3.

### miR-574-5p binds and co-localizes with hTLR8 and mTlr7

To determine whether miR-574-5p can serve as a TLR ligand, we analyzed the ability of miR-574-5p to specifically bind to human or mouse TLRs. We first expressed histidine (His)-tagged ligand-binding domains of hTLR7/8 or mTlr7/8 in *Drosophila melanogaster* S2 cells as reported previously ^31^. The truncated-TLR proteins were then purified and incubated with either digoxin (Dig)-labeled miR-574-5p or a negative control miR-16 ^23^ to allow for miRNA-TLR interactions. Following pull-down of Dig-labeled miR-574-5p or miR-16 by anti-Dig antibodies, immunoblots with anti-His antibody showed that only the truncated-domains of hTLR8 or mTlr7 but not the truncated-domains of hTLR7 or mTlr8 were detected in the anti-miR-574-5p-Dig-antibody immuno-precipitates (Figures 2***a*** & 2***c***). In HEK-293T cells expressing the full-length hTLR7/8/9 or mTlr7/8/9, we found that only hTLR8 and mTlr7 but not the hTLR7/9 or mTlr8/9 were present in the anti-miR-574-5p-Dig-antibody co-immunoprecipitates (Figures 2***b*** & 2***d***). In contrast to miR-574-5p, there was no interaction between miR-16 and mTlr7/hTLR8. Confocal fluorescence microscopy revealed that transfected miR-574-5p was predominantly localized in the endolysosomal compartments (Figures 2***e***). Furthermore, miR-574-5p was found to co-localize with hTLR8 but not hTLR4 in the cytoplasm of HeLa cells (Figure 2***f***). Although miR-574-5p appeared to co-localize with hTLR7, the interaction was significantly weaker. In the peritoneal macrophages from the wild-type C57BL/6 (B6.WT) mice, miR-574-5p was found to co-localize with mTlr7, and this co-localization was absent in the macrophages from the *mTlr7*-deficient (B6.*Tlr7^-/-^*) mice (supplemental Figure S1). Together, these results suggest that miR-574-5p specifically interacts specifically with hTLR8 or mTlr7.

**Figure 2.**
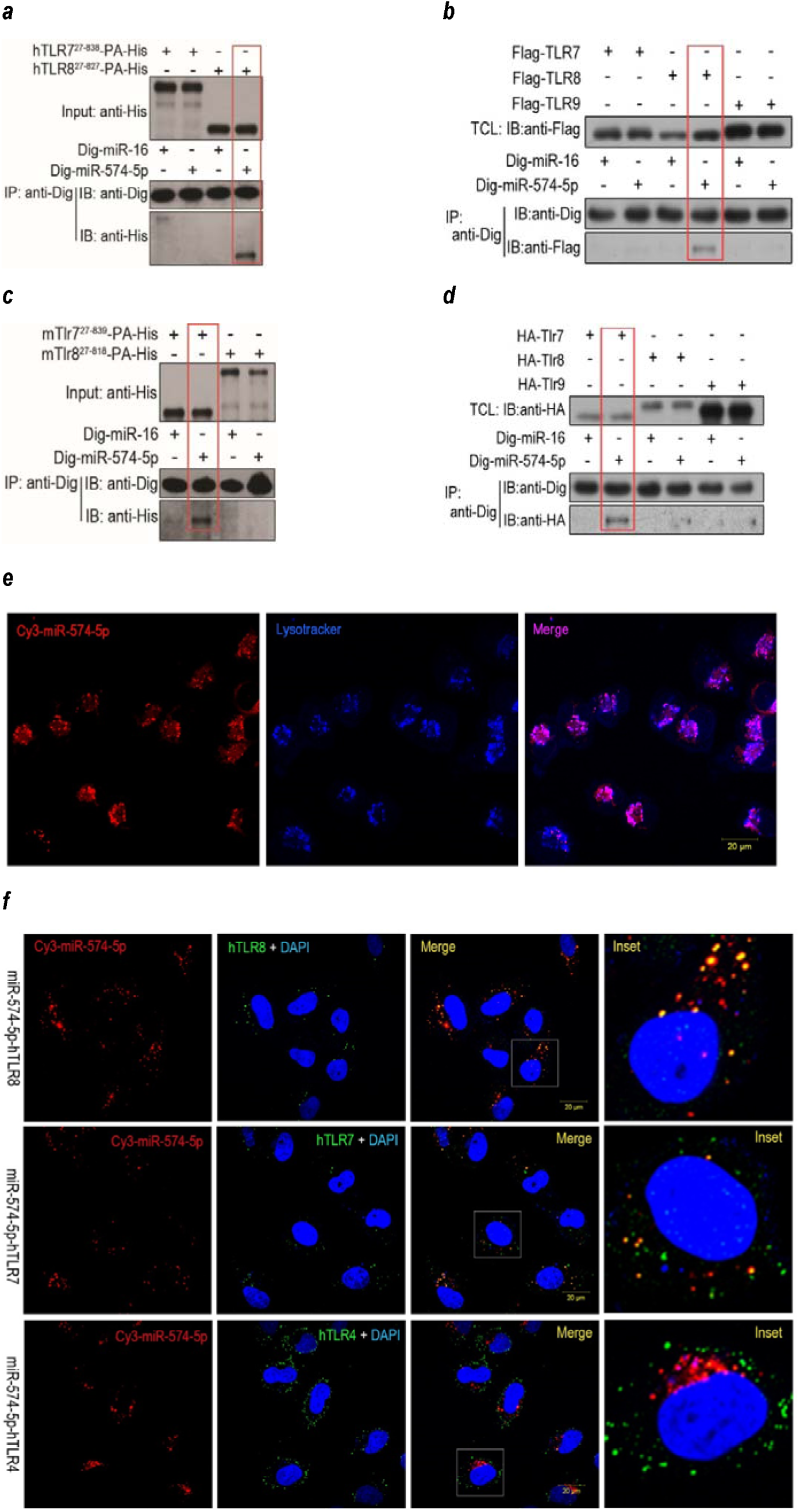
miR-574-5p binds and col-localizes with hTLR8 and mTlr7. Results were representative for 2-3 independent experiments. TCL, total cell lysate; IP, immunoprecipitation; IB, immunoblot. RNA-protein co-immunoprecipitation (co-IP) and fluorescence confocal microscopy were performed as described in the Online Method section. (***a***) Co-IP assays demonstrating the binding of Dig-miR-574-5p with a truncated hTLR8 but not a truncated hTLR7. His, histidine; PA, protein A. (***b***) Co-IP assays demonstrating the binding of Dig-miR-574-5p with the full-length hTLR8 but not the full-length hTLR7 and hTLR9. (***c***) Co-IP assays demonstrating that the binding of Dig-miR-574-5p with a truncated mTlr7 but not a truncated mTlr8. (***d***) Co-IP assays demonstrating that the binding Dig-miR-574-5p with the full-length mTlr7 but not the full-length mTlr8/9. (***e***) Localization of transfected miR-574-5p in the endolysosomal compartments. About 1×10^5^ HeLa cells were seeded onto 20 mm cell culture plates and grown to 50% confluency. Cells were treated with 3 μg/well Dotap-formulated Cy3-labelled miR-574-5p (Genscript, Nanjing, China) and incubated for 12 h in darkness. The resultant cells were washed four times with PBS and further stained with LysoTracker blue DND-22 (Cat#L7525, Invitrogen, diluted 1:1,000 in Medium) for 10 min before examination under a Zeiss LSM 780 confocal microscope. Scale bar, 20 μm. (***f***) Confocal fluorescence microscopy demonstrating co-localization of miR-574-5p with hTLR7/8 but not with hTLR4. HeLa cells were transfected with Dotap-formulated Cy3-miR-574-5p (red) stained with either anti-hTLR8 or anti-hTLR7 or anti-TLR4 (green) and DAPI (blue) and visualized under a confocal microscope. The yellow spots in the merged images represented the co-localization of miR-574-5p with the TLRs. A selected cell in the merged images were also digitally magnified (the last column). The scale bars represented 20 μm.

### Ligation of hTLR8 by miR-574-5p induces potent immune or inflammatory responses in human cells

To evaluate the functional significance of the binding of miR-574-5p to hTLR8, we first overexpressed or knocked-down miR-574-5p in human monocytic THP1 cells (Figure 3***a***). Luciferase reporter assays indicated that overexpression of miR-574-5p significantly increased whereas inhibition of miR-574-5p significantly suppressed the transcriptional activity mediated by NFκB and the interferons, with only minor exceptions (Figure 3***b***). Importantly, western blot analyses showed that overexpression of miR-574-5p markedly increased total signal transducers and activators of transcription-1 (STAT1) and its activated form, Y^701^–phosphorylated STAT1 (pSTAT1^Y701^), whereas inhibition of miR-574-5p significantly suppressed protein expression of MyD88, tumor TNF receptor-associated factor-3 (TRAF3), STAT1 and pSTAT1^Y701^ (Figure 3***c***). These results indicate a direct activation of inflammatory interferon and cytokine/chemokine signaling transducers by miR-574-5p. To examine the outcomes of miR-574-5p-mediated signaling, we analyzed the ability of miR-574-5p in stimulating the production of interferons and pro-inflammatory cytokines. Human PBMCs from healthy donors were collected and treated with Dotap-formulated miR-574-5p or infected with lentiviruses overexpressing miR-574-5p, with chemical hTLR8 agonist R848 (1 μg/ml) and miR-16 served as a positive and a negative control respectively. ELISA analyses indicated that miR-574-5p significantly stimulated the production of interferon-α and interferon-γ (IFNα/γ), tumor necrosis factor-α (TNFα) and interleukin-6 (IL6), as compared with that of human PBMC controls including cells transfected with miR-16 (Figures 3***d*** & 3***e***). Meanwhile, flow cytometry analyses showed that miR-574-5p exposure significantly increased the proportion of TNFα-positive human PBMCs, which was comparable to that induced by R848 (supplemental Figure S2***a***). To demonstrate that miR-574-5p mediated these inflammatory actions via hTLR8 but not hTLR7, we performed transfection studies with Dotap-formulated miR-574-5p in cells stably overexpressing either hTLR7 or hTLR8 (HEK-Blue-TLR7 or HEK-Blue-TLR8). Luciferase reporter assays indicated that miR-574-5p stimulates the NFκB transcriptional activity in the hTLR8-overexpressing cells but not the hTLR7-overexpressing cells (Figure 3***f***). Interestingly, the presence of uridine significantly enhanced the stimulatory effects of miR-574-5p on hTLR8, which is consistent with a recent finding from hTLR8 X-ray crystal structural studies ^31^. The specific activation of hTLR8 but not hTLR7 by miR-574-5p was further verified by a series of transfections studies in hTLR8- or hTLR7-knockdown cells (supplemental Figure S3).

**Figure 3.**
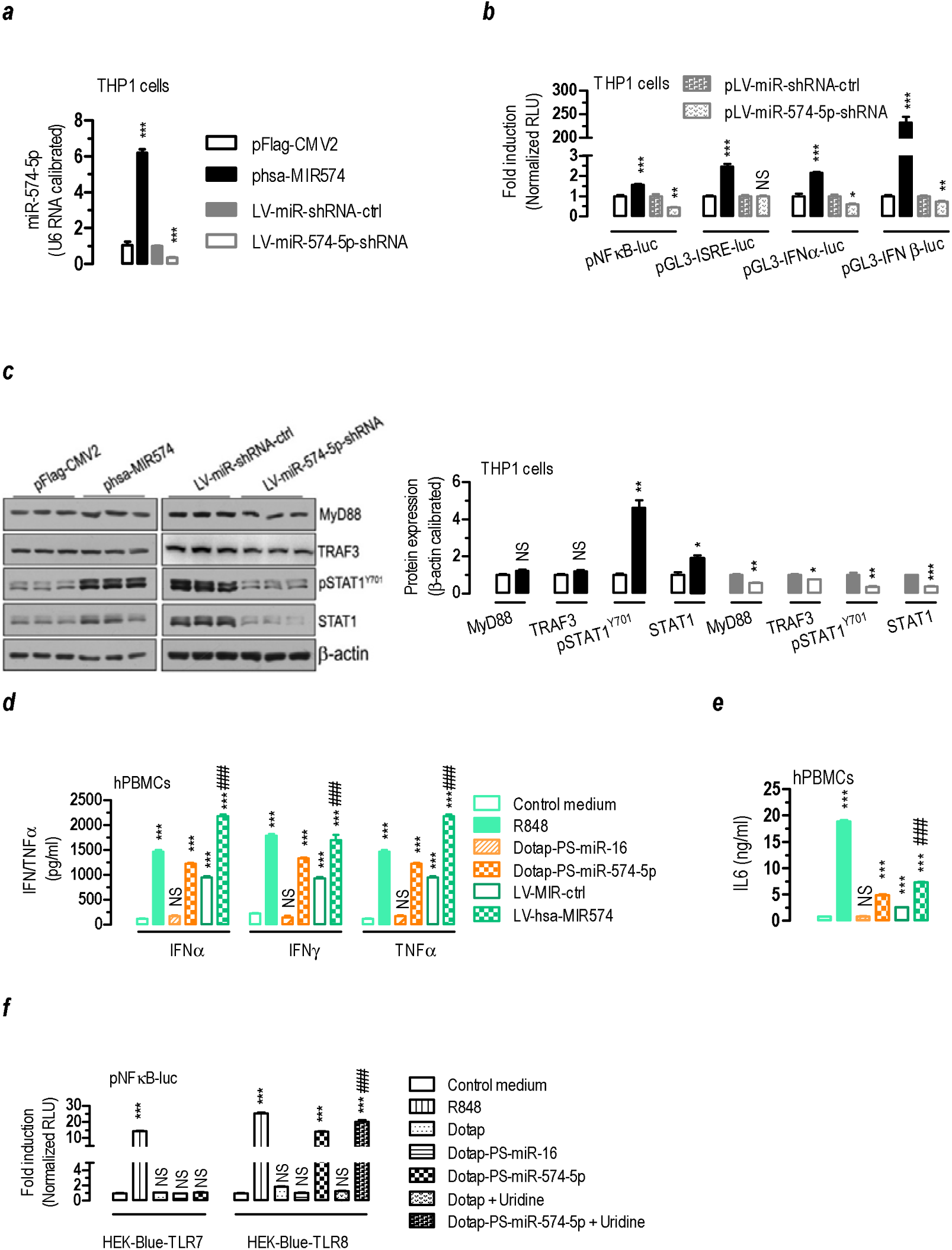
miR-574-5p overexpression stimulates whereas miR-574-5p knockdown suppresses interferon and cytokine responses in human cells. NS, not significant; * *P <* 0.05, ** *P <* 0.01, *** *P <* 0.001, compared to an appropriate group. (*a*) Verification of miR-574-5p overexpression or knockdown in THP1 cells (*n* = 3-4). (*b*) miR-574-5p overexpression stimulated whereas miR-574-5p knockdown suppressed NFκB and interferon-mediated transcriptional activities in THP1 monocytic cells. THP1 cells were co-transfected with plasmids as indicated and incubated for 24 h. Cells were harvested for the luciferase reporter assays and β-galactosidase activity assay (*n* = 3). (*c*) Effects of miR-574-5p overexpression or knockdown on the protein expression of myeloid differentiation primary response gene-88 (MyD88), TNF receptor-associated factor-3 (TRAF3), STAT1 and pSTAT1^Y701^ in THP1 cells. THP1 cells were transfected with indicated plasmids for 24 h, or infected with lentiviruses as indicated for 96 h. Cells were harvested for western blotting assays (*n* = 3). (*d*-*e*) miR-574-5p exposure potently stimulated the secretion of IFNα/γ, TNFα and IL6 in human PBMCs (*n* = 3). About 1×10^6^ human PBMCs were seeded in 6-well plate and treated with 1 μg/ml R848 or 10 μg/ml of Dotap-PS-miR-16 or Dotap-PS-miR-574-5p for 24 h. Alternatively, human PBMCs were infected with LV-MIR-ctrl or LV-hsa-MIR574 for 96 h. Subsequently, the media were harvested for ELISA analyses. ### *P <* 0.001, LV-MIR-ctrl versus LV-hsa-MIR574. (*f*) Dotap-formulated miR-574-5p transfection stimulated NFκB transcriptional activity in HEK-Blue-TLR8 cells but not in HEK-Blue-TLR7 cells. HEK-Blue-TLR7 or HEK-Blue-TLR8 cells were grown and co-transfected with a luciferase reporter plasmid (pNFκB-luc) and pSV40-β-galactosidase (3:1). 24 h after the transfection, cells were stimulated with 1 μg/ml of R848 or 10 μg/ml of Dotap-PS-miR-16 or Dotap-PS-miR-574-5p and in the absence or presence of 4 mM of uridine for 8 h. Cells were collected for luciferase activity assay (*n* = 3). ### *P <* 0.001, Dotap-PS-miR-574-5p versus Dotap-PS-miR-574-5p + uridine.

### miR-574-5p-mediated responses in mouse cells are MyD88- and mTlr7-dependent

To determine if miR-574-5p regulates immune and inflammatory responses through the TLR-MyD88 dependent signaling pathway, we further performed a series of ELISA, flow cytometry and western blot assays. In cultured *MyD88*^+/+^ RAW264.7 cells, miR-574-5p alone or miR-574-5p + uridine significantly stimulated Tnfα production, whereas in *MyD88*^-/-^ cells, the stimulatory effects were completely absent (Figure 4***a***). *In vivo*, lentivirus-mediated miR-574-5p overexpression significantly increased serum Tnfα and Il6 levels in B6.WT mice but not as much for B6.*Tlr7^-/-^* mice (Figure 4***b***). Likewise, miR-574-5p exposure significantly increased Tnfα-positive cells and Tnfα production in cultured bone marrow derived DCs (mBMDCs) prepared from the B6.WT mice but not that from the B6.*Tlr7^-/-^* mice (supplemental Figure S4). In mouse splenocytes isolated from the wildtype B6.WT mice, miR-574-5p exposure not only induced a significant upregulation of both Stat1 and pStat1^Y701^, it also appeared to stimulate the protein expression of tripartite motif containing-21/52-kDa ribonucleoprotein autoantigen Ro/SS-A (Trim21/Ro52) (Figure 4***c***). In splenocytes isolated from the B6.*Tlr7^-/-^*mice, the induction of these proteins was virtually absent. These data establish that the responses mediated by miR-574-5p are *mTlr7*- and *MyD88*-dependent in mouse cells.

**Figure 4.**
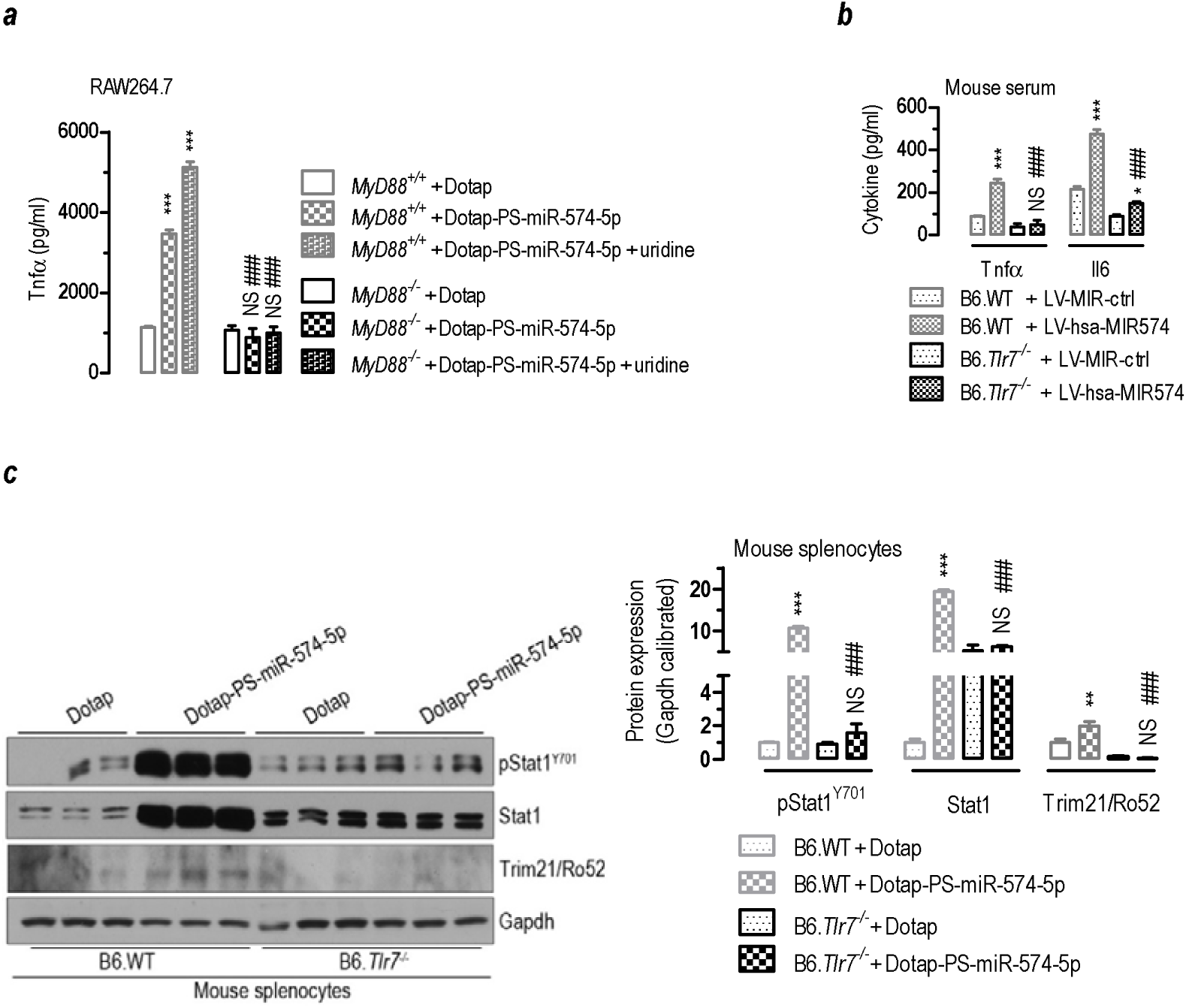
miR-574-5p-mediated immune and inflammatory responses is mTlr7- and MyD88-dependent in mouse cells. NS, not significant. (***a***) miR-574-5p or miR-574-5p + uridine exposure significantly induced Tnfα secretion in *MyD88^+/+^* but not *MyD88^-/-^* RAW264.7 macrophage cells. About 2×10^5^ *MyD88^+/+^* or *MyD88^-/-^* RAW264.7 cells were treated with Dotap only, 10 μg/ml Dotap-PS-miR-574-5p or 10 μg/ml Dotap-PS-miR-574-5p + 4 mM uridine respectively for 24 h. *** *P <* 0.001, compared to *MyD88^+/^*^+^ + Dotap or *MyD88^-/^*^-^ + Dotap; ### *P <* 0.001, *MyD88^+/^*^+^ + Dotap-PS-miR-574-5p versus *MyD88^-/^*^-^ + Dotap-PS-miR-574-5p or *MyD88^+/^*^+^ + Dotap-PS-miR-574-5p + uridine versus *MyD88^-/^*^-^ + Dotap-PS-miR-574-5p + uridine; *n* = 4. (***b***) miR-574-5p overexpression significantly increased serum Tnfα and Il6 levels in the B6.WT mice but not the B6.*Tlr7^-/-^* mice. * *P <* 0.05, *** *P <* 0.001, compared to B6.WT + LV-MIR-ctrl or B6.*Tlr7^-/-^* + LV-MIR-ctrl; ### *P <* 0.001, B6.WT + LV-hsa-MIR574 versus B6.*Tlr7^-/-^* + LV-hsa-MIR574; *n* = 4-5. (***c***) miR-574-5p exposure significantly increased Stat1, pStat1^Y701^ and Trim21/Ro52 in splenocytes prepared from the B6.WT mice but not the B6.*Tlr7^-/-^* mice. ** *P <* 0.01, *** *P <* 0.001, compared to B6.WT + Dotap or B6.*Tlr7^-/-^* + Dotap; ### *P <* 0.001, B6.WT + Dotap-PS-miR-574-5p versus B6.*Tlr7^-/-^* + Dotap-PS-miR-574-5p; *n* = 4-5.

### miR-574-5p plays a pivotal role in both innate and adaptive immunity

To systematically evaluate the immune regulatory roles of miR-574-5p, we used lentivirus-mediated RNA interference to knockdown the expression of miR-574-5p in HeLa cells, which express substantial levels of miR-574-5p (supplemental Figure S5***a***) and reported to express a high level of hTLR8^32^. Microarray analyses showed that upon miR-574-5p knockdown, 151 genes (Data File S1) were significantly up-regulated whereas 661 genes significantly down-regulated by at least 2-fold (supplemental Figure S5***b***). The trends of the expression of 12 differentially expressed mRNAs as revealed by the array analyses were further verified by qPCR analyses of mRNAs and/or Western blot analyses (supplemental Figures S5***c*** & S5***d***). Gene ontology and clustering analyses revealed that the differentially-expressed genes were highly enriched for the immune system and its related signal transduction pathways. Of particular interests, however, were the 661 genes that were significantly down-regulated in miR-574-5p knockdown HeLa cells, as they apparently were not caused directly by miR-574-5p through the canonical miRNA-mediated posttranscriptional silencing. Indeed, pathway enrichment analyses showed that the down-regulated genes were highly enriched for antigen processing and presentation, cytokine and interferon signaling and endosome/vacuolar pathway (supplemental Figure S5***e***), *etc*. Moreover, the significantly down-regulated genes were also highly enriched in pathways associated with infection, graft-versus-host disease, rheumatoid arthritis and SLE and Type I diabetes mellitus (supplemental Figure S5***f***).

Next, we isolated mouse splenocytes from the B6.WT mice and transfected the isolated immune cells with Dotap-formulated miR-574-5p. RNA profiling by RNA-Seq analyses showed that in comparison with the control Dotap-treated cells, miR-574-5p transfection for 24 h resulted in significant up-regulation of 277 genes (Data File S2) and down-regulation of 80 genes by at least 2-fold in the miR-574-5p-transfected splenocytes (supplemental Figures S6***a****-**c**,* supplemental Table S2-S4). qPCR analyses of 28 selected mRNAs verified the validity of the RNA-Seq assays (supplemental Figure S6***d***). Gene ontology and pathway enrichment analyses showed that the 277 significantly up-regulated genes in miR-574-5p-transfected cells were highly enriched and clustered in pathways associated with interferon and cytokine signaling, antiviral mechanisms, translesion synthesis and autoimmunity (Figure 5): 1) Of the 277 significantly upregulated genes, about 82 (29.6%) can be attributed directly to the immune system. 2) miR-574-5p exposure potently activated the interferon/cytokine signaling and antigen processing and presentation. In particular, miR-574-5p exposure potently stimulated a large set of interferon α/β/γ-stimulated genes (ISGs) that is typical of viral infection, which included 2’-5’ oligoadenylate synthetase proteins (*Oas*), lymphocyte antigen 6 complex proteins (*Ly6*), MX dynamin-like GTPase 1/2 (*Mx1/2*), histocompatibility 2 proteins, 15-kDa interferon-stimulated protein (*Isg15*), interferon regulatory factor 1/7 (*Irf1/7*), tripartite motif containing-21/25 (*Trim21/25*) and promyelocytic leukemia protein (*Pml*); 3) miR-574-5p exposure not only stimulated the mRNA expression of ssRNA-sensing *mTlr7* itself, it also significantly stimulated the expression of dsRNA-sensing *Tlr3,* protein kinase RNA-activated (*Pkr*), the cytosolic RNA-sensing receptors DEXD/H-box helicase-58 (*Ddx58*), DEXH-box helicase-58 (*Dhx58*), interferon induced with helicase C domain-1 (*Ifih1*) and the DNA-sensing Z-DNA binding protein-1 (Z*bp1*) and multiple interferon-inducible p200 family proteins (*Ifi203/204/205/207/209, Mnda, Mndal*); 4) miR-574-5p exposure significantly stimulated the expression of Ifnγ-inducible nuclear autoantigens including component of Sp100-rs (*Csprs*), *Sp100*, *Sp110*, *Sp140*, and *Trim21/Ro52*; 5) miR-574-5p exposure stimulated the expression of non-canonical poly-A RNA polymerase D7 (*Papd7*), replication factor C subunit-3 (*Rfc3*). Papd7 and Rfc3 play important roles in translesion synthesis and DNA repair bypass. qPCR analyses of 14 miR-574-5p-induced genes in splenocytes again confirmed that the induction was largely mTlr7-dependent (supplemental Figure S6***e***). These results highlight the systemic and comprehensive regulatory roles of miR-574-5p-mTlr7 signaling in innate and adaptive immune responses and autoimmunity.

**Figure 5.**
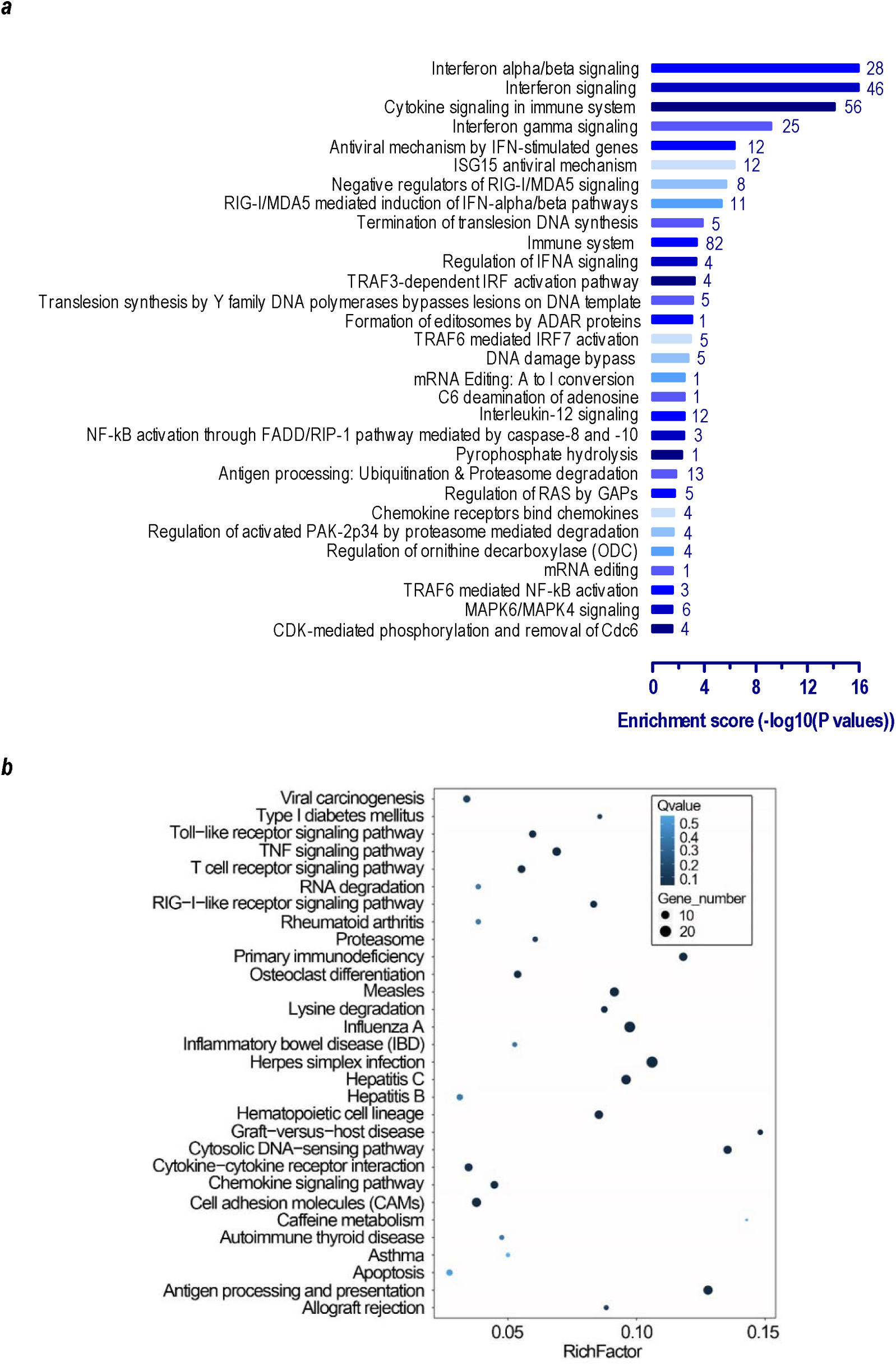
miR-574-5p exposure in mouse splenocytes induces significant alterations in the expression of genes associated with immune and inflammatory responses. Splenocytes from 8-wk old female B6.WT mice (*n* = 3) were divided into two equivalent parts, one for 10 μg/ml of Dotap-PS-miR-574-5p and one for Dotap only transfection for 24 h. Gene expression was profiled by RNA-Seq as described on Supplementary Methods. (*a*) 30 top signaling pathways that were significantly enriched for genes that were up-regulated by at least 2-fold as a consequence of miR-574-5p exposure (Dotap-PS-miR-574-5p versus Dotap-only, *n* = 3) as analyzed by the Reactome pathway analysis. The digital number next to each bar represented the count of the significantly-up-regulated genes enriched in the pathway. (*b*) 30 top biological or disease pathways that were significantly enriched for genes that were up-regulated by at least 2-fold as a consequence of miR-574-5p exposure (Dotap-PS-miR-574-5p-treated versus Dotap-only treated, *n* = 3) as analyzed by the KEGG pathway analysis. The rich factor is the ratio between the number of the significantly-up-regulated genes mapped to the pathway and the total number of genes in the pathway and the Q-values were the *p*-values corrected with the false discovery rates.

### miR-574-5p exposure leads to the activation of human and mouse B cells

B cells play a crucial role in SLE development. Thus, we examined if miR-574-5p exposure will result in the activation of the B cells. In human PBMCs, both R848 and miR-574-5p not only significantly increased the number of CD19^+^ B cells but also the number of activated CD19^+^CD69^+^ B cells (Figure 6***a***), suggesting that miR-574-5p not only promotes the proliferation or preferential survival but also the activation of B cells. qPCR analyses of B-cell activation markers including B-cell activating factor receptor (BAFF-R), B-lymphocyte antigen CD19 and transmembrane activator and CAML interactor (TACI) confirmed the potential effects of miR-574-5p on B-cell activation (Figure S2***b*)**. In negatively selected human B cells, both R848 and miR-574-5p potently stimulated the activation of hPBMC-derived B cells but did not increase the number of CD19^+^ B cells (Figure 6***b***), suggesting the effect on hPBMC-derived B cell activation might be indirect. In mice, the splenic B cells derived from the B6.WT mice but not that from the B6.*Tlr7^-/-^* mice were significantly activated by both R848 and miR-574-5p, as indicated by the alterations in B220^+^CD69^+^ cells (Figure 6***c***). Together these results suggest that miR-574-5p-induced B cells activation probably is hTLR8-dependent in human and mTlr7-dependent in mice.

**Figure 6.**
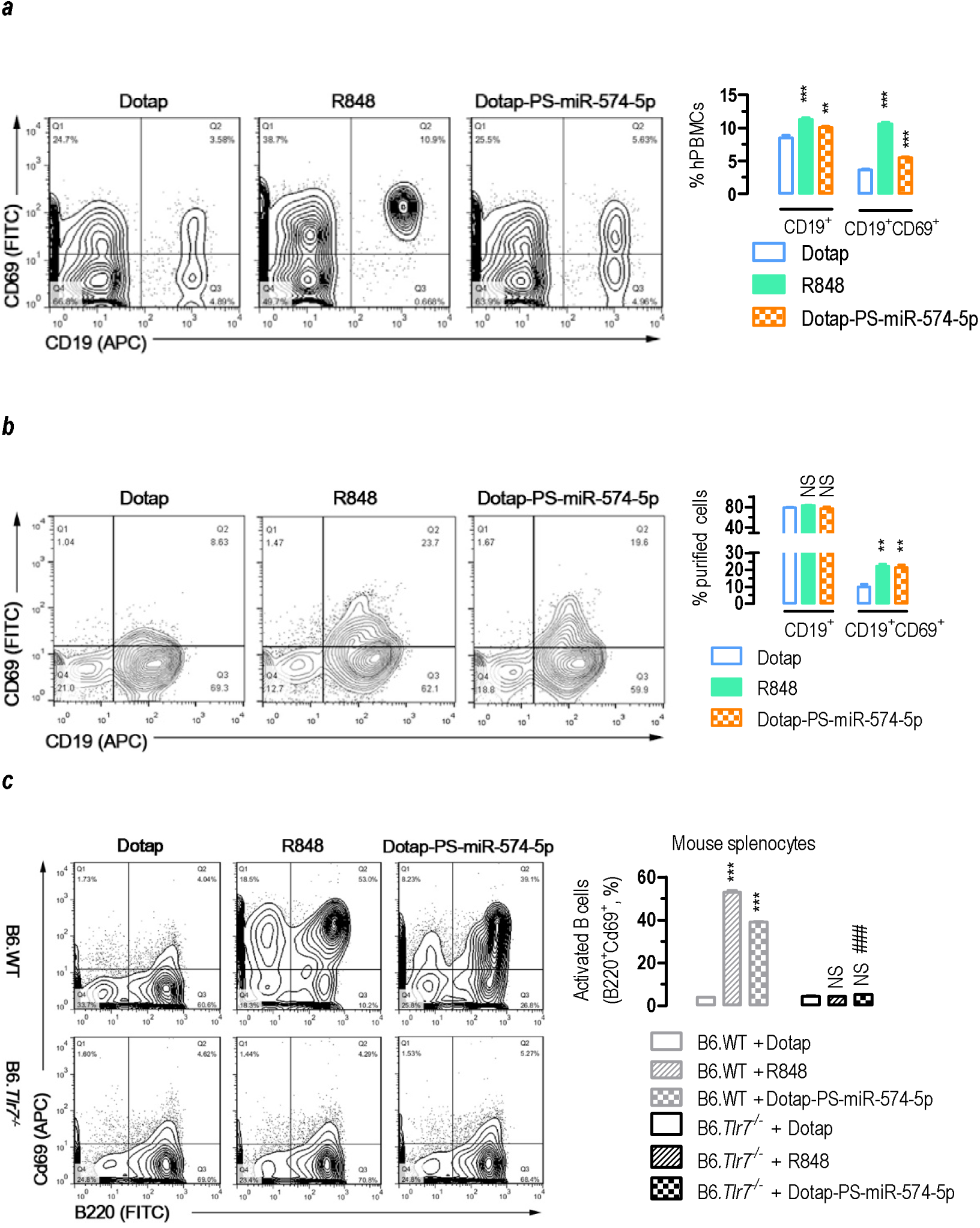
miR-574-5p exposure leads to potent activation of human and mouse B lymphocytes. (***a***) miR-574-5p exposure in human PBMCs significantly increased both total B cells (CD19^+^) and activated B cells (CD19^+^CD69^+^) (*n* = 3). About 1×10^6^ human PBMCs were seeded in 6-well plate and treated with 1 μg/ml R848 or 10 μg/ml of Dotap-PS-miR-574-5p for 24 h. Subsequently, the cells were harvested for flow cytometry. ** *P <* 0.01, *** *P <* 0.001, compared with Dotap only. (***b***) miR-574-5p exposure potently stimulated the activation of human PBMC-derived B cells (*n* = 4-5). B cells were purified from human PBMCs of three normal healthy subjects and treated as human PBMCs as described above. NS, not significant; ** *P <* 0.01, compared with Dotap only. (***c***) miR-574-5p exposure significantly increased splenic B cell activation in the B6.WT mice but not the B6.*Tlr7^-/-^* mice. About 1×10^5^ splenocytes were seeded in 6-well plates and treated with 10 μg/ml of Dotap-PS-miR-574-5p or 1 μg/ml of R848 for 24 h. B cell activation was analyzed by flow cytometry gated on B220^+^Cd69^+^. *** *P <* 0.001, compared to B6.WT + Dotap or B6.*Tlr7^-/-^* + Dotap; ### *P <* 0.001, B6.WT + Dotap-PS-miR-574-5p versus B6.*Tlr7^-/-^* + Dotap-PS-miR-574-5p; *n* = 4.

### Induction of miR-574-5p expression in vivo is associated with increased systemic inflammation and the production of autoantigens and autoantibodies in transgenic mice

To determine the effects of *in vivo* expression of miR-574-5p, we created lines of transgenic mice carrying a human *MIR574* transgene under the control of the Doxycycline-controlled Tet inducible systems (supplemental Figures S7***a***). Remarkably, induction of miR-574-5p expression (supplemental Figure S7***b***) greatly increased the levels of anti-dsDNA and antinuclear (ANA) (Figure 7***a-b***) but not anti-RNA antibodies (supplemental Figures S7***c***) in the serum in transgenic mice following 6-wk of Dox induction but not the controls. In transgenic mice following 10 weeks of Dox induction, the levels of serum Ifnα and Tnfα in Dox-induced transgenic mice were slightly increased, although the differences were not significant. However, the levels of serum Il6 and autoantigen Sp100 did increase significantly (supplemental Figures S7***d-g*)**. Meanwhile, induction of transgenic expression of miR-574-5p also caused significant renal deposit of complement C3 and IgG (Figure 7***c***) and milder levels of renal damage (supplemental Figure S7***h***) and elevated mRNA expression of *Il6, Il12, Irf7* and *Tnf*α in the spleen (supplemental Figure S7***i***). Together these results clearly link elevated expression of miR-574-5p with ANA and anti-dsDNA autoantibody production and systemic inflammation.

**Figure 7.**
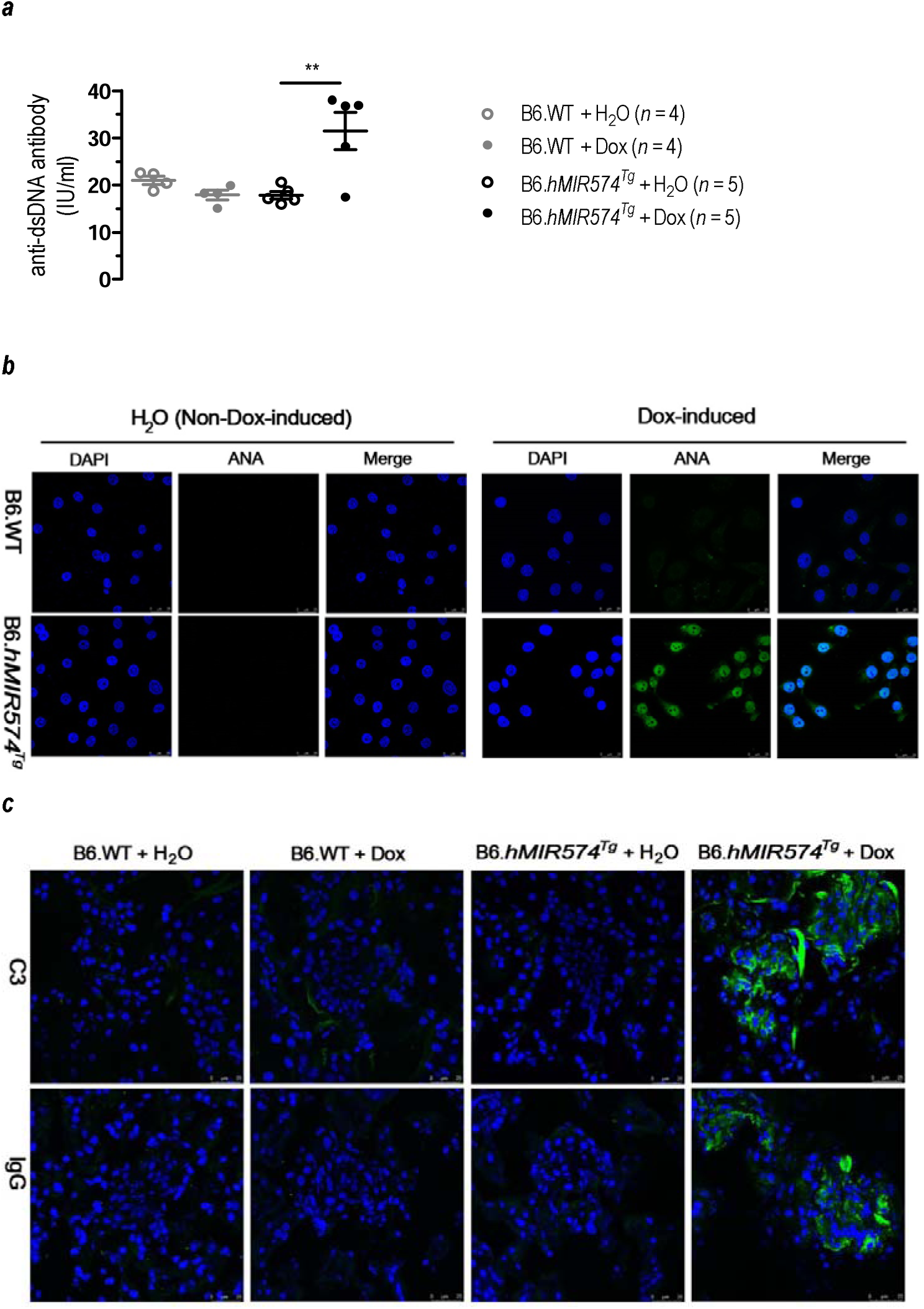
Transgenic overexpression of miR-574-5p stimulates the secretion of antinuclear antibody (ANA) and anti-dsDNA and increases the renal deposit of complement C3 and IgG. Mice carrying a Dox-inducible human *MIR574* transgene and its controls were prepared as described. Transgene expression was induced by providing mice with drinking water containing 2 mg/ml Dox starting from the age of 8-10 wk. (*a*) Stimulated secretion of serum ANA in the transgenic mice following 6-wk Dox-induction but not in the un-induced. The indirect immunofluorescence ANA tests were performed with 1:5 diluted serum samples from the transgenic mice or its controls using the HEp2 slides and FITC-tagged antibodies as instructed. Scale bar, 25 μm. Results were representative of at least three mice per group. (*b*) Increased serum levels of anti-dsDNA antibody in the transgenic mice following 6-wk Dox-induction but not in the un-induced. ELISA assays were performed as described. ** *P <* 0.01, B6. B6.*hMIR574^Tg^* + H_2_O versus B6.*hMIR574^Tg^* + Dox. (*c*) Enhanced renal deposit of complement C3 and IgG in the transgenic mice following 10-wk of Dox-induction but not in the un-induced. Renal tissues were prepared and stained with fluorescent antibodies against C3 and IgG. Results were representative of at least three mice per group.

### Depleting miR-574-5p greatly ameliorates Fas-deficiency induced SLE and lupus nephritis in the lupus-prone mice

To determine the effects of miR-574-5p silencing on the development of SLE, we injected 8-wk old female B6.*Fas^lpr^* mice with lentiviruses carrying shRNAs against miR-574-5p (LV-miR-574-5p-shRNA) or the control lentiviruses (LV-miR-shRNA-ctrl) once every two weeks at the dosage of 2×10^6^ transducing units/mouse for six times. Two weeks after the final virus injection (20-wk old), a significant reduction of miR-574-5p was observed in both the kidney and liver tissues and the serum (supplemental Figure S8***a-b***). As a consequence of miR-574-5p-knockdown, *Fas*-deficiency induced upregulation of both Stat1 and pStat1^Y701^ were largely ameliorated in the splenocytes, although the difference was not significant between the control lentivirus treated B6.*Fas^lpr^* mice and the LV-miR-574-5p-shRNA treated B6.*Fas^lpr^* mice for Stat1 (Figure 8***a***). Interestingly, activated B cells (B220^+^Gl7^+^ or B220^+^Cd69^+^, Figure 8***b***) were significantly reduced whereas T follicular helper (T_fh_ cells, Cxcr5^hi^Pd1^hi^Cd4^+^, Figure 8***c***) was slightly but not significantly reduced in splenocytes isolated from the miR-574-5p-knockdown B6.*Fas^lpr^* mice. Immunohistochemical analyses also showed that spontaneous germinal center (Spt-GC) formation (IgD^-^Pna^+^, Figure 8***d***) was significantly ameliorated in the miR-574-5p-knockdown B6.*Fas^lpr^*mice.

**Figure 8.**
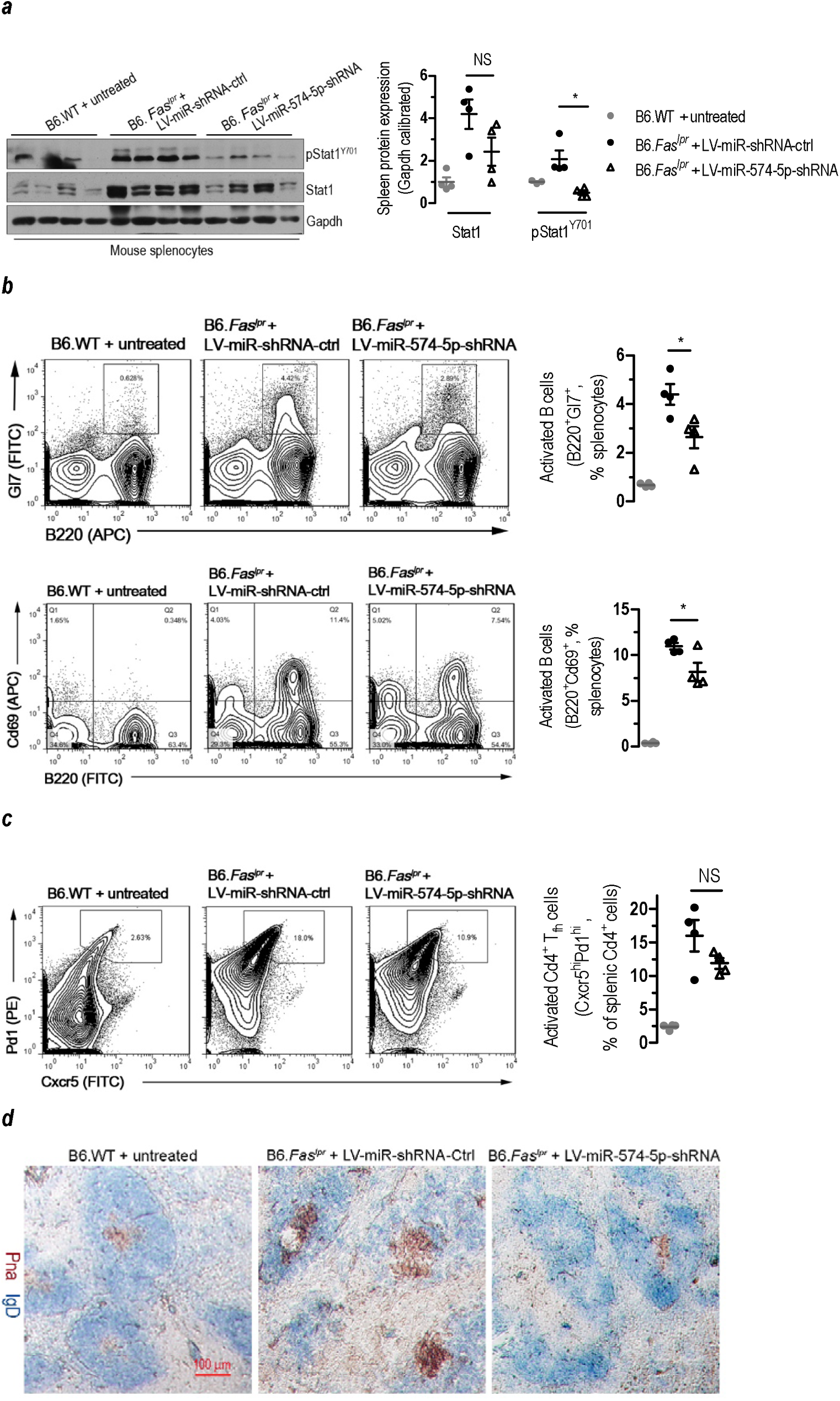
In vivo silencing of miR-574-5p in the lupus-prone mice causes significant changes in Stat1 signaling and activation of B lymphocytes. *In vivo* silencing of miR-574-5p was achieved by treatment with lentiviruses carrying shRNA against miR-574-5p as described. NS, not significant; * *P <* 0.05, B6.*Fas^lpr^*+ LV-miR-shRNA-ctrl versus B6.*Fas^lpr^* + LV-miR-574-5p-shRNA. (*a*) *S*ilencing of miR-574-5p significantly attenuated Stat1 signaling (*n* = 4). (*b*) Silencing of miR-574-5p significantly attenuated lupus-associated activation of B cells (*n* = 4). B cell activation was analyzed by flow cytometry gated on B220^+^Gl7^+^ or B220^+^Cd69^+^. (*c*) *S*ilencing of miR-574-5p slightly but not significantly attenuated lupus-associated activation of T_fh_ cells (Cxcr5^hi^Pd1^hi^ Cd4^+^, *n* = 4). Cd4^+^ splenocytes were isolated by FACS. T_fh_ cell activation was analyzed by flow cytometry gated on Cxcr5^hi^Pd1^hi^. (*d*) Typical results showing reduced formation of splenic Spt-GC following *in vivo* silencing of miR-574-5p (*n* = 2-3). Spleen tissues were dissected and stained with antibodies against IgD and peanut agglutinin (Pna) as described (Soni *et al.*, 2014). Brown staining indicated germinal center B cells (IgD^-^Pna^+^) while blue staining indicated follicle B cells (IgD^+^Pna^-^). Scale bar, 100 μm.

In 20-wk old miR-574-5p-knockdown B6.*Fas^lpr^* mice, *Fas* deficiency-associated splenomegaly appeared to be significantly ameliorated (Figure 9***a***). Despite that the serum level of Ifnα was only slightly and not significantly reduced (Figure 9***b***), the miR-574-5p-knockdown B6.*Fas^lpr^* mice had significantly lower levels of Ifnβ/γ than that of the control-treated lupus-prone mice (Figure 9***c***-***d***). In addition, with the silencing of miR-574-5p, autoantigens Trim21/Ro52 and Sp100 (Figure 9***e***-***f***), and cytokines Tnfα and Il6, blood urea nitrogen and proteinuria all appeared to be significantly reduced (supplemental Figures S8***c***-***f***). Consistently, the serum levels of antinuclear antibody (supplemental Figure S9***a***) and anti-dsDNA antibody, IgG, IgG1, IgM and IgG2b all appeared to be reduced in the miR-574-5p-knockdown B6.*Fas^lpr^* mice, although the differences were not statistically significant for IgG2b (Figure 9***g***-***k***). Immunohistochemical analysis and periodic acid-Schiff staining demonstrated that following the silencing of miR-574-5p, renal fibrotic lesions, IgG deposition (Figure 9***l***) and the ratio of mesangial to glomerular area (supplemental Figure S9***b***) were greatly ameliorated in both the renal cortex and the medulla of miR-574-5p-silenced B6.*Fas^lpr^* mice (supplemental Figure S9***c***). In the renal cortex, renal medulla and liver tissues of miR-574-5p-silenced B6.*Fas^lpr^* mice, the expression of cluster of differentiation-68 (Cd68) was also markedly reduced (supplemental Figures S9***d****-**e***), suggesting a significant suppression of monocyte/macrophage infiltration. Collectively, these results demonstrate that a generalized knockdown of miR-574-5p is protective against the autoimmune responses and the development of *Fas*-deficiency-induced SLE and lupus nephritis in the lupus-prone mice.

**Figure 9.**
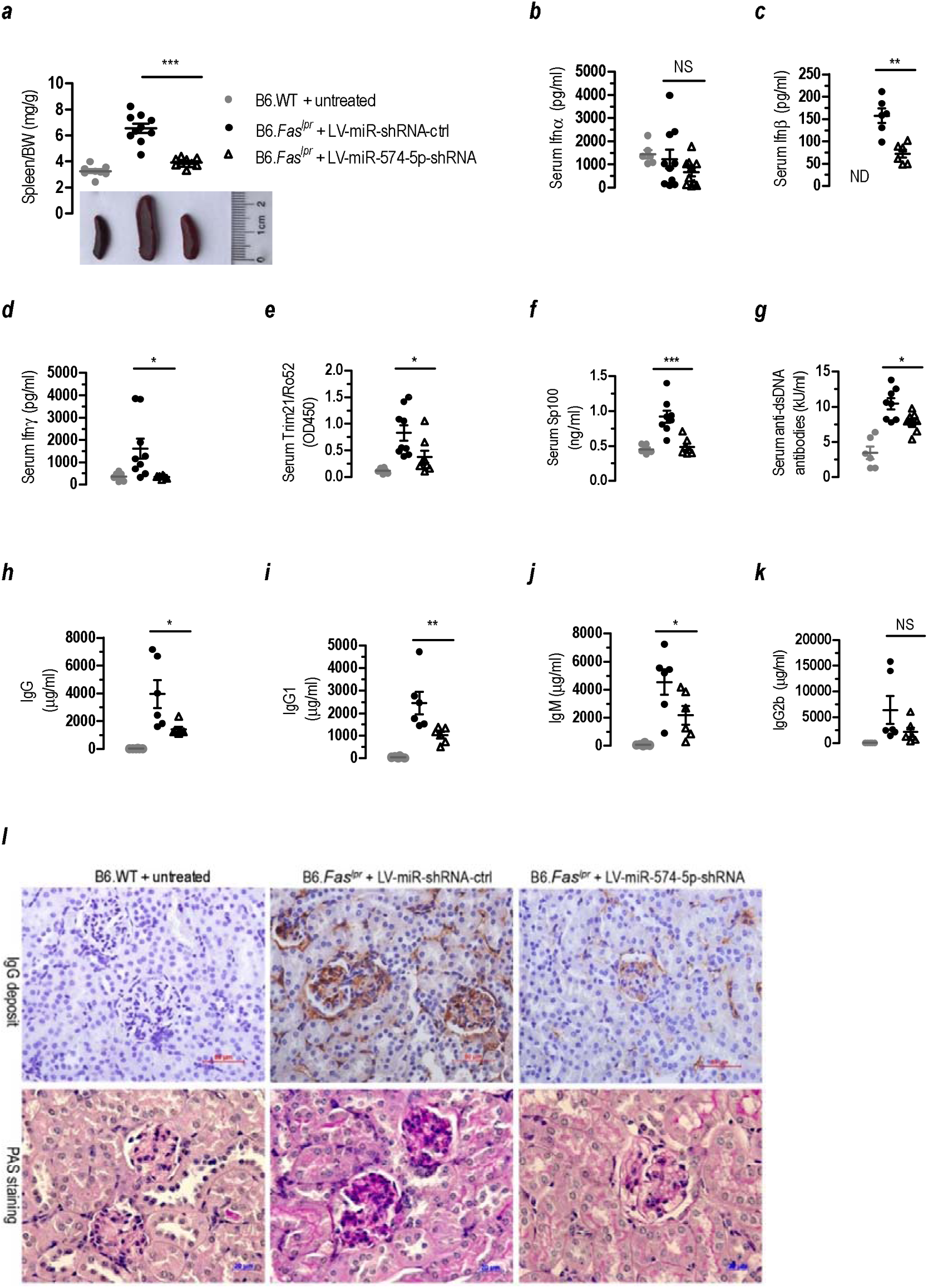
Silencing of miR-574-5p significantly ameliorates SLE and lupus nephritis in the B6.Fas^lpr^ mice. NS, not significant; ND, not detectable; * *P <* 0.05, ** *P <* 0.01, *** *P <* 0.001, LV-miR-shRNA-ctrl versus LV-miR-574-5p-shRNA. *In vivo* silencing of miR-574-5p was achieved by treatment with lentiviruses carrying shRNA against miR-574-5p as described. (***a***) Silencing of miR-574-5p significantly ameliorated lupus-associated splenomegaly (*n* = 8-10). (***b***-***d***) Silencing of miR-574-5p in the lupus-prone B6.*Fas^lpr^* mice significantly reduced the serum levels of Ifnβ/γ but not Ifnα (*n* = 6-9). (***e***-***f***) Silencing of miR-574-5p in the lupus-prone B6.*Fas^lpr^* mice greatly reduced the serum levels of Trim21/Ro52 and Sp100 (*n* = 6-8). (***g***), Silencing of miR-574-5p in the lupus-prone B6.*Fas^lpr^* mice led to reduced levels of serum anti-dsDNA antibody (*n* = 6-8). (***h-k***) Silencing of miR-574-5p significantly reduced the serum levels of IgG, IgG1 and IgM but not IgG2b (*n* = 6). (***l***) Silencing of miR-574-5p significantly ameliorated IgG deposit as determined by immunohistochemistry and abnormal renal structure and morphology as determined by the periodic acid-Schiff (PAS) staining. Results were typical for at least three mice/group. Scale bars, 50 (IgG) or 20 (PAS) μm.

## DISCUSSION

Our results prove a critical role for TLR8-mediated miR-574-5p activity in lupus pathogenesis or progression. Firstly, in human patients and lupus-prone mice we find that miR-574-5p levels in circulating plasma and a number of tissues are increased and positively associate with lupus severity. Secondly, we demonstrate that miR-574-5p binds to hTL8 and mTlr7 specifically to induce a series of inflammatory and immune responses leading to increased secretion of cytokines and interferons, the activation of B lymphocytes. Finally, we demonstrate the ability of miR-574-5p to induce lupus-like features in mice, including systemic inflammation, increased production of autoantigen Sp100 and autoantibodies ANA and anti-dsDNA, the renal deposit of complement C3 and IgG, whereas its down regulation significantly alleviates most symptoms associated with lupus and lupus nephritis. These findings reveal a novel axis of autoimmune signaling.

miR-574-5p is known to be one of the RNA cargos in exosomes or small extracellular vesicles ^24–26^. Many studies have shown that miR-574-5p is often dysregulated in conditions such as microbial ^33^ and viral infection ^34^, drug or toxin-induced injury ^35,36^ and cancers ^37–40^. For instance, miR-574-5p and miR-146a were among the most significantly up-regulated miRNAs in mouse peritoneal macrophages following vesicular stomatitis virus infection ^34^. In human dermal fibroblasts, miR-574-5p is one of the most significantly up-regulated miRNAs following UV exposure ^41^. Moreover, γ-irradiation or LPS exposure dose- and time-dependently upregulated miR-574-5p expression in dermal fibroblast and the serum of rhesus macaques ^42^. In sentinel lymph nodes from melanoma patients, it was recently shown that miR-574-5p was the most significantly upregulated among the differentially expressed miRNAs that were associated with dysfunctional immune responses ^43^. Together these observations raised the possibility for miR-574-5p to serve as a danger signal or alarmin that alerts the host either pathogen invasion or cell/tissue damage in order to mount appropriate defense mechanisms.

Although hTLR7 and hTLR8 are two highly homologous proteins that are capable of signaling in response to viral ssRNAs, functionally hTLR7 appears to differ significantly from hTLR8. In addition, m*Tlr7* is believed to be the murine ortholog of hTLR8 but not hTLR7 ^44^. The functional difference between mTlr7 and mTlr8 is even bigger than that of hTLR7 versus hTLR8. In fact, mTlr8 is not responsive to most ligands for hTLR7/8 and mTlr7. Surprisingly, mTlr8 deficiency rather than its activation leads to autoimmunity in mice ^45^. In human, however, gain-of-function variants in hTLR8 lead to a novel childhood-onset in born errors of immunity with lymphoproliferation, neutropenia, infectious susceptibility, B-and T-cell defects, and in some cases, bone marrow failure ^27^. When a human hTLR8 transgene was introduced into mice, the high copy number chimeras were unable to pass germline and developed severe inflammation targeting the pancreas, salivary glands, and joints, suggesting that RNA recognition by hTLR8 can lead to autoimmune inflammation ^29^. Mice with relatively lower levels of transgene expression survived but had increased susceptibility to collagen-induced arthritis. These observations highlight the critical role of hTLR8 in autoimmunity.

A number of miRNAs have been shown to interact with hTLR7/8 ^46^. Most of the TLR-interacting miRNAs that have been identified so far, however, were reported to act either on hTLR8 or its murine ortholog mTlr7 ^23,44,47–50^, and so far there is very limited cases reported for hTLR7. As stated above, in Salvi et al.’s report ^26^, the direct experiment linking miR-574-5p with hTLR7 was a NF-κB luciferase reporter assay in TLR-transfected HEK293 cells and as a matter of fact, both hTLR8 and hTLR7 were activated by miR-574-5p in these assays. In more recent studies, the same group identified a group of SARS-CoV-2-associated molecular patterns (SAMPs) that have the potential to serve as ligands for human TLR7/8 ^51^. However, in the HEK293 NF-κB luciferase reporter assays, the activation of hTLR7 by SAMPs is far less significant than that of hTLR8. In neutrophiles, the activation of hTLR8 but not hTLR7 is observed ^52^. Further, in an acute graft-versus-host disease study, serum miR-29a has been shown to be able to activate mTlr7 in mouse DCs and hTLR8 in human DCs ^44^. In our current study, the inability of miR-574-5p to bind to and activate hTLR7 was demonstrated not only by the TLR-transfected HEK293 cell NFκB-luciferase reporter assays (Figure 3***f*** & supplemental Figure S3) but also by the more direct antibody pull-down (Figure 2***a***-***d***), which were in conflict with that of similar experiments by Salvi et al.^26^. Our results, however, are in accordance with most demonstrations of the preferential binding of the small RNA ligands with hTLR8 and mTlr7 but not hTLR7, which included RNA40 ^6^, miR-21 and miR-29 ^23^, miR-29a ^44^ and HERVK(HML-2) RNA ^53^. The binding specificity of miR-574-5p with hTLR8 or mTlr7 probably is dependent on the higher order structure TLRs and the miRNAs. However, the exact nature of the RNA motifs recognized by TLR7/8 *in vivo* remains to be fully investigated.

An important and unexpected result is our demonstration that miR-574-5p exposure induces the overproduction of autoantigens through activating TLR7/8 signaling. TLR stimulation has previously been shown to induce the upregulation of *TRIM* genes ^54^, especially the induction of *TRIM21/Ro52* by TLR3 ^55^. Surprisingly, miR-574-5p exposure significantly upregulated the mRNA expression of Ifnγ-inducible *Trim21/Ro52* and *Sp100* as well as other nuclear autoantigens *Sp110, Sp140* and *Csprs* (supplemental Figure S6 and Data File S2). The induction of autoantigen by miR-574-5p was further confirmed by the demonstration that *in vivo* induction of miR-574-5p transgenic expression induced a significant increase in serum autoantigen Sp100 (supplemental Figure S7***g***), whereas miR-574-5p knockdown in the lupus-prone mice resulted in a significant reduction of both Trim21/Ro52 and Sp100 (Figure 9***e***-***f***). MiR-574-5p-mediated up-regulation of autoantigens on one hand might feed-back to TLR signaling to further amplify the immune and inflammatory signaling. On the other hand, the complexes these autoantigens can form with cytosolic ssRNAs might further facilitate the production of autoantibodies. To the best of our knowledge, this is not only the first demonstration that miR-574-5p is associated with the production of autoantigens, it is also the first demonstration that Tlr7/hTLR8 is associated with the production of autoantigens.

Endosomal TLRs are known to play a pivotal role in the development of autoinflammation and autoimmunity ^11,22,56^. Recognition of self-nuclei acids by the endosomal TLRs on DCs or the B cells, in particular, is thought to be an important step in the pathogenesis of autoimmune diseases, which is believed to initiate the production of interferons and anti-nuclear antibodies. Previous studies on TLR-driven autoimmune responses were almost exclusively focused on hTLR7 and hTLR9 ^57,58^, whereas little is known about the roles of hTLR8. As stated, hTLR8 differs significantly from hTLR7 in many ways including the patterns of expression, ligand recognition and the function. It has been demonstrated previously that although R837 or R848 both were able to induce proliferation and activation of human B cells. R848, however, was almost 100-fold more potent than R837 ^59^. Furthermore, R848 but not R837 induced IgM synthesis and class switching. These results appear to highlight the critical importance of hTLR8 in B cell activation. In this study, we showed that miR-574-5p exposure led to a significant increase in both CD19^+^ cells and CD+CD69^+^ cells in human PBMCs (Figure 6***a***), suggesting a potent stimulation and activation on human B cells. Noteworthy, however, is that in negatively enriched B cells, CD+CD69^+^ cells but not CD19^+^ cells were increased (Figure 6***b***), probably suggesting the lack of direct stimulation. MiR-574-5p-mediated B cells activation thus probably needs the assistance of monocytes or DCs with high hTLR8 expression in human PBMCs or T_fh_ cells in the germinal center, which remains to be further investigated. In mice, mTlr7, the murine ortholog of hTLR8, appears to be critical for Spt-GC formation, T_fh_ and B cell development ^60^. Further, IFNγ-IFNγ receptor and STAT1 signaling in B cells has been shown to be central for Spt-GC formation and B cell responses ^61,62^. Consistent with these observations, we demonstrate herein that miR-574-5p exposure leads to both enhanced production of IFNγ and activation of STAT1 signaling in both human and mouse cells (Figures 3-4). Furthermore, miR-574-5p exposure leads to the activation of mouse splenic B cells, whereas this stimulation is absent in mice deficient for mTlr7. Since hTLR8 probably is not expressed in the B cells, the activation of B cell by R848 or miR-574-5p should be indirect.

Consistent with previous studies of SLE and rheumatoid arthritis human patients ^30,63,64^, we showed that miR-574-5p was not only significantly elevated in the serum of SLE patients but also in serum and a number of tissues in the lupus-prone B6.*Fas^lpr^* mice (Figure 1). Furthermore, the up-regulation in miR-574-5p expression appears to be positively correlated with the development of lupus or lupus nephritis in both humans and mice. In cell cultures miR-574-5p was shown to be able to induce a series of MyD88-dependent immune and inflammatory responses, which include the overproduction of cytokines and interferons, the activation of STAT1 signaling and B lymphocytes, and the production of autoantigens. In the lupus-prone mice we showed that *in vivo* silencing of miR-574-5p led to a significant and systemic alleviation in almost all the lupus-associated parameters assayed in the lupus-prone mice (Figures 8-9 and supplemental Figures S8-9), which includes the amelioration in the lupus-associated aberrant Ifnγ and Stat1 signaling, Spt-GC formation and the activation of B cells, proteinuria, blood nitrogen, the *Fas* deficiency-associated splenomegaly, renal fibrosis, renal IgG deposit and leukocyte infiltration in the liver and the kidney and the overproduction of serum Ifnβ/γ, Tnfα and Il6, IgG and IgM, antinuclear and anti-dsDNA autoantibodies and Trim21/Ro52 and Sp100. Most striking, however, is our demonstration utilizing the *in vivo* transgenic mouse models. In these transgenic mice, we demonstrate for the first time that *in vivo* transgenic overexpression of miR-574-5p leads to symptoms resembling the development of human SLE. Transgenic expression miR-574-5p for 6 or 10-wk evokes the production of ANA and anti-dsDNA but not anti-RNA antibodies, the renal deposit of IgG and C3 (Figure 7) and the increased renal tissues damage (supplemental Figure S7). Consistent with the results from cultured splenocytes, *in vivo* overexpression of miR-574-5p expression leads to increased serum level of Il6 (supplemental Figure S7***f***) and increased mRNA expression of *Tnfα*, *Il6*, *Il12* and *Irf7* (supplemental Figure S7***i***). Particularly noteworthy is that as a consequence of miR-574-5p overexpression in the transgenic mice, serum autoantigen Sp100 is significantly increased (supplemental Figure S7***g***). These *in vivo* results clearly exemplify how dysregulation of miR-574-5p can trigger abnormal miR-574-5p-hTLR8 signaling leading to systemic inflammation and autoimmunity. They represent the systemic and comprehensive demonstration that TLR-triggering miRNAs are indeed involved in the onset or development of complex autoimmune diseases such as SLE.

In summary, our results strongly support a major role for the aberrant over-activation of miR-574-5p-hTLR8/mTlr7 in driving severe inflammation and autoimmunity. Conversely, the suppression of this signaling is effective in mitigating inflammation and autoimmune responses. These findings provide substantial insights into the pivotal role of miR-574-5p as a prominent immune and inflammation regulator. This positions miR-574-5p as a valuable target for the development of novel immunotherapeutic drugs for the prevention or treatment of diseases including autoimmune disorders, viral infection and cancers.

## MATERIALS AND METHODS

### Construction of plasmids and lentiviruses

Full-length hTLR7/8/9-overexpressing plasmids pFlag-TLR7, pFlag-TLR8 and pFlag-TLR9 were generous gifts of Prof. Jiahuai Han, Xiamen University.

Plasmid vectors expressing small hairpin RNA (shRNA) against hTLR7 (NM_016562.3) and *hTLR8* (NM_016610.3) were constructed by inserting chemically-synthesized double-strand DNA fragments containing *hTLR7* and *hTLR8*-targeting shRNA sequences as listed in supplemental Table S5 into plasmid pLentiLox3.7 at the HapI and XhoI sites, generating plasmids pLV-sh-TLR7-1, pLV-sh-TLR7-2, pLV-sh-TLR8-1, pLV-sh-TLR8-2, pLV-sh-TLR8-3 and pLV-sh-ctrl. Plasmid vectors expressing small hairpin RNA (shRNA) against miR-574-5p were constructed by inserting chemically-synthesized double-strand DNA fragments containing miR-574-5p-targeting shRNA sequences as listed in supplemental Table S5 into plasmid pLentiLox3.7 at the HapI and XhoI sites, generating plasmids pLV-miR-574-5p-shRNA and pLV-miR-shRNA-ctrl. The inserted DNA fragments were verified by DNA sequencing.

To create a plasmid overexpressing the ligand-binding domain (not including the transmembrane and the cytosolic domains) of hTLR8, a truncated hTLR8 (amino acids 27-827, TLR8^27–827^) was amplified from pFlag-TLR8 whereas a protein A (PA) cDNA fragment was amplified from *Staphylococcus aureus subsp. Aureus* (strain USA300_TCH1516, Cat#29213, ATCC, Manassas, VA, USA) using primers listed in supplemental Table S6. TLR^827–827^ and PA cDNA fragments were fused by overlapping PCR and the resultant TLR8^27–827^-PA fragment was inserted into the *Drosophila* Expression System vector pMT-BIP-V5-His (Cat#V413020, Biofeng, Shanghai, China), using *BglII* and *EcoRI* restriction sites to generate PA- and His-double-tagged TLR8^27–827^ overexpressing plasmid pTLR8^27–827^-PA-His. In a similar way, other plasmids containing the truncated hTLR7 or mTlr7/8, which included pTLR7^27–838^-PA-His, pTlr7^27–839^-PA-His and pTlr8^27–818^-PA-His, were prepared using primers listed in supplemental Table S6.

Plasmids overexpressing human miR-574-5p (phsa-MIR574) and lentiviruses carrying shRNAs for miR-574-5p (LV-miR-574-5p-shRNA) or a negative control shRNA (LV-miR-shRNA-ctrl) were as described previously ^39^. To construct a lentiviral vector overexpressing miR-574-5p, a 345-bp human *MIR574* gene DNA fragment was PCR amplified from human genomic DNA with the primers listed in supplemental Table S6. The PCR-amplified fragment was inserted to a lentiviral vector pLV-EF1a-MCS-IRES-Puro (pLV-MIR-ctrl) to generate pLV-hsa-MIR574. Viral vector pLV-hsa-MIR574 or pLV-MIR-ctrl as well as three lentivirus packaging plasmids (pMDL, pVSVG and pREV) were co-transfected into HEK-293T cells. pLV-EF1a-MCS-IRES-Puro, pMDL, pVSVG and pREV were kind gifts from Prof. Jiahuai Han. Media containing lentiviruses (LV-hsa-MIR574 and LV-MIR-ctrl) were collected every 24 h for 3 times and the lentiviruses were purified by ultra-speed centrifugation.

### Animals and treatments

Mice were housed in the specific pathogen-free conditions in the Xiamen University Laboratory Animal Center or Dalian Medical University Animal Facility, with a 12 h-12 h light-dark cycle and regular chow and water provided *at libitum*. All experimental procedures involving animals were performed in accordance with animal protocols approved by the Institutional Animal Use and Care Committee of Xiamen University and Dalian Medical University. Lupus-prone mice (B6.MRL-Fas^lpr^/J, B6.*Fas^lpr^*) were obtained from Nanjing University, Nanjing, Jiangshu, China. *mTlr7* deficient mice (B6.129S1-Tlr7^tm1Flv^/J, B6.*Tlr7^-/-^*) were purchased from the Jackson Lab (Cat#008380, Jackson Lab, Bar Harbor, Maine, USA). Normal wild-type C57BL/6 mice (B6.WT) were used as controls for both B6.*Tlr7^-/-^* and B6.*Fas^lpr^*.

To knockdown miR-574-5p *in vivo*, 8-wk old female B6.*Fas^lpr^*mice were administered intravenously with 2×10^6^ TU/mouse of LV-miR-574-5p-shRNA or its control lentiviruses LV-miR-shRNA-ctrl once every two weeks up to a total of 1.2×10^7^ TU/mouse. 2 weeks after the final injection, serum, urine, kidney and liver tissue samples were collected from 20-wk old mice for analyses. Urine was collected by bladder message. Blood samples were collected by sinus puncture.

For transient miR-574-5p overexpression *in vivo*, 10-wk old male B6.*Tlr7^-/-^ or* B6.WT mice were infected with lentiviruses overexpressing miR-574-5p or the control viruses at a dosage of 1 ×10^7^ transforming unit (TU)/mouse once by intravenous injection. 72 h after the lentiviral administration, mice were sacrificed and serum and tissue samples were collected for analyses.

Doxycycline (Dox)-inducible human MIR574 transgenic mice were prepared by the Cyagen Biosciences (Guangzhou) Inc. Briefly, a human pre-MIR574 DNA fragment was cloned into a pTRE-tight vector such that the expression of the transgene is under the control of the Dox-controlled Tet systems, as illustrated in the plasmid map (supplemental Figure S7***a***). Pronuclear microinjection was performed and the founder transgenic lines were in the C57BL/6 background. The founder mice were crossed with normal C57BL/6 mice to generate heterozygous F1 mice and the controls (B6.WT and B6.*hMiR574^Tg^*) for the experiments. Genotyping of transgenic mice was performed by PCR analyses of the tail genomic DNA using transgene-specific primers (5’-CCCCTGAACCTGAAACATAAA and 5’-AACAGCTAAAGTG-CGAAAGCG) and the WT control primers (5’-CTATCAGGGATACTCCTCTTTGCC and 5’-GATACAGGAATGACAAGCTCATGGT).

8-10 wk transgenic mice and the control littermates were provided with drinking water in the absence or presence of doxycycline (Dox, final concentration 2 mg/ml) (Beyotime, ST039B, China). 6 or 10 wk after the Dox treatment, blood samples were collected. The serum levels of antinuclear antibody (ANA), anti-dsDNA and anti-RNA were examined. The mice were sacrificed 10 weeks after Dox induction and serum, kidney and spleen tissues were collected. Mice were housed in a specific pathogen-free facilities at Xiamen University and Dalian medical University with free access to food and water.

### Cell culture, transfection and treatment

Human monocytic THP1, cervical cancer HeLa and embryonic kidney (HEK-293T) cells were purchased from American Type Culture Collection (Manassas, VA, USA) and maintained as instructed. HEK-293T cells overexpressing hTLR7/8 (HEK-Blue-TLR7/8) were purchased from InvivoGen (San Diego, California, USA) and cultured in DMEM supplemented with 10% (vol/vol) FBS, Normocin (50 μg/mL), Blasticidin (10 μg/mL), and Zeocin (100 μg/mL) (InvivoGen, San Diego, CA, USA). The wild-type (*MyD88^+/+^*) and MyD88-deficient (*MyD88^-/-^*) RAW264.7 cells were kind gifts provided by Jiahuai Han of Xiamen University. RAW264.7 cells were cultured in DMEM (Gibco) medium containing 10% (vol/vol) FBS and 100 IU/ml penicillin and 100 μg/ml streptomycin.

Human PBMCs were isolated from the whole blood of healthy human donors by centrifugation through a Ficoll-hypaque gradient centrifugation. These cells were cultured in RPMI1640 (Gibco, Grand Island, NY, USA) supplemented with 10% (vol/vol) fetal bovine serum (FBS).

Mouse peritoneal cells were harvested by peritoneal lavage in 7-10 wk old female B6.WT mice with 8-10 mL of ice-cold PBS. Peritoneal macrophages were centrifuged at 350×g for 5 min and the resulting peritoneal macrophages were replated at 5 × 10^5^ cells/ml in DMEM supplemented with 10% FBS, 2 mM L-glutamine and 100 IU/ml penicillin and 100 μg/ml streptomycin (all purchased from Sangon Biotech, Shanghai, China) and cultured overnight prior to transfection or stimulation.

Splenic cell suspensions were prepared from the spleen tissues dissected from 7-10 wk old female B6.WT and B6.*Tlr7^-/-^* mice. Red blood cells were eliminated by osmotic lysis using red blood cell lysis buffer (Cat#00-4300-54, eBioscience, San Diego, CA, USA) for 5 min. Splenocytes were obtained by centrifugation at 350×g for 5 min and the resulting cells were plated at 5 × 10^5^ cells/ml in DMEM supplemented with 10% FBS, 2 mM L-glutamine and 100 IU/ml penicillin and 100 μg/ml streptomycin. Mouse splenocytes were cultured overnight prior to transfection or stimulation.

Lipofectamine-3000 reagent (Invitrogen, Carlsbad, CA, USA) was used for plasmid DNA transfections. Phosphorothioated or digoxin labeled miRNAs were chemically synthesized by Invitrogen, Guangzhou, Guangdong, China or Genscript, Nanjing, Jiangsu China as listed on supplemental Table S7. Cationic lipid N-[1-(2,3-Dioleoyloxy)propyl]-N,N,N-trimethylammonium methyl-sulfate (Dotap, Liposomal Transfection Reagent, Cat#1202375, Roche, Nonnenwald, Penzberg, Germany) was used for the conjugation of phosphorothioated miRNAs and Dotap-formulated miRNAs were used for *in vitro* delivery of miRNAs, according to the manufacturer’s protocols. Unless otherwise indicated, Dotap-formulated and phosphorothioated miRNAs were used for cell transfection at the concentration of 10 μg/ml. hTLR7/8 dual-agonist resiquimod (R848, Cat#tlrl-r848, Invitrogen) was used in cell studies at the working concentration of 1 μg/ml. Lipopolysaccharide (LPS) was purchased from Sigma-Aldrich (Cat#L2630, Sigma-Aldrich, St Louis, MO, USA) and used in cell cultures at the concentration of 100 ng/ml.

### Luciferase reporter assays

A luciferase reporter for NFκB activity (pNFκB-luc) was a kind gift from Prof. Jiahuai Han, Xiamen University whereas reporters for interferon activity (pISRE-luc, pGL3-IFNα-luc and pGL3-IFNβ-luc) were provided by Prof. Rongtuan Lin, McGill University, Montreal, Canada. Luciferase reporter activities in cells co-transfected with any of the luciferase reporter were determined using a luciferase reporter gene assay system (Cat#E1601, Promega, Madison, WI, USA) as instructed. For all luciferase assays, β-galactosidase activities were determined to calibrate for the transfection efficiency. The calibrated value for a proper control was used to normalize all other values to obtain the normalized relative luciferase units (RLU).

### miRNA-TLR protein co-immunoprecipitation (co-IP) assays

To express and purify a truncated hTLR8 protein (the ligand-binding domain), *Drosophila melanogaster* Schneider 2 (S2) cells (Cat#R690-07, Invitrogen, Carlsbad, CA, USA) were transfected with pTLR8^27–827^-PA-His, in the presence of plasmids pCoHygro (Cat#K4130-01 Biofeng, Shanghai, China). 48 h after the transfection, cells were selected with Sf-900 II SFM medium (Cat#10902-096, Gibco, Grand Island, NY, USA) containing 300 μg/mL hygromycin (Cat#10687-010 Invitrogen, Carlsbad, CA, USA) to obtain cells stably-overexpressing TLR8^27–827^-PA-His. Approximately 2.5×10^8^ stable cells were induced with CuSO_4_ at the concentration of 5 mM for 72 h and 50 ml of the media were collected as described ^65^. Protein TLR8^27–827^-PA-His was purified with IgG beads (Cat#17-0969-01, GE Healthcare, Connecticut, CT, USA) and resuspended in 200 μl of NT2 buffer ^66^ and used for miRNA-protein co-IP. In a similar way, truncated hTLR7^27–838^-PA-His, mTlr7^27–839^-PA-His and Tlr8^27–818^-PA-His were overexpressed and purified.

For *in vitro* co-IP of miRNA with the truncated PA- and His-double tagged-TLRs, firstly 1 μg of 5’-Dig-labeled miR-16 or miR-574-5p (Dig-miR-16 and Dig-miR-574-5p, supplemental Table S7) were mixed with 100 μl above purified hTLR8^27–827^-PA-His and 900 μl of in NET2 buffer ^66^ to allow potential miR-TLR interactions at 4 °C for 3 h. Following this, 1 μg of anti-Dig antibody was added to the mixtures and incubated at 4 °C overnight to allow anti-Dig-(Dig-miR) interactions. Protein A/G beads (Genscript, Nanjing, Jiangsu, China) were subsequently added to the mixtures and incubated for another 5 h. Beads were washed 6 times with the NT2 buffer. The eluted samples were blotted by anti-His and anti-Dig antibodies (supplemental Table S8). Similar co-IPs were performed with truncated proteins hTLR7^27–838^-PA-His, mTlr7^27–839^-PA-His and Tlr8^27–818^-PA-His.

For co-IP of with the full-length TLRs, 5×10^6^ HEK-293T cells were seeded and transfected with approximately 5 μg Flag-tagged hTLR7/8/9 overexpressing plasmids or HA-tagged mTlr7/8/9 overexpressing plasmids. 24 h after the transfection, cells were harvested and lysed by ultrasonification in 300 μl of the polysome lysis buffer on ice, as previously described ^67^. Lysates were centrifuged at 14,000 ×g for 15 min. 100 μl resultant supernatant were incubated with 1 μg of 5’-Dig-labled and phosphorothioated miRNAs in NET2 buffer for 3 h at 4 °C. Subsequently, anti-Dig antibody were added to the supernatants and the mixtures were incubated at 4 °C overnight. Pull-down was achieved by incubating the mixtures with protein A/G beads in a final volume of 1 ml of NET2 buffer for 5 h at 4 °C. Beads were centrifuged and washed six times with NT2 buffer. The eluted samples were analyzed by immunoblots using anti-Dig, anti-Flag or anti-HA antibodies (supplemental Table S8).

### Localization of miR-574-5p in the endolysosomes and co-localization of miR-574-5p with TLRs in HeLa cells or mouse macrophages as demonstrated by fluorescence microscopic analyses

For the localization of miR-574-5p in the endolysosomes, HeLa cells were seeded onto cell culture plates (1×10^5^ /well) and grown to 50% confluence. Then cells were transfected with Dotap-conjugated and Cy3-labelled miR-574-5p (3 μg/well, supplemental Table S7) and incubated for 12 h. The resultant cells were washed four times with PBS and further stained with LysoTracker blue DND-22 (Cat#L7525, Invitrogen, diluted 1:1000 in PBS) for 2 h before examination under a confocal microscope.

For the co-localization of miR-574-5p with human TLR4/7/8 in the cytoplasm, HeLa cells were seeded into 6-well plates (1×10^5^/ well) with coverslips inside the wells and grown to 50% confluence. Then cells were transfected with Dotap-formulated and Cy3-labelled miR-574-5p (3 μg/well) and incubated for 12 h in darkness. The resultant cells were washed twice with PBS and fixed with 4% paraformaldehyde at room temperature for 10_min. After washing twice with PBS, the cells were incubated with 0.2% Triton X-100 and 0.2% albumin bovine V (BSA) in PBS at 4°C for 10 min to permeate the cell membrane. Then, a blocking solution (0.02% Triton X-100 and 5% BSA in PBS) was added and the plates were incubated at room temperature for 30 min. The cells were respectively incubated with antibodies against human TLR4 or hTLR7 or hTLR8 (supplemental Table S8) in the blocking solution at 4 °C for 16 h. The cells were washed three times (5 min/each) with washing buffer, 0.02% Triton X-100 and 1.5% BSA in PBS and incubated with secondary anti-rabbit or anti-goat antibody labeled with Alexa Fluor 488 (supplemental Table S8) in blocking solution at 37 °C for 2_h in darkness. The secondary antibodies were washed off with above-mentioned washing buffer for 15 min and finally slides were mounted with DAPI (Vector Laboratories, Burlingame, USA). The stained cells were examined under a Zeiss LSM 780 fluorescence microscope (Carl Zeiss MicroImaging, Jena, Germany).

For the co-localization of miR-574-5p with mTlr7, mouse peritoneal macrophages were isolated from 8-weeks-old female B6.WT and B6.*Tlr7^-/-^* mice by peritoneal lavage with 8-10 mL of ice-cold PBS and centrifugation at 400×g for 5 min. Macrophages were plated in 6-well plates (1×10^5^/ well) with coverslips inside the wells. In a similar way stated above for human TLRs, macrophage cells were transfected with Dotap-formulated and Cy3-labelled miR-574-5p and stained with anti-mTlr7 and secondary antibodies (supplemental Table S8).

### Preparations of mouse BMDCs

Mouse mBMDCs were prepared by methods described previously ^68^, with minor modifications. Briefly, bone marrow cells obtained from mouse tibias and femurs from 7-10 wk old female B6.WT and B6.*Tlr7^-/-^* mice were passed through a nylon mesh to remove debris, and approximately 3×10^6^ cells were placed in 6-well plates containing 3 ml dendritic cell medium (RPMI1640 supplemented with 10% FBS, 10 ng/ml GM-CSF (Cat#14-8331-62, eBioscience, San Diego, CA, USA) and 10 ng/ml IL4 (Cat#14-8041-62, eBioscience, San Diego, CA, USA). On day-4 and day-7, 50% of the medium was replaced with fresh media. On day-7 or day-8, the loosely adherent clusters were dislodged and harvested gently for subsequent experiments.

### Flow cytometry

For cell-surface staining, human PBMCs, mBMDCs, or mouse splenocytes were seeded at the density of 1×10^6^/well on 6-well plate and treated with 10 μg/ml of Dotap-conjugated miR-574-5p or 1 μg/ml of R848 or 100 ng/ml of LPS. 24 h after the stimulation, cells were washed, resuspended in cold FACS buffer (PBS containing 0.1% sodium azide (Cat#S2002, Sigma-Aldrich (China), Shanghai, China) and 2% FBS), and then incubated for 10 min at 4 °C. Subsequently, they were stained with antibodies as listed on Table S8 for 30 min. For intracellular staining, cells were washed with cold FACS buffer and fixed in Fixation buffer (Cat#420801, BioLegend, San Diego, CA, USA) at 4 °C for 1 h. Cells were then washed with Permeabilization buffer (Cat#421002, BioLegend, San Diego, CA, USA) and stained with anti-TNFα antibody in Permeabilization buffer at 4 °C for 4 h. The stained cells were washed with FACS buffer and analyzed by flow cytometry (LSRFortessa, Becton Dickinson, San Jose, CA, USA).

### Human B cell isolation and flow cytometry analyses

Blood samples were collected and pooled from 3 healthy normal volunteers (20 ml/each). Human PBMCs were harvested after Ficoll separation. B cells were then isolated using a human B cells enrichment kit (Stemcell Technologies, Cat#19054). Briefly, human PBMCs were suspended at a 1×10^7^/ml concentration in a medium buffer containing 1 M EDTA and 2% fetal bovine serum. Cells then were mixed with a biotin labeled negative selection antibodies to remove non-B cells by the magnet beads. Isolated cells were cultured for 4 hours and then treated with R848 (1 μg/ml) or transfected with DOTAP or DOTAP formulated miR-574-5p (10 μg/ml). Cells were collected after 24 h and incubated with an APC labeled anti-CD19-APC and a FITC labeled anti-CD69. CD19 and CD69 single or double positive cells were counted by a flow cytometry.

### Urinary protein and blood urea nitrogen analyses

To assess renal function, urinary protein (Cat#C035-2, Jiancheng Bioengineering Institute, Nanjing, Jiangsu, China) concentrations and blood urea nitrogen (BUN, Cat#C013-2) were measured using commercial kits from Jiancheng Bioengineering Institute (Nanjing, Jiangshu, China), following the manufacturer’s instructions.

### Histochemistry and immunohistochemistry analyses

For histological and immunohistological analyses, mice were sacrificed at the age of 20 weeks and kidney and liver tissues were rapidly dissected and fixed in 4% paraformaldehyde (v/v) for 24 h before being embedded in paraffin. Kidney tissue sections at 5-μm thickness were stained with periodic acid-Schiff (PAS, Cat#YM0715LA13, Yuanye BioTechnology, Shanghai, China) and examined under a light microscope. For IgG deposits in the kidney, renal sections were incubated with peroxidase-conjugated anti-mouse IgG. Staining was visualized using the chromogenic substrate 3-3’ diaminobenzidine (Cat#KIT-0017, Maixin Biotech, Fuzhou, Fujian, China). In addition, kidney and liver tissue sections were de-paraffinized, rehydrated and subjected to antigen retrieval in citrate buffer solution (pH 6.0). Sections were then incubated with mouse anti-Cd68 antibody (Cat#Ab955, Abcam, Cambridge, London, UK) overnight at 4 °C, and then incubated with peroxidase-conjugated secondary antibody for 2 hours at 37 °C. Finally, sections were stained with 3,3’-diaminobenzidine, then examined under a light microscope. A Vector Laboratories Mouse Ig Blocking Reagent (Cat#MKB-2213, Vector Laboratories, Vector Laboratories, Burlingame, CA, USA) was used to aid in the detection of mouse antigen using mouse primary antibodies.

Spt-GC staining was performed according to a method described by Soni *et al*. (Soni *et al.*, 2014). Briefly, spleen tissues dissected from three groups of 20-wk old female mice (B6.WT + untreated, B6.*Fas^lpr^* + LV-miR-shRNA-ctrl and B6.*Fas^lpr^* + LV-miR-574-5p-shRNA) were sliced into serial 10 µm transverse cryosections. GC-B cells and follicle B cells were stained with Pna-HRP (Sigma) and anti-mouse IgD-BIOT (SouthernBiotech, 1:100 dilution in blocking buffer, supplemental Table S8) together with a Streptavidin Alkaline Phosphatase kit (Cat#SA-5100, Vector Laboratories) and a Vector Blue Reagent kit (Cat#SK5300, Vector Laboratories) and a Vector Red Reagent kit (Cat#SK4800, Vector Laboratories) as instructed. Finally the slides were mounted with Vectamount (H-5000, Vector Laboratories) and examined under a light microscope.

For other fluorescence confocal microscopy, five-micrometer cryosections were fixed in ice-cold acetone, subsequently permeabilized and blocked with 0.1% Triton X-100 and 5% bovine serum albumin. Rabbit anti-mouse C3 antibody (1:10) and rabbit anti-mouse-IgG (1:100) were used as the primary antibodies. After washing three times, the slides were incubated with Alexa Fluor^®488^goat anti-rabbit IgG, (1:200, Cat#, Invitrogen, Carlsbad, CA, USA). The slides were mounted using 15% Mowiol (Cat#, Sigma, St. Louis, MO, USA). Stained images were obtained at the same light exposure using confocal laser-scanning microscopy (Leica TCS SP8, Germany).

### Serum ANA assays

Briefly, HEp2 cells were seeded into 96-well plates (1×10^4^/well) and grown for 24 h. Cells were washed three times with PBS (5 min/each) and fixed with 4% paraformaldehyde at room temperature for 10 min. Washing with PBS three times again, the cells were incubated with 0.2% Triton X-100 and 0.2% albumin bovine V (BSA) in PBS at 4°C for 10 min to permeate the cell membrane. Subsequently, a blocking solution (0.02% Triton X-100 and 5% BSA in PBS) was added and the plates were incubated at 37°C for 30 min.

For the detection of ANA in the mouse sera, the above treated HEp2 cells were incubated with diluted sera from three groups of 20-wk old mice as described above (B6.WT + untreated, B6.*Fas^lpr^ +* LV-miR-shRNA-ctrl and B6.*Fas^lpr^ +* LV-miR-574-5p-shRNA) respectively in the blocking solution at a final dilution of 1:100 at 4 °C for 16 h. The cells were washed three times with PBS and incubated with peroxidase-conjugated anti-mouse antibody for 2 h at 37°C. After washing thrice with PBS, cells were visualized using the chromogenic substrate 3-3’ diaminobenzidine, then examined under a light microscope.

Alternatively, sera from the B6.*hMIR574^Tg^* and the control B6.WT mice treated with or without Dox were diluted 1:5 and incubated with HEp2 cells for 60 minutes. Cells were subsequently washed with PBS, and re-incubated with an Alexa Fluor488 labeled goat anti-mouse lgG (HþL) for additional 60 minutes. Cell immunofluorescence was recorded under a confocal fluorescence microscopy.

### Clinical samples and analyses

All clinical samples were collected with the informed consent of the patients and study protocols that were in accordance with the ethical guidelines of the Declaration of Helsinki (as revised in Edinburgh 2000) and were approved by the Institutional Medical Ethics Committee of the Dalian Medical University. Human SLE patients and healthy controls were recruited by the Second Hospital, Dalian Medical University, Dalian, China. All SLE patients fulfilled the 1997 American College of Rheumatology revised criteria for SLE. Patients with an obvious sign of infection within a month (e.g., those with elevated serum C-reactive protein) were excluded. 47 SLE patients were enrolled in this study. 18 controls were normal subjects without a history of major diseases such as cancer, diabetes, connective tissue disease. SLE disease activity index (SLEDAI) was scored for 47 SLE patients by using a combination of the clinical history, physical examination, organ specific functional tests, and serologic studies ^69^. Fasting morning blood was collected. Clinical laboratory studies including C-reactive protein, serum creatinine and urea nitrogen, liver function, urinalysis and immunologic tests were performed per standard protocols in the Clinical Laboratory Department, the Second Hospital, Dalian Medical University. Serum levels of miR-574-5p for 47 SLE patients and 18 normal controls were determined by qPCR as described in the Extended Materials and Methods in the Supplementary Information. For the isolation and analyses of human PBMCs, blood samples from multiple healthy human individuals were combined.

### Other data acquisition, image processing and statistical analyses

Western blot images were captured by Biosense SC8108 Gel Documentation System with GeneScope V1.73 software (Shanghai BioTech, Shanghai, China). Gel images were imported into Photoshop for orientation and cropping. Data are the means ± SEM. One-way ANOVA with Bonferonni’s post-test was used for multiple comparisons and unless indicated otherwise, the unpaired Student’s *t-*test (two-tailed) for pair-wise comparisons.

## ACKNOWLEDGMENTS

The authors thank Dr. Jiahuai Han for the provision of hTLR7/8/9 plasmids and wild-type and MyD88 deficient RAW264.7 cells and Dr. Rongtuan Lin for the provision of the plasmid reporters for interferon activity. The authors thank Yang other lab members for their assistance with certain parts of the experiments.

## FUNDING

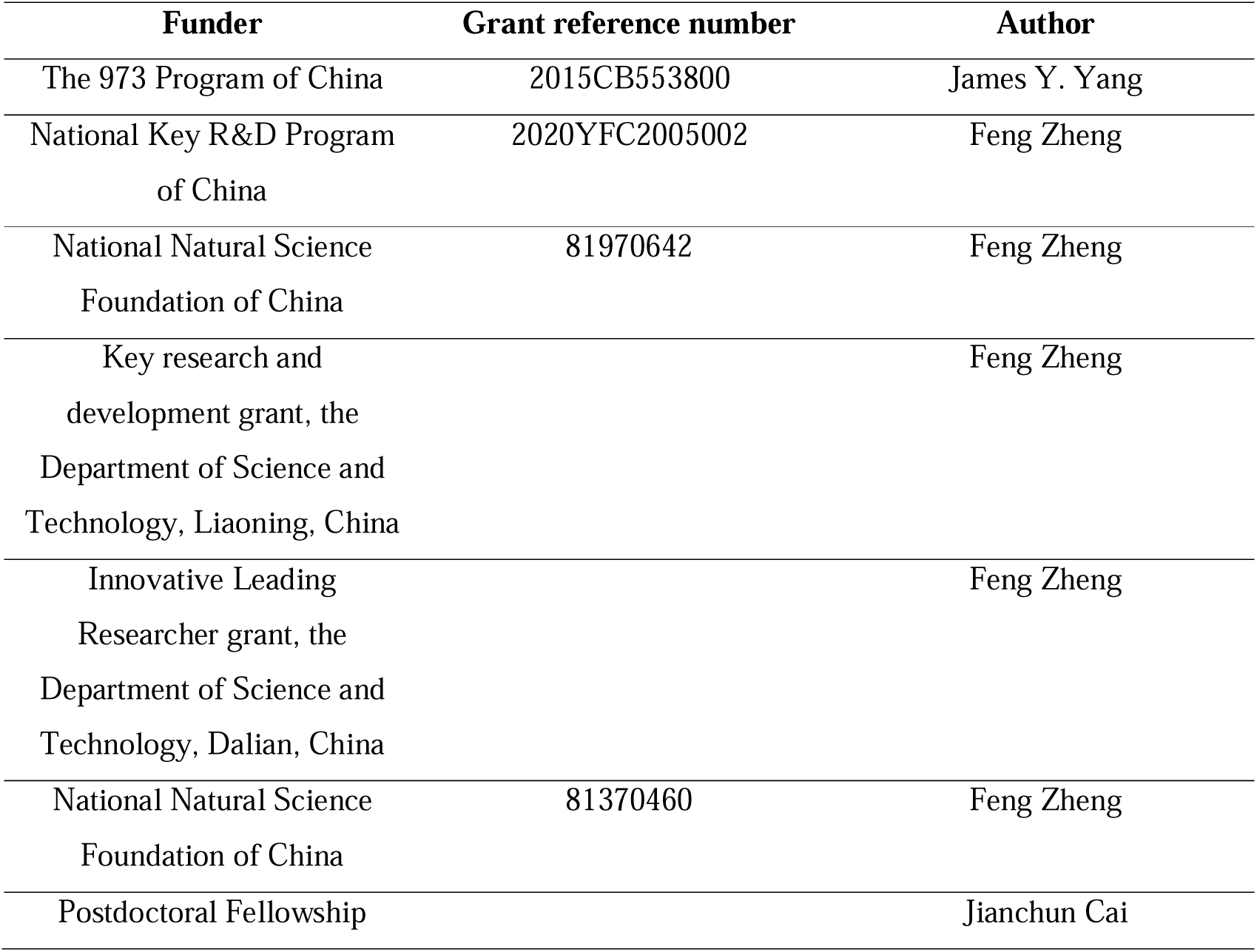

| **Funder** | **Grant reference number** | **Author** |
| --- | --- | --- |
| The 973 Program of China | 2015CB553800 | James Y. Yang |
| National Key R&D Program<br>of China | 2020YFC2005002 | Feng Zheng |
| National Natural Science<br>Foundation of China | 81970642 | Feng Zheng |
| Key research and<br>development grant, the<br>Department of Science and<br>Technology, Liaoning, China |  | Feng Zheng |
| Innovative Leading<br>Researcher grant, the<br>Department of Science and<br>Technology, Dalian, China |  | Feng Zheng |
| National Natural Science<br>Foundation of China | 81370460 | Feng Zheng |
| Postdoctoral Fellowship |  | Jianchun Cai |

## AUTHOR CONTRIBUTION

Tao Wang, Dan Song, Xuejuan Li, Yu Luo, Dianqiang Yang, Xiaoyan Liu, Xiaodan Kong, Yida Xing, Shulin Bi, Yan Zhang, Tao Hu, Yunyun Zhang, Shuang Dai, Zhiqiang Shao, Dahan Chen, Jinpao Hou: Methodology, Investigation, Data curation, Software, Formal Analysis, Validation; Esteban Ballestar: Formal Analysis, Reviewing and Editing; Jianchun Cai: Funding acquisition, Supervision, Formal Analysis, Reviewing and Editing; Feng Zheng: Conceptualization, Funding Acquisition, Supervision, Reviewing and Editing; James Y. Yang: Conceptualization, Funding acquisition, Supervision, Original draft, Reviewing and Editing

## ETHICS

The use of animals and the study protocols were reviewed and approved by the Institutional Animal Use and Care Committees of Xiamen University and Dalian Medical University.

All clinical samples were collected with the informed consent of the patients and study protocols that were in accordance with the ethical guidelines of the Declaration of Helsinki (1975) and were approved by the Institutional Medical Ethics Committee of Dalian Medical University.

## DATA AVAILABILITY

Two sets of sequencing data (GSE235926 for miR-574-5p knockdown HeLa cells and GSE236337 for miR-574-5p transfected mouse splenocytes) reported in this article are accessible online on the Gene Expression Omnibus.

