## Supplementary Material for "MiR-574-5p activates human TLR8 to promote autoimmune signaling and lupus"

### *SUPPLEMENTARY INFORMATION*

***Extended Materials and Methods***

- *Real-time quantitative PCR (qPCR)*
- *Western blots or immunoblots.*
- *ELISA assays of human or mouse interferons, cytokines, serum anti-dsDNA autoantibody and other proteins.*
- *mRNA microarray analyses in miR-574-5p-knockdown HeLa cells.*
- *RNA-Seq analyses in Dotap-PS-miR-574-5p-transfected splenocytes.*

***Supplemental Figures***

- *Figure S1 Co-localization of miR-574-5p and mTlr7 in mouse peritoneal macrophages isolated from B6.WT and B6.Tlr7$^{-/-}$ mice.*
- *Figure S2 miR-574-5p exposure significantly stimulated TNFα secretion and the mRNA expression of markers for B cell activation in human PBMCs.*
- *Figure S3 Specific activation of hTLR8 but not hTLR7 by miR-574-5p.*
- *Figure S4 miR-574-5p exposure potently stimulated Tnfα secretion in mouse bone marrow derived dendritic cells (mBMDCs).*
- *Figure S5 Gene expression profiling in miR-574-5p knockdown in HeLa cells.*
- *Figure S6 RNA-Seq analyses in miR-574-5p-transfected mouse splenocytes.*
- *Figure S7 Characterization of miR-574-overexpressing transgenic mice.*
- *Figure S8 Effects of lentivirus-mediated silencing of miR-574-5p on the developlemt of lupus and lupus nephritis-1.*

- *Figure S9 Effects of lentivirus-mediated silencing of miR-574-5p on the developlemt of lupus and lupus nephritis-2.*

***Supplemental Tables***


- *Table S1 The general characteristics of 47 SLE patients and 18 healthy controls.*
- *Table S2 A summary of sequencing data for six splenocyte samples.*
- *Table S3 Alignment statistics of reads alignment to the reference genes.*
- *Table S4 Alignment statistics of reads alignment to the reference genome.*
- *Table S5 A list of lentiviral shRNA vectors for miRNAs and mRNAs.*
- *Table S6 Plasmids and the primers used for their construction.*
- *Table S7 A list of chemically-synthesized and HPLC-purified miRNAs.*
- *Table S8 A list of the antibodies used for flow cytometry, immunoblots (IB), immunoprecipitation (IP), immunohistochemistry (IHC) or immunofluorescence (IF).*
- *Table S9 A list of the primers used for RT-PCR or qPCR analyses of mRNAs and miRNAs.*
- *Table S10 A list for the ELISA kits used.*


- ***Supplemental References***

#### *Extended Materials and Methods*

***Real-time quantitative PCR*** (***qPCR***). For qPCR analyses of mRNA, reverse transcription was performed with TRIzol (Invitrogen, Carlsbad, CA, USA)-extracted total RNAs using a ReverTra Ace-α® Kit as instructed (Cat#FSQ-101, Toyobo, Tokyo, Japan). qPCR was performed using the SYBR Green Real-Time PCR Master Mix (Cat#QPK-212, Toyobo) and the Step One Plus Real-Time PCR system (Applied Biosystems Inc., Foster City, CA, USA) using appropriate primer pairs as listed in supplemental Table S9, according to the manufacturers' protocols and with 18S rRNA as a control.

For miRNA assays, serum total RNAs was extracted using a mirVana miRNA isolation kit (Cat#AM1556, Ambion, Austin, TX, USA) whereas total RNAs from cultured cells or tissues were extracted using TRIzol, according to the manufacturer's protocols. Five microliters of total RNA was reverse transcribed using the ReverTra Ace-α-® Kit as instructed (TOYOBO, Shanghai, China) and miRNA-specific stem-loop primers listed in supplemental Table S9. qPCR was performed with total RNAs, using universal primer and miRNA-specific reverse LNA-primers as listed in supplemental Table S9, with U6 RNA served as an internal control. For human serum miR-574-5p analyses by qPCR, serum samples were prepared from blood samples collected from human SLE patients or healthy controls.

***Western blots or immunoblots.*** Western blots or immunoblots were performed by standard protocols using antibodies listed on supplemental Table S8.

***ELISA assays of human or mouse interferons, cytokines, serum anti-dsDNA autoantibody and other proteins.*** The levels of IFNα/β/γ, TNFα, IL6, IgG/1/2b, IgM, Trim21/Ro52 and Sp100, serum anti-dsDNA and serum anti-RNA were analyzed for supernatants of cell cultures or serum samples using ELISA kits as listed in supplemental Table S10 according to the manufacturer's instructions.

***mRNA microarray analyses in miR-574-5p-knockdown HeLa cells.*** HeLa cells were seeded on plates at the density of 20-30% confluency and incubated overnight. Lentiviral infection of HeLa cells was performed with lentiviruses LV-miR-574-5p-shRNA or LV-miR-shRNA-ctrl at multiplicity of infection of 1:1 and in the presence of Polyberene (Cat#107689, Sigma-Aldrich, St Louis, MO, USA). 96 h after viral infection, cells were harvested and total RNAs were prepared for miR-574-5p and mRNA expression analyses by qPCR and microarray analyses.

Microarray analyses of mRNA expression of miR-574-5p-knockdown HeLa cells were performed with NimbleGen 12x135K microarrays (Roche NimbleGen, Inc., Madison, WI, USA) by the KangChen Biotech (Shanghai, China). Briefly, total RNA of each sample was used for labeling and array hybridization as the following steps: 1) Reverse transcription with Invitrogen Superscript ds-cDNA synthesis kit; 2) ds-cDNA labeling with NimbleGen one-color DNA labeling kit; 3) Array hybridization using the NimbleGen Hybridization System and followed by washing with the NimbleGen wash buffer kit; 4) Array scanning using the Axon GenePix 4000B microarray scanner (Molecular Devices Corporation). Scanned images were then imported into NimbleScan software (version 2.5) for grid alignment and expression data analysis. Expression data were normalized through quantile normalization and the Robust Multichip Average algorithm included in the NimbleScan software. Further data analyses were performed using Agilent GeneSpring GX v11.5.1 software. Significant differentially expressed genes (DEGs) were identified through Volcano Plot filtering. Gene ontology analyses and pathway analyses by Agilent GeneSpring GX v11.5.1 or the Reactome V58 (http://www.reactome.org/, projected to human) were performed to determine the enrichment of these DEGs in signaling or disease pathways.

***RNA-Seq analyses in Dotap-PS-miR-574-5p-transfected splenocytes.*** Mouse splenocytes were isolated from three 8-wk old female B6.WT mice, with each sample divided into two equivalent aliquots, one for Dotap-only and one for Dotap-PS-miR-574-5p transfection. Splenocytes were cultured in DMEM supplemented with 10% FBS, 2 mM L-glutamine and 100 IU/ml penicillin and 100 μg/ml streptomycin for 4 h. Subsequently, cultured splenocytes were transfected with Dotap-only or Dotap-PS-miR-574-5p (Dotap-only-1, -2 and -3 and Dotap-miRNA-1, -2 and -3) at the concentration of 10 μg/ml. After 24 h of incubation, cells were harvested for RNA extraction and analyses.

RNA extraction, library preparations and RNA-Seq for the six groups of splenocytes were performed by the BGI Co. Ltd. (Shenzhen, China). Briefly, total RNA was isolated from six groups of splenocytes using TRIzol Reagent (Invitrogen, Life Technologies) according to the manufacturer's protocol, including treatment with DNaseI (Invitrogen, Life Technologies). Starting with approximately 1 μg of total RNA/sample, mRNA was enriched by oligo-(dT) magnetic beads to select for mRNA with poly-A tail, and broken into short fragments (about 130-160 bp) with a disruption buffer at high temperature. The mRNA fragments were reverse-transcribed into double stranded cDNA using the random hexamers. cDNAs were subsequently end-repaired by T4 polynucleotide kinase, T4 DNA polymerase and the Klenow fragment (Y904-1500, P708-1500, P706-300, Enzymatics, Beverly, MA, USA), with 5'-phosphorylated and a sticky "A" at the 3'-end. T-tailed sequencing adaptors were attached to the A-tailed cDNAs by T4 DNA ligase T4 DNA ligase (Cat#L603-HC-1500, Enzymatics). The adaptor-attached fragments were enriched by PCR amplification, and denatured into the single-stranded cDNAs. The single-stranded cDNAs were circularized by the splint oligonucleotides and DNA ligase.

The six circular single-stranded DNA libraries were sequenced by single-ended 50-base (SE50) sequencing using the BGIseq500 platform (BGI, Shenzhen, China), generating about 24,065,298 raw sequencing reads/sample. Following exclusion of low-quality reads, on average 23,910,522 clean reads/sample were obtained. The clean reads were mapped to the UCSC Genome Reference Consortium Mouse Build 38 using the HISAT (Kim et al., 2015) and the Bowtie2 tools (Langmead et al., 2009). The average mapping ratio with the reference genes was 70.72% and the average mapping ratio with the reference genome was 93.21%. The sequencing passed all the quality control analyses.

The gene expression levels were determined by the RSEM software package (Li and Dewey, 2011)and expressed as fragments per kilobase of transcript per million mapped reads (FPKM). Significant DEGs in three individual pairs of samples (Dotap-only-1 versus Dotap-miR-1; Dotap-only-2 versus Dotap-miR-2 and Dotap-only-3 versus Dotap-miR-3) were identified by the method of Poisson distribution, which was largely based on Audic S. et al. (Audic and Claverie, 1997). A false discovery rate (FDR) >= 0.001 and an absolute value of

$log_2$[(Dotap-PS-miR-574-5p treated)/(Dotap-only treated)] >= 1 were used as the thresholds to determine the significance of the gene expression difference. DEGs that were upregulated or down-regulated by at least 2-fold between the combined controls and the combined treatment samples were identified by the NOISeq method (Tarazona et al., 2011). Further data analyses including gene ontology (http://www.geneontology.org/) and KEGG (http://www.genome.jp/kegg/) and Reactome pathway enrichment analyses were performed as described.

### *Supplemental Figures*

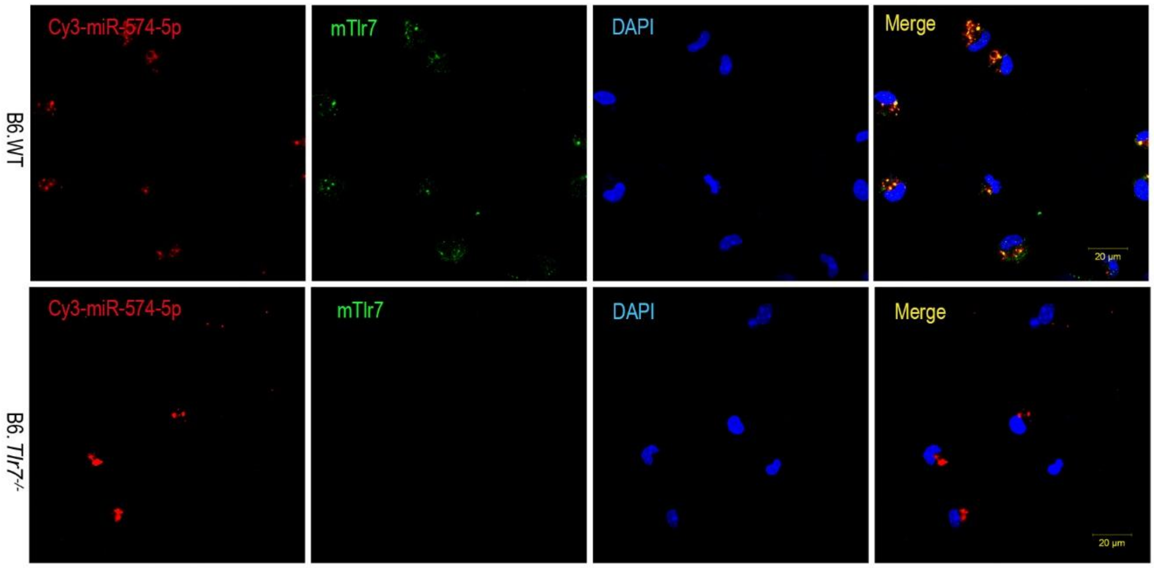


***Figure S1 Co-localization of miR-574-5p and mTlr7 in mouse peritoneal macrophages isolated from B6.WT and B6.Tlr7<sup>-/-</sup> mice***. Mouse peritoneal cells were harvested by peritoneal lavage with 8-10 mL of ice-cold PBS. Peritoneal macrophages were 400×g centrifuged for 5 minutes and the resulting peritoneal macrophages were plated at $1 \times 10^5$ cells/mL in DMEM supplemented with FBS (10%), L-glutamine (2 mM) and penicillin/streptomycin (100 U/mL, 100 μg/mL) (all purchased from Sangon Biotech, Shanghai, China). Peritoneal macrophages were cultured overnight prior to transfection or stimulation. Cells were treated with 3 μg Dotap-formulated Cy3-labelled miR-574-5p (Genscript, Nanjing, China) and incubated for 12 h in darkness. Stained cells were examined under a Zeiss LSM 780 confocal microscope. Magnification 100×; Scale bar, 20 μm.

***a***

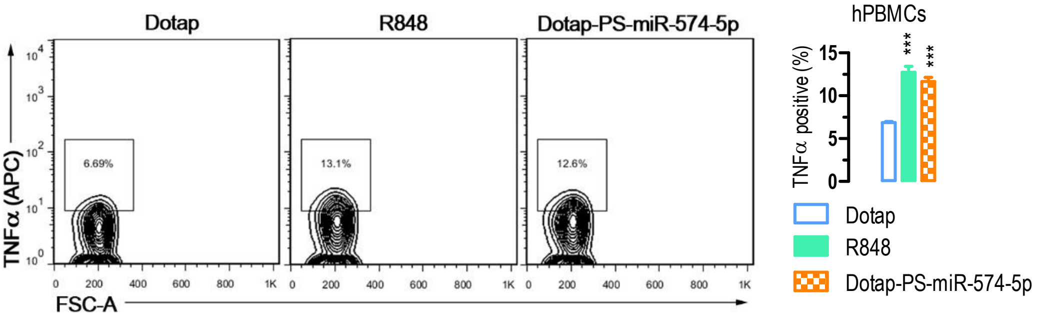


***b***

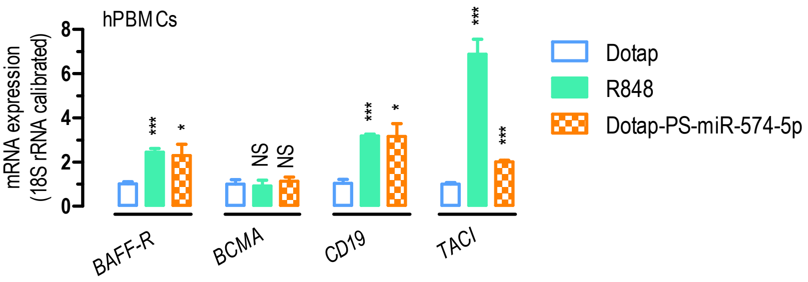


***Figure S2 miR-574-5p exposure significantly stimulated TNFα secretion and the mRNA expression of markers for B cell activation in human PBMCs***. About $1\times10^6$ human PBMCs were seeded in 6-well plate and treated with 1 μg/ml R848 or 10 μg/ml of Dotap-PS-miR-574-5p for 24 h. Subsequently, the cells were harvested for flow cytometry or qPCR analyses. NS, not significant; * $P < 0.05$, ** $P < 0.01$, *** $P < 0.001$, compared with Dotap only. (***a***) miR-574-5p exposure significantly increased the percentage of TNFα-secreting human PBMCs as determined by flow cytometry analyses ($n = 3$). (***b***) Stimulation of BAFF-R, CD19 and TACI but not B-cell maturation antigen (BCMA) mRNA expression by miR-574-5p in human PBMCs as determined by qPCR analyses ($n = 3\text{-}6$).

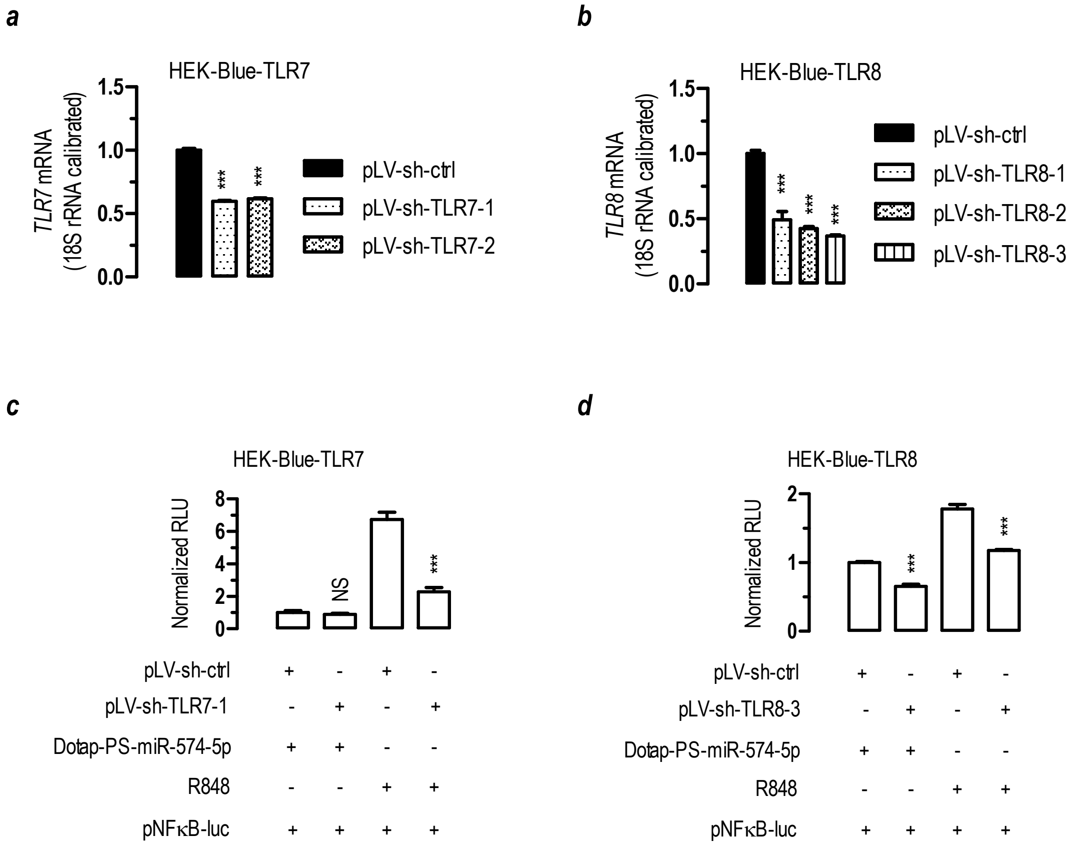


***Figure S3 Specific activation of hTLR8 but not hTLR7 by miR-574-5p.*** Luciferase reporter assays were performed as described. NS, not significant; ***, $P < 0.001$. (***a***) TLR7 knocking-down by shRNA constructs in HEK-Blue-TLR7 cells ($n$ = 3-4). Cells were transfected as indicated. 24 hours after the transfection, cells were harvested for qPCR analysis. Statistical comparisons were made for pLV-sh-ctrl versus pLV-sh-TLR7-1/2. (***b***) TLR8 knocking-down by shRNA constructs in HEK-Blue-TLR8 cells ($n$ = 3-4). Cells were transfected as indicated. 24 hours after the transfection, cells were harvested for qPCR analysis. Statistical comparisons were made between pLV-sh-ctrl versus pLV-sh-TLR8-1/2/3. (***c***) Knocking down TLR7 significantly suppressed activation of NFκB induced by R848 but not miR-574-5p ($n$ = 3). HEK-Blue-TLR7 cells were co-transfected with pLV-sh-TLR7-1 or pLV-sh-ctrl and a luciferase reporter plasmid (pNFκB-luc) plus pSV40-β-galactosidase (4:3:1) and grown for 24 h. Cells were subsequently stimulated with 10 μg/ml of Dotap-PS-miR-574-5p or 1 μg/ml of R848 for 8 h. Statistical comparisons were made for pLV-sh-ctrl + Dotap-

PS-miR-574-5p versus pLV-sh-TLR7-1 + Dotap-PS-miR-574-5p or pLV-sh-ctrl + R848 versus pLV-sh-TLR7-1 + R848. (***d***) Knocking down TLR8 significantly suppressed activation of NFκB induced by both R848 and miR-574-5p (*n* = 3). HEK-Blue-TLR8 cells were co-transfected with pLV-sh-TLR8-3 or pLV-sh-ctrl and a luciferase reporter plasmid (pNFκB-luc) plus pSV40-β-galactosidase (4:3:1) and grown for 24 h. Cells were subsequently stimulated with 10 μg/ml of Dotap-PS-miR-574-5p or 1 μg/ml of R848 for 8 h. Statistical comparisons were made for pLV-sh-ctrl + Dotap-PS-miR-574-5p versus pLV-sh-TLR8-3 + Dotap-PS-miR-574-5p or pLV-sh-ctrl + R848 versus pLV-sh-TLR8-3 + R848.

***a***

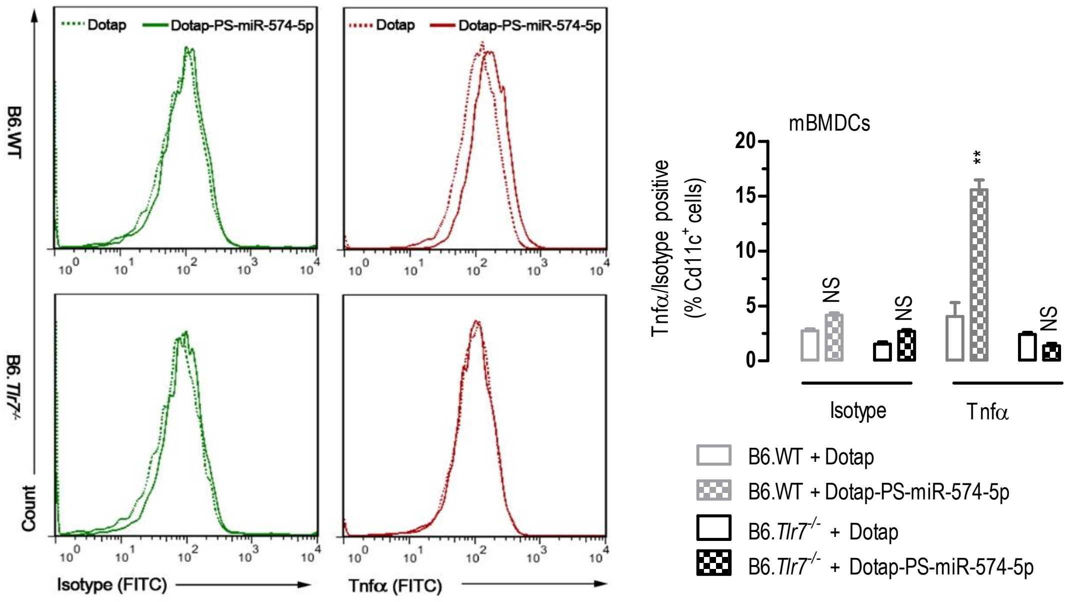

Dotap
Dotap-PS-miR-574-5p
B6.WT
B6.Tlr7-/-
Count
Isotype (FITC)
Tnfα (FITC)
mBMDCs
Tnfα/Isotype positive (% Cd11c+ cells)
NS
**
Isotype
Tnfα
B6.WT + Dotap
B6.WT + Dotap-PS-miR-574-5p
B6.Tlr7-/- + Dotap
B6.Tlr7-/- + Dotap-PS-miR-574-5p


***b***

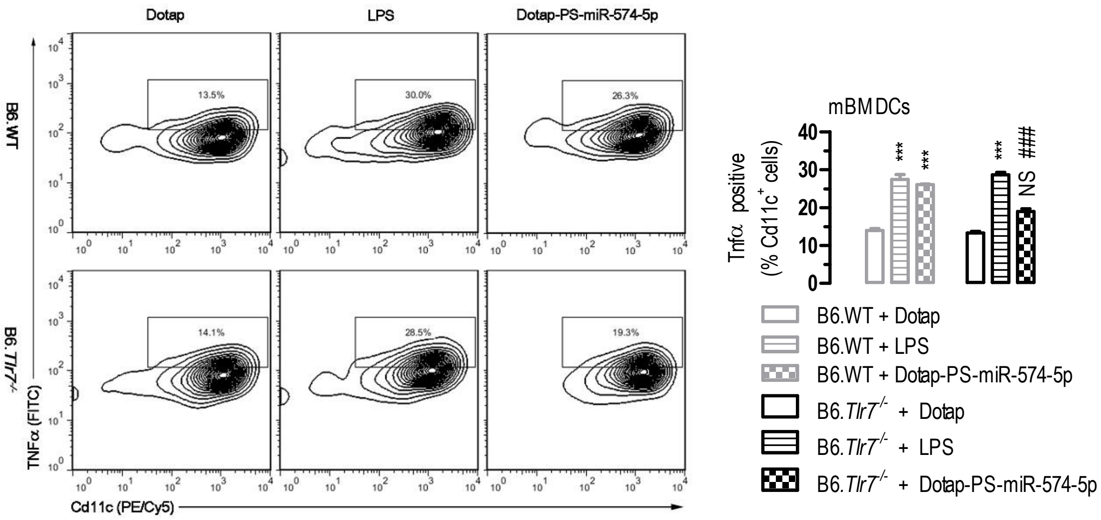

Dotap
LPS
Dotap-PS-miR-574-5p
B6.WT
B6.Tlr7-/-
13.5%
30.0%
26.3%
14.1%
28.5%
19.3%
TNFα (FITC)
Cd11c (PE/Cy5)
mBMDCs
Tnfα positive (% Cd11c+ cells)
***
NS
###
B6.WT + Dotap
B6.WT + LPS
B6.WT + Dotap-PS-miR-574-5p
B6.Tlr7-/- + Dotap
B6.Tlr7-/- + LPS
B6.Tlr7-/- + Dotap-PS-miR-574-5p


***c***

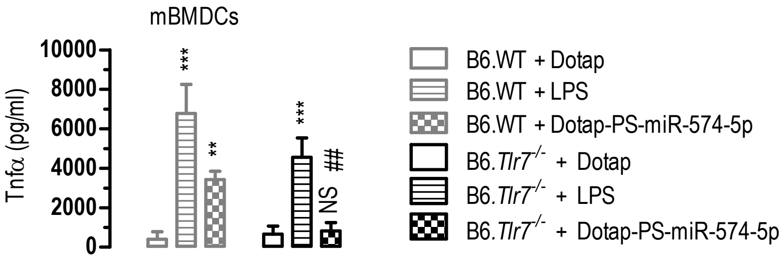

mBMDCs
Tnfα (pg/ml)
***
**
NS
##
B6.WT + Dotap
B6.WT + LPS
B6.WT + Dotap-PS-miR-574-5p
B6.Tlr7-/- + Dotap
B6.Tlr7-/- + LPS
B6.Tlr7-/- + Dotap-PS-miR-574-5p

***Figure S4 miR-574-5p exposure potently stimulated Tnfα secretion in mouse bone marrow derived dendritic cells (mBMDCs)***. About $1\times10^5$ or $1\times10^6$ mBMDCs prepared from either the B6.WT or B6.*Tlr7*$^{-/-}$ mice were seeded in 6-well plates and treated with 100 ng/ml LPS or 10 μg/ml Dotap-PS-miR-574-5p for 24 h ($n = 3$). (***a***) Tnfα secretion in response to miR-574-5p stimulation in mBMDCs prepared from the B6.WT or B6.*Tlr7*$^{-/-}$ mice as analyzed by flow cytometry with an isotype antibody as a control. Cells were gated by Cd11c$^+$ and isotype-antibody positive respectively. NS, not significant; ** $P < 0.01$, compared to B6.WT + Dotap or B6.*Tlr7*$^{-/-}$ + Dotap. (***b***) miR-574-5p stimulated Tnfα secretion in mBMDCs prepared from the B6.WT but not B6.*Tlr7*$^{-/-}$ mice whereas LPS stimulated Tnfα secretion in mBMDCs isolated from both the B6.WT and the B6.*Tlr7*$^{-/-}$ mice, as analyzed by flow cytometry. Cells were gated by Cd11c$^+$. NS, not significant; *** $P < 0.001$, compared to B6.WT + Dotap or B6.*Tlr7*$^{-/-}$ + Dotap; ### $P < 0.001$, B6.WT + Dotap-PS-miR-574-5p versus B6.*Tlr7*$^{-/-}$ + Dotap-PS-miR-574-5p. (***c***) miR-574-5p exposure significantly increased Tnfα secretion by mBMDCs prepared from the B6.WT mice but not the B6.*Tlr7*$^{-/-}$ mice as analyzed by ELISA. NS, not significant; ** $P < 0.01$, *** $P < 0.001$, compared to B6.WT + Dotap or B6.*Tlr7*$^{-/-}$ + Dotap; ### $P < 0.001$, B6.WT + Dotap-PS-miR-574-5p versus B6.*Tlr7*$^{-/-}$ + Dotap-PS-miR-574-5p.

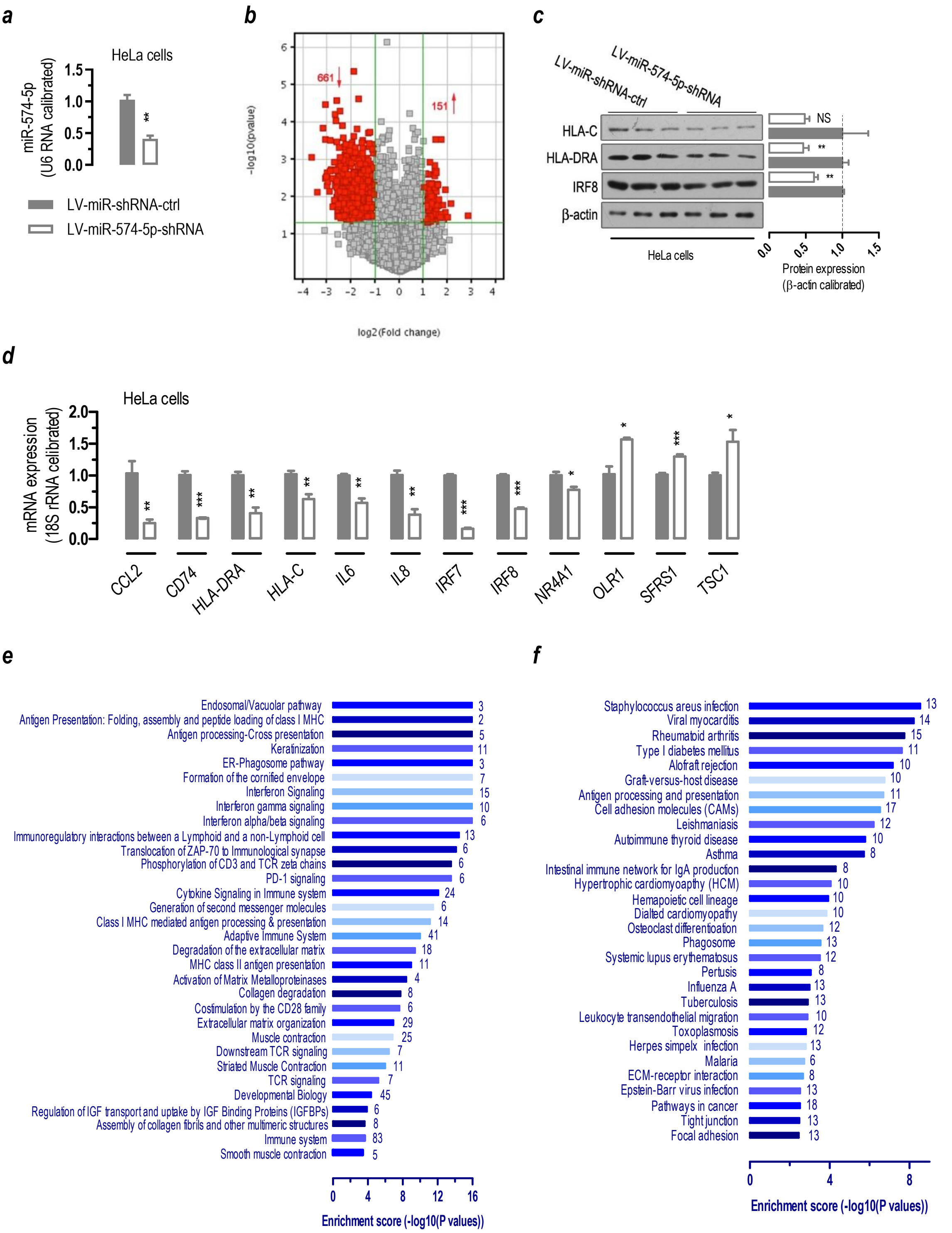


***Figure S5 Gene expression profiling in miR-574-5p knockdown in HeLa cells.*** NS, not significant; * $P < 0.05$, ** $P < 0.01$, *** $P < 0.001$, LV-miR-shRNA-ctrl versus LV-miR-574-5p-shRNA. (***a***) Lentivirus-mediated knockdown of miR-574-5p in HeLa cells for microarray analyses. HeLa cells were seed and infected with LV-miR-574-5p-shRNA or control virus. 96 hours after the infection, cells were harvested for analysis (*n* = 3). (***b***) A volcano plot showing mRNA expression in miR-574-5p knockdown HeLa cells (*n* = 3). HeLa cells were infected with lentiviruses carrying shRNAs against miR-574-5p to knock-down miR-574-5p Gene expression analyses were performed with NimbleGen 12×135K microarrays as described in the Extended Materials and Methods. Further data analyses were performed using the Agilent GeneSpring GX v11.5.1 software or the Reactome V58 (http://www.reactome.org/). The red point in the plot represented the DEGs with statistical significance. The vertical green lines in the plot divided genes that were up-regulated (151 genes) or down-regulated (661 genes) by at least 2 folds, respectively, whereas the horizontal green line represents a *P*-value of 0.05. (***c***) Protein expression of selected genes in miR-574-5p knockdown HeLa cells as determined by Western blots. (***d***) mRNA expression of selected genes in miR-574-5p knockdown HeLa cells as determined by qPCR (*n* = 3-5). *CCL2*, (C-C Motif) ligand-2; *CD74,* cluster of differentiation-74; *HLA-DRA*, HLA class II histocompatibility antigen, DR alpha chain; *HLA-C,* major histocompatibility complex, Class I, C; *IL8,* interleukin-8; *IRF8,* interferon regulatory factor-8; *NR4A1,* nuclear receptor subfamily-4, group-A, member-1; *OLR1,* oxidized low-density lipoprotein receptor-1; *SFRS1,*serine/arginine-rich splicing factor-1; *TSC1,* tuberous sclerosis-1. (***e***) 32 top signaling pathways that were enriched for the significantly-down-regulated genes in miR-574-5p knockdown HeLa cells as analyzed by the Reactome package (http://www.reactome.org/). The digital number next to each bar represented the count of the significantly-down-regulated genes enriched in the pathway. (***f***) 30 top biological or disease pathways that were enriched for the significantly-down-regulated genes in miR-574-5p knockdown HeLa cells as analyzed by the Agilent GeneSpring package. The digital number next to each bar represented the count of the significantly down-regulated genes enriched in the pathway.

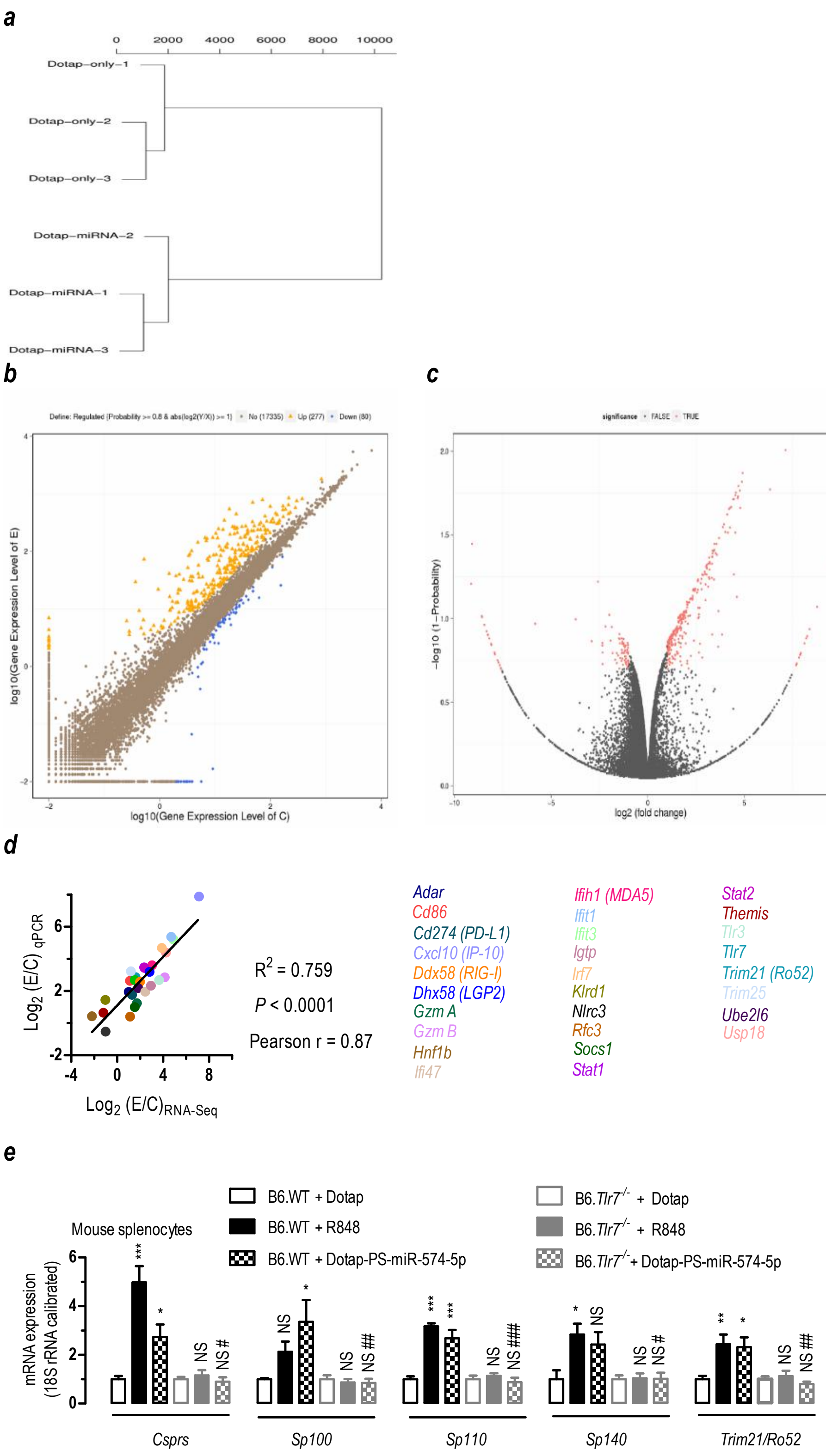
a
0
2000
4000
6000
8000
10000
Dotap–only–1
Dotap–only–2
Dotap–only–3
Dotap–miRNA–2
Dotap–miRNA–1
Dotap–miRNA–3
b
log10(Gene Expression Level of E)
log10(Gene Expression Level of C)
c
significance
FALSE
TRUE
-log10 (1-Probability)
log2 (fold change)
d
Log2 (E/C) qPCR
Log2 (E/C)RNA-Seq
R2 = 0.759
P < 0.0001
Pearson r = 0.87
Adar
Cd86
Cd274 (PD-L1)
Cxcl10 (IP-10)
Ddx58 (RIG-I)
Dhx58 (LGP2)
Gzm A
Gzm B
Hnf1b
Ifi47
Ifih1 (MDA5)
Ifit1
Ifit3
Igtp
Irf7
Klrd1
Nlrc3
Rfc3
Socs1
Stat1
Stat2
Themis
Tlr3
Tlr7
Trim21 (Ro52)
Trim25
Ube2l6
Usp18
e
Mouse splenocytes
B6.WT + Dotap
B6.WT + R848
B6.WT + Dotap-PS-miR-574-5p
B6.Tlr7-/- + Dotap
B6.Tlr7-/- + R848
B6.Tlr7-/- + Dotap-PS-miR-574-5p
mRNA expression
(18S rRNA calibrated)
Csprs
Sp100
Sp110
Sp140
Trim21/Ro52

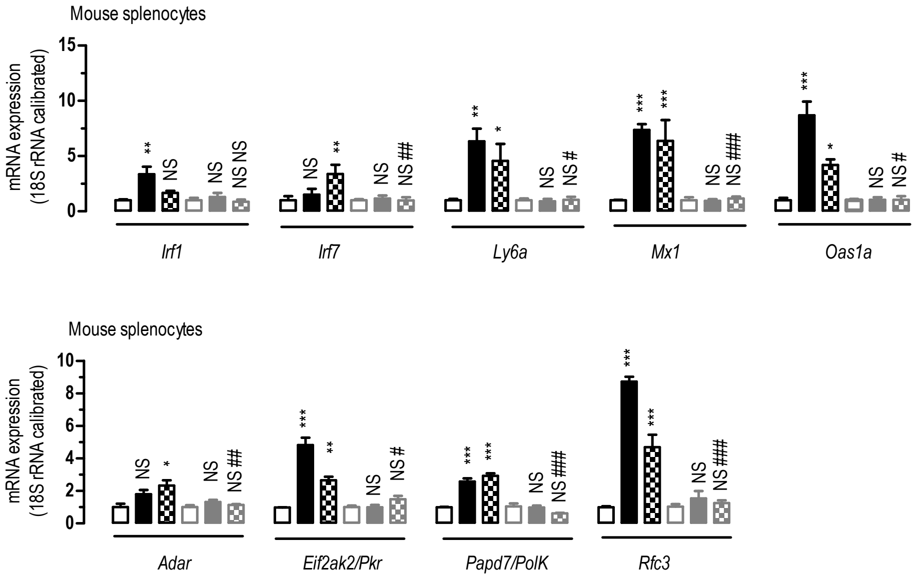


***Figure S6 RNA-Seq analyses in miR-574-5p-transfected mouse splenocytes.*** (***a***) A distance tree showing the relationship of expressed genes in six splenocyte samples as determined by the Euclidean method. (***b***) A scatter plot of showing the DEGs between the Dotap-PS-miR-574-5p treated and Dotap-only treated splenocytes ($n$ = 3). A false discovery rate (FDR) >= 0.001 and an absolute value of $\log_2$[(Dotap-PS-miR-574-5p treated)/(Dotap-only treated)] >= 1 were used as the thresholds to determine the significance of the gene expression difference. Blue dots represent down-regulated genes while orange dots represent up-regulated gene and brown dots represent those that are not significantly altered. In comparison with the Dotap-only treated cells, 277 genes were significantly upregulated whereas 80 genes were down-regulated in Dotap-PS-miR-574-5p treated splenocytes. E, Dotap-PS-miR-574-5p treated, C, Dotap-only treated. (***c***) A volcano plot showing the DEGs the Dotap-PS-miR-574-5p treated and Dotap-only treated splenocytes. Red dots represented DEGs which passed screening threshold and those in black did not. (***d***) A high level of concordance of the differential expression measurements between qPCR and RNA-Seq at varying abundances. qPCR analyses were performed with 28 selected mRNAs (which included 4 down-regulated genes and 24 up-regulated genes as determined by the RNA-Seq) in Dotap-PS-miR-574-5p treated and Dotap-only treated splenocytes as described. (***e***) miR-574-5p-mediated induction of autoantigens, ISGs and other genes is mTlr7-dependent. Splenocytes were isolated from 8-wk female B6.WT and B6.*Tlr7*$^{-/-}$ mice and were treated with control Dotap, 1 μg/ml R848 or 10 μg/ml miR-574-5p (Dotap-PS-miR-574-5p) for 24 h. Total RNAs were extracted from the treated cells and subjected to qPCR using primers as listed in supplemental Table S9. NS, not significant; * $P < 0.05$, ** $P < 0.01$, *** $P < 0.001$, compared either to B6.WT + Dotap or B6.*Tlr7*$^{-/-}$ + Dotap; # $P < 0.05$, ## $P < 0.01$, ### $P < 0.001$, B6.WT + Dotap-PS-miR-574-5p versus B6.*Tlr7*$^{-/-}$ + Dotap-PS-miR-574-5p; $n = 3\text{-}4$.

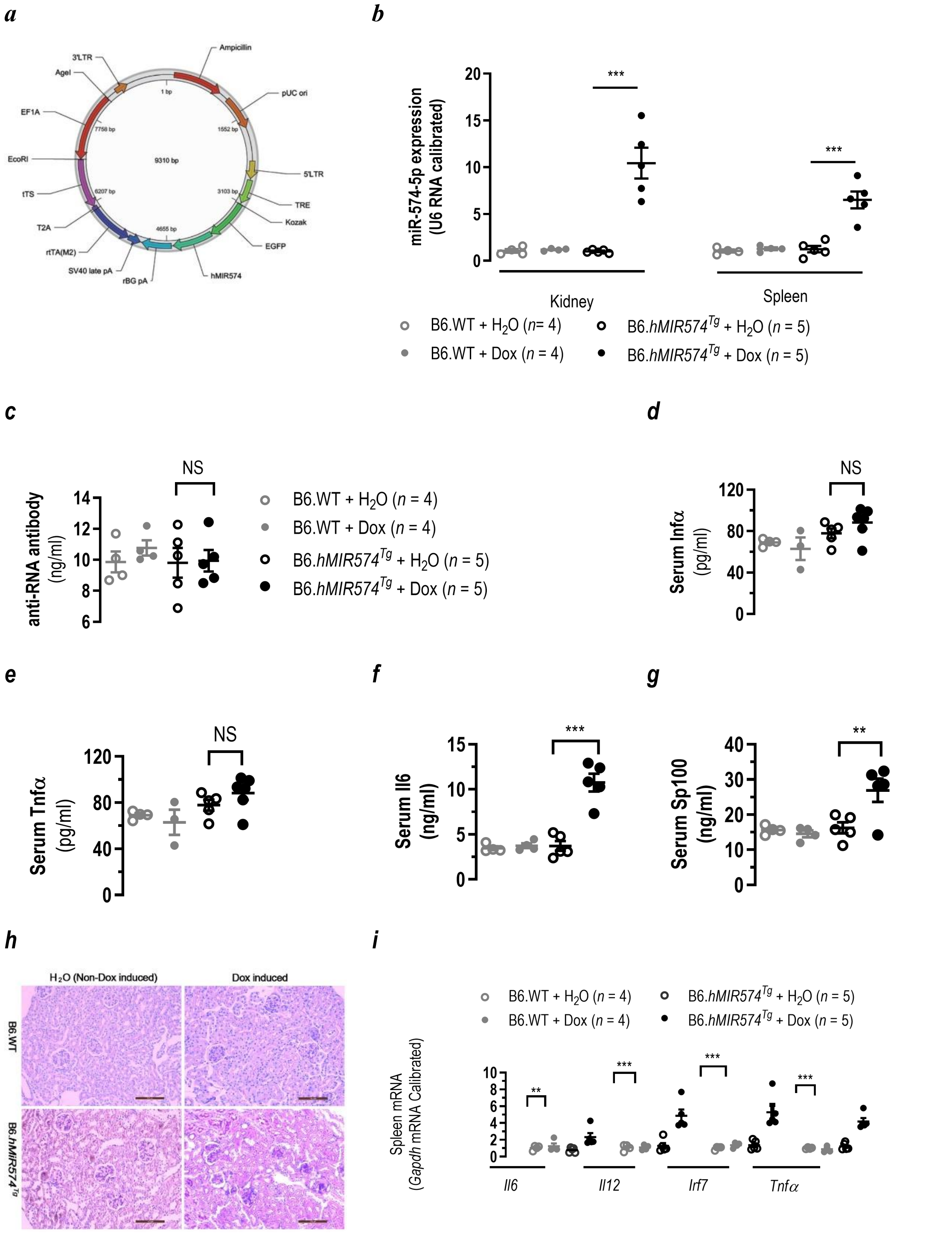


***Figure S7 Characterization of miR-574-overexpressing transgenic mice.*** (***a***). A plasmid map illustrating the construction of doxycycline-inducible human miR-574-5p expression vector pPB[Exp]-TRE>EGFP:hMIR574-EF1A>tTS:T2A:rtTA. TRE, tetracycline response element; EGFP, enhanced green fluorescent protein; Kozak, Kozak sequence; hMIR574, human *MIR574* gene; rBG pA, rabbit beta-globin poly-A; SV40 pA, SV40 poly-A; rtTA;

recombinant tetracycline-controlled transcription factor; tTS, tetracycline-controlled transcription silencer; T2A, a promoter; EF1A, elongation factor 1A. 3'LTR, 3'-long terminal repeats; 5'LTR, 5'-long terminal repeats. (***b***) The expression of miR-574-5p in female transgenic mice carrying a human miR574 transgene (B6.hMIR574$^{Tg}$) and the control (B6.WT) mice as analyzed by qPCR. The induction of transgene expression was achieved by providing the mice with 2 mg/ml Dox in the drinking water, starting from the age of 8-10 wk. Spleen and kidney samples were collected for analyses after 10 wk of Dox induction. *** $P <$ 0.001, B6.*hMIR574*$^{Tg}$ + $H_2O$ versus B6.*hMIR574*$^{Tg}$ + Dox. (***c-g***) Serum levels of anti-RNA antibody (***c***) in transgenic mice following 6 wk of Dox induction and the serum levels of Infα (***d***), Tnfα (***e***), Il6 (***f***) and Sp100 (***g***) following 10 wk of Dox induction and the un-induced controls. NS, not significant; **, $P < 0.01$, B6.*hMIR574*$^{Tg}$ + $H_2O$ versus B6.*hMIR574*$^{Tg}$ + Dox. (***h***) PAS staining showing renal tissue damage in transgenic mice following 10 wk of Dox induction but not the un-induced. Mice carrying a Dox-inducible human *MIR574* transgene and its controls were prepared as described. Results were typical for at least three mice per group. Scale bar, 100 μm. (***g***) Spleen mRNA expression of Il6, Il12, Irf7 and Tnfα in transgenic mice following 10 wk of Dox induction and the controls as analyzed by qPCR. **, $P < 0.01$, ***, $P < 0.001$, B6.*hMIR574*$^{Tg}$ + $H_2O$ versus B6.*hMIR574*$^{Tg}$ + Dox.

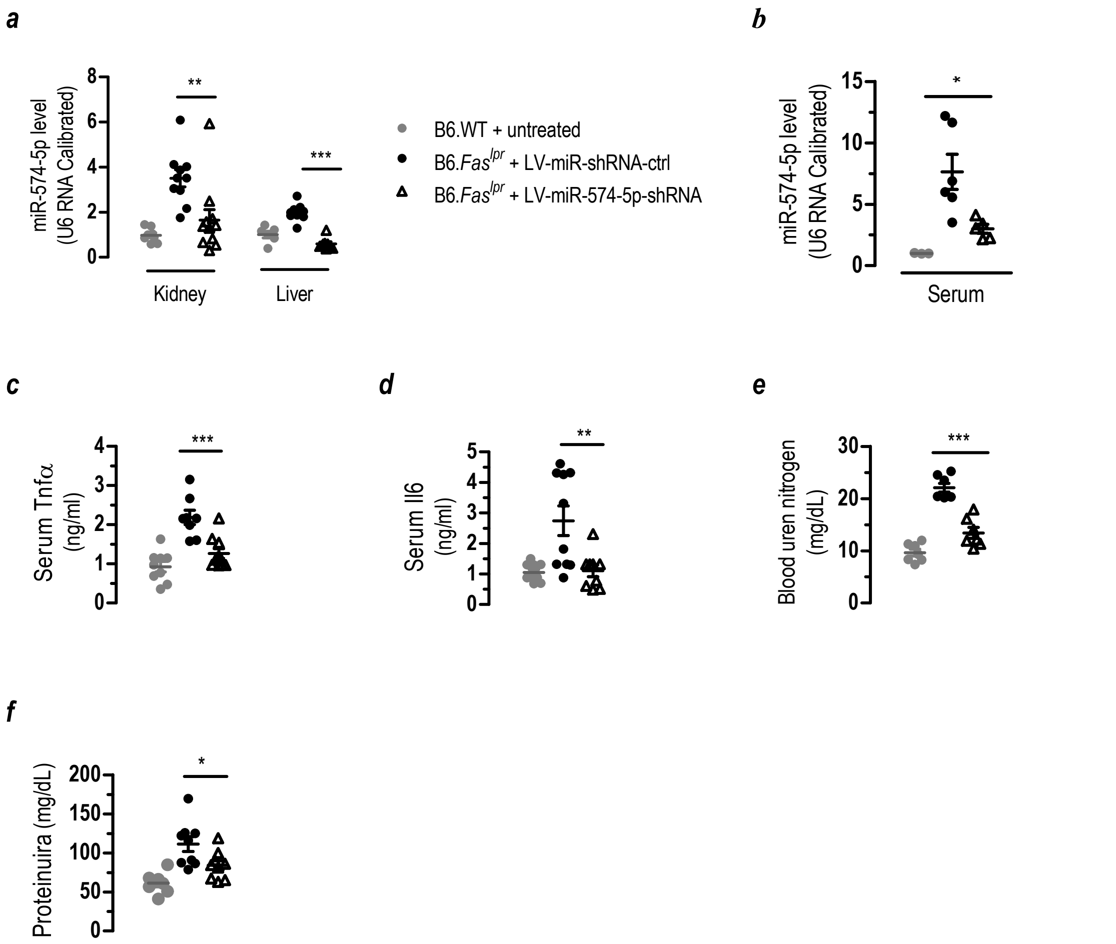


***Figure S8 Effects of lentivirus-mediated silencing of miR-574-5p on the developlemt of lupus and lupus nephritis-1.*** * $P < 0.05$, ** $P < 0.01$, *** $P < 0.001$. (***a***) Reduced miR-574-5p levels in the kidney and liver of 20-wk old B6.*Fas^lpr^* mice ($n$ = 8-10). (***b***) Reduced levels of miR-574-5p the sera of B6.$Fas^{lpr}$ mice ($n$ = 3-5). (***c-f***) Silencing of miR-574-5p in the lupus-prone B6.$Fas^{lpr}$ mice greatly reduced serum Tnfα and Il6, blood urea nitrogen and proteinuria ($n$ = 7-10.

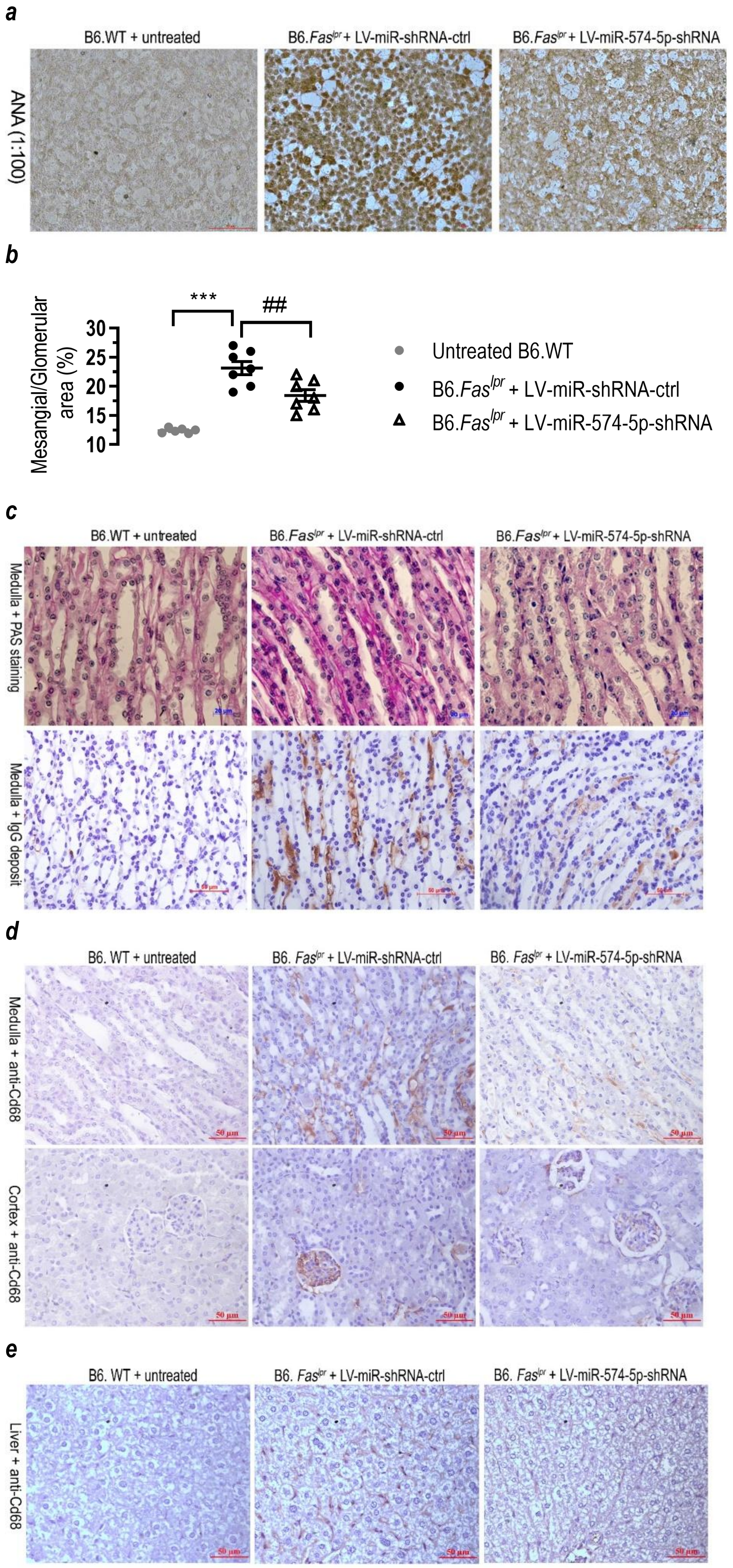
a
B6.WT + untreated
B6.Faslpr + LV-miR-shRNA-ctrl
B6.Faslpr + LV-miR-574-5p-shRNA
ANA (1:100)
b
Mesangial/Glomerular area (%)
30
25
20
15
10
***
##
Untreated B6.WT
B6.Faslpr + LV-miR-shRNA-ctrl
B6.Faslpr + LV-miR-574-5p-shRNA
c
B6.WT + untreated
B6.Faslpr + LV-miR-shRNA-ctrl
B6.Faslpr + LV-miR-574-5p-shRNA
Medulla + PAS staining
Medulla + IgG deposit
d
B6. WT + untreated
B6. Faslpr + LV-miR-shRNA-ctrl
B6. Faslpr + LV-miR-574-5p-shRNA
Medulla + anti-Cd68
Cortex + anti-Cd68
50 µm
e
B6. WT + untreated
B6. Faslpr + LV-miR-shRNA-ctrl
B6. Faslpr + LV-miR-574-5p-shRNA
Liver + anti-Cd68
50 µm

***Figure S9 Effects of lentivirus-mediated silencing of miR-574-5p on the developlemt of lupus and lupus nephritis-2.*** (***a***) Silencing of miR-574-5p in the lupus-prone B6.Fas$^{lpr}$ mice attenuated the antinuclear reactivities of sera samples from three groups of 20-wk old mice. Data were typical for 4 mice/group. HEp2 cells were incubated with sera from three groups of 20-wk old mice (B6.WT + untreated, B6.*Fas*$^{lpr}$ + LV-miR-shRNA-ctrl and B6.*Fas*$^{lpr}$ + LV-miR-574-5p-shRNA) and peroxidase-conjugated anti-mouse secondary antibody. Cells was visualized using the chromogenic substrate 3-3' diaminobenzidine. Scale bars, 50 μm. ANA, antinuclear antibody. (***b***) The ratio of mesangial to glomerular area in untreated B6.WT, LV-miR-shRNA-ctrl-treated B6.Fas$^{lpr}$ and LV-miR-574-5p-shRNA treated B6.Fas$^{lpr}$ at the age of 20-wk. *In vivo* silencing of miR-574-5p was achieved by treatment with lentiviruses carrying shRNA against miR-574-5p as described. Glomerular and mesangial area were quantitated in PAS stained kidneys as previously described (Zheng et al., 1998). ** $P < 0.001$, untreated B6.WT versus B6.*Fas*$^{lpr}$ + LV-miR-shRNA-ctrl; ## $P < 0.01$, B6.*Fas*$^{lpr}$ + LV-miR-shRNA-ctrl vs, B6.*Fas*$^{lpr}$ + LV-miR-shRNA-ctrl versus B6.*Fas*$^{lpr}$ + LV-miR-574-4p-shRNA; $n$ = 6-7. (***c-e***) Histochemical and immunohistochemical staining analyses of the renal and liver tissues in untreated B6.WT, LV-miR-shRNA-ctrl-treated B6.Fas$^{lpr}$ and LV-miR-574-5p-shRNA treated B6.Fas$^{lpr}$ at the age of 20-wk. Results were typical for at least three mice. (***c***) Histochemical staining by the PAS staining and immunohistochemistry staining by anti-IgG antibody in the renal medulla. Scale bars, 20 μm (PAS) and 50 μm (IgG) respectively. (***d***) Monocyte/macrophage infiltration in the renal medulla and cortex as determined by immunohistochemistry staining of anti-Cd68 antibody. Scale bar, 50 μm. (***e***) Macrophage infiltration in the liver as determined by immunohistochemistry staining of anti-Cd68 antibody. Scale bar, 50 μm.

#### *Supplemental Tables*

***Table S1*** The general characteristics of SLE patients and healthy controls. N/A, not assayed

| | **SLE patients (*n* = 47)** | **Controls (*n* = 18)** |
|---|---|---|
| Age (Years) | 34.77 ± 12.58 | 33.56 ± 10.98 |
| Gender (Female/Male) | 42/5 | 11/7 |
| Duration (Years) | 6.25 ± 5.27 | N/A |
| Body weight (Kg) | 60.82 ± 9.6 | 68.33 ±11.97 |
| Body mass index | 22.47 ± 3.58 | 23.8 ± 3.01 |
| Systolic blood pressure (mmHg) | 120.15 ± 16.73 | 122.72 ± 15.16 |
| Diastolic blood pressure (mmHg) | 77.15 ± 11.57 | 75.28 ± 11.98 |
| C-reactive protein (μg/ml) | 4.15 ± 2.97 | N/A (normally 0-10) |
| Albumin (g/L) | 42.20 ± 6.27 | 47.25 ± 1.6 |
| Alanine aminotransferase (U/L) | 23.29 ± 14.16 | 19.56 ± 9.07 |
| Aspartate aminotransferase (U/L) | 25.7 ± 18.27 | 19.46 ± 3.57 |
| Serum creatinine (μmol/L) | 59.82 ± 26.76 | 60.91 ± 9.73 |
| Blood urea nitrogen (mmol/L) | 5.34 ± 2.99 | 5.16 ± 0.76 |
| SLEDAI score | 4.6 ± 4.07 | N/A |

***Table S2*** *A* summary of sequencing data for six splenocyte samples.

| Sample | Sequencing Strategy | Raw Data Size (bp) | Raw Reads | Clean Data Size (bp) | Clean Reads | Clean Data Rate (%)* |
|---|---|---|---|---|---|---|
| Dotap-miRNA-1 | SE50 | 1,203,301,000 | 24,066,020 | 1,195,721,600 | 23,914,432 | 99.37 |
| Dotap-miRNA-2 | SE50 | 1,203,634,150 | 24,072,683 | 1,195,908,700 | 23,918,174 | 99.35 |
| Dotap-miRNA-3 | SE50 | 1,202,669,450 | 24,053,389 | 1,193,352,450 | 23,867,049 | 99.22 |
| Dotap-only-1 | SE50 | 1,203,061,500 | 24,061,230 | 1,194,646,000 | 23,892,920 | 99.30 |
| Dotap-only-2 | SE50 | 1,203,215,800 | 24,064,316 | 1,195,828,600 | 23,916,572 | 99.38 |
| Dotap-only-3 | SE50 | 1,203,707,700 | 24,074,154 | 1,197,699,300 | 23,953,986 | 99.50 |

*Clean Data Rate (%) = # Clean Reads/# Raw Reads

***Table S3*** Alignment statistics of reads alignment to the reference genes.

| Sample | Total Reads | Unique Match (%) | Multi-position Match (%) | Total Unmapped Reads (%) | Total Mapped Reads* (%) |
|---|---|---|---|---|---|
| Dotap-miRNA-1 | 23,914,432 | 65.17 | 6.04 | 28.79 | 71.21 |
| Dotap-miRNA-2 | 23,918,174 | 66.23 | 5.90 | 27.87 | 72.13 |
| Dotap-miRNA-3 | 23,867,049 | 65.06 | 5.94 | 29.01 | 70.99 |
| Dotap-only-1 | 23,892,920 | 64.63 | 4.93 | 30.45 | 69.55 |
| Dotap-only-2 | 23,916,572 | 65.47 | 4.89 | 29.63 | 70.37 |
| Dotap-only-3 | 23,953,986 | 65.17 | 4.87 | 29.96 | 70.04 |

* Total Mapped Reads (%) = Unique Match (%) + Multi-position Match (%)

***Table S4*** Alignment statistics of reads alignment to the reference genome.

| **Sample** | **Total Reads** | **Unique Match (%)** | **Multi-position Match (%)** | **Total Unmapped Reads (%)** | **Total Mapped Reads* (%)** |
|---|---|---|---|---|---|
| Dotap-miRNA-1 | 23,914,432 | 68.22 | 25.21 | 6.58 | 93.43 |
| Dotap-miRNA-2 | 23,918,174 | 67.76 | 25.63 | 6.61 | 93.39 |
| Dotap-miRNA-3 | 23,867,049 | 66.45 | 25.90 | 7.66 | 92.35 |
| Dotap-only-1 | 23,892,920 | 67.58 | 25.59 | 6.83 | 93.17 |
| Dotap-only-2 | 23,916,572 | 69.02 | 24.81 | 6.17 | 93.83 |
| Dotap-only-3 | 23,953,986 | 68.63 | 24.49 | 6.88 | 93.12 |

* Total Mapped Reads (%) = Unique Match (%) + Multi-position Match (%)

***Table S5*** A list of lentiviral shRNA vectors for miRNAs and mRNAs.

| **Plasmid** | **Backbone vector** | **Target gene** | **Target sequence (5' →3')** |
|---|---|---|---|
| pLV-sh-ctrl | pLentiLox3.7 | | AAAATCTCCCTAAATCATACA |
| pLV-sh-TLR7-1 | pLentiLox3.7 | TLR7 | CCTGTGAGTTAGATCTGACTA |
| pLV-sh-TLR7-2 | pLentiLox3.7 | TLR7 | GCTCTCCCTGAAAGATAACAA |
| pLV-sh-TLR8-1 | pLentiLox3.7 | TLR8 | CCGCTTAGAAATACTTGACT |
| pLV-sh-TLR8-2 | pLentiLox3.7 | TLR8 | GTCTGGATTTATCCCTTAAT |
| pLV-sh-TLR8-3 | pLentiLox3.7 | TLR8 | GATTCCATTAAGCAATACTA |
| pLV-miR-shRNA-ctrl | pLentiLox3.7 | | TTCTCCGAACGTGTCACGT |
| pLV-miR-574-5p-shRNA | pLentiLox3.7 | miR-574-5p | ACACACTCACACACACACACTCA |

***Table S6*** Plasmids and the primers used for their construction. Lic, ligation independent cloning; PA, protein A; His, histidine.

| **Plasmid** | **Backbone vector** | **cDNA template** | **Primer sequence (from 5'→ 3')** | **Gene ID** | **Insert size (bp)** |
|---|---|---|---|---|---|
| pLV-hsa-MIR574 | pLV-EF1a-MCS-IRES-Puro | human genomic DNA | Forward (Lic): *AGAGAATTCGGATCC*CCTCTGCGTTAGTGAGAAGCAG<br>Reverse (Lic): *GGGAATATTGGATCC*TCTGTCTTACAGGGACCTGCTC | MI0003581 (miRBase) | 345 |
| pTLR7$^{27-838}$-PA-His | pMT-BIP-V5-His | pFlag-TLR7 | Forward: CGCTCGGGAGATCTGGGGCTAGATGGTTTCCTAAAAC<br>Reverse: *ACCCTGGAAGTACAGGTTTTC*AGTCAGATCTAACTCACAGG | hTLR7$^{27-838}$ | 2442 |
| | | *S. aureus* | Forward: *GAAAACCTGTACTTCCAGGGTGC*TGCGCAACACGATGAAGCTC<br>Reverse: ATATCTGCAGAATTCTTATAGTTCGCGACGA | Protein A | 1221 |
| pTLR8$^{27-827}$-PA-His | pMT-BIP-V5-His | pFlag-TLR8 | Forward: CTCGCTCGGGAGATCTGAAGAAAATTTTTCTAGAAG<br>Reverse: *ACCCTGGAAGTACAGGTTTTC*AGTGACATCTGAAACACAAGTTG | hTLR8$^{27-827}$ | |
| | | *S. aureus* | Forward: *GAAAACCTGTACTTCCAGGGTGC*TGCGCAACACGATGAAGCTC<br>Reverse: ATATCTGCAGAATTCTTATAGTTCGCGACGA | Protein A | |

| | | | | | |
|---|---|---|---|---|---|
| pTlr7$^{27\text{-}839}$-PA-His | pMT-BIP-V5-His | pHA-Tlr7 | Forward: CGCTCGGGAGATCTGGGTTTCGATGGTTTCCTAA Reverse: *ACCCTGGAAGTACAGGTTTTC*TGTGAGATCTAACTCACACG | mTLR7$^{27\text{-}839}$ | 2445 |
| | | *S. aureus* | Forward: *GAAAACCTGTACTTCCAGGGTGC*TGCGCAACACGATGAAGCTC Reverse: ATATCTGCAGAATTCTTATAGTTCGCGACGA | Protein A | |
| pTlr8$^{27\text{-}818}$-PA-His | pMT-BIP-V5-His | pHA-Tlr8 | Forward: CGCTCGGGAGATCTAAAGCGAACTATTCCAGAAG Reverse: *ACCCTGGAAGTACAGGTTTTC*AGTGGTATCCGATACAC | mTLR8$^{27\text{-}818}$ | 2376 |
| | | *S. aureus* | Forward: GAAAACCTGTACTTCCAGGGTGCTGCGCAACACGATGAAGCTC Reverse: ATATCTGCAGAATTCTTATAGTTCGCGACGA | Protein A | |

***Table S7*** A list of chemically-synthesized and HPLC-purified miRNAs. Dig, digoxin; PS, phosphorothioated; s, phosphorothioate linkage.

| **Cat#** | **Supplier** | **miR** | **Sequence (5' →3')** |
|---|---|---|---|
| SS14121100 4-477 | Invitrogen, Guangzhou, China | PS-miR-16 | UsAsGsCsAsGsCsAsCsGsUsAsAsAsUsAs UsUsGsGsCsG |
| SS14121100 4-478 | Invitrogen, | PS-miR-574-5p | UsGsAsGsUsGsUsGsUsGsUsGsUsGsUsG sAsGsUsGsUsGsU |
| C26424-01 | Genscript, Nanjing, Jiangsu, China | Dig-miR-574-5p | Dig-UGAGUGUGUGUGUGAGUGUGU |
| C26424-02 | Genscript | Dig-miR-16 | Dig-UAGCAGCACGUAAAUAUUGGCG |
| E22112-01 | Genescript | Cy3-miR-574-5p | Cy3-UGAGUGUGUGUGUGAGUGUGU |

***Table S8*** A list of the antibodies used for flow cytometry, immunoblots (IB), immunoprecipitation (IP), immunohistochemistry (IHC) or immunofluorescence (IF).

| Antibody | Cat# | Dilution/ Working concentration | Supplier | Usage |
|---|---|---|---|---|
| Goat anti-Mouse IgG | 31430 | 1:5000 (IB) 1:200 (IHC) | Pierce Biotechnology, Inc., Rockford, IL, USA | IB, IHC |
| Goat anti-Rabbit IgG | 31360 | 1:5000 | Pierce | IB, IP |
| anti-β-actin | sc-47778 | 1:5000 | Santa Cruz Biotechnology, Inc., CA, USA | IB |
| anti-HLA-DRA | D154097 | 1:5000 | Sangon Biotech, Inc., Shanghai, China | IB |
| anti-HLA-C | sc-166134 | 1:1000 | Santa Cruz | IB |
| anti-GAPDH (Gapdh) | 2118 | 1:5000 | Cell Signaling, Danvers, USA | IB |
| anti-STAT1 | D120086 | 1:5000 | Sangon Biotech | IB |
| anti-STAT1 (phospho-Tyr701) | D155017 | 1:5000 | Sangon Biotech | IB |
| anti-Trim21/Ro52 | sc-21365 | 1:3000 | Santa Cruz | IB |
| anti-TRAF3 | AB60776 a | 1:1000 | Life Science Products and Services | IB |
| anti-MyD88 | D121009 | 1:5000 | Sangon Biotech | IB |
| anti-IRF8 | sc-365042 | 1:3000 | Santa Cruz | IB |
| anti-Flag | sc-807 | 1:5000 | Santa Cruz | IB, IP |
| anti-HA | sc-805 | 1:5000 | Santa Cruz | IB, IP |
| anti-His | H1029 | 1:10000 | Sigma-Aldrich, St., Louis, MO, | IB, IP |

| | | | | |
|---|---|---|---|---|
| | | | USA | |
| anti-Dig | ab420 | 1:5000 | Abcam, Cambridge, London, UK | IB, IP |
| anti-human CD19 (APC-CD19) | 302211 | 0.25 μg/$10^6$ cells | BioLegend, San Diego, CA, USA | Flow cytometry |
| anti-human CD69 (FITC-CD69) | 310903 | 0.5 μg/$10^6$ cells | BioLegend | Flow cytometry |
| anti-human TNFα (APC- TNFα) | 502913 | 1 μg/$10^6$ cells | BioLegend | Flow cytometry |
| anti-mouse B220 (APC-B220) | 103211 | 0.2 μg/$10^6$ cells | BioLegend | Flow cytometry |
| anti-mouse B220 (FITC-B220) | 103205 | 0.25 μg/$10^6$ cells | BioLegend | Flow cytometry |
| anti-mouse Cd4 (APC-Cd4) | 100411 | 0.2 μg/$10^6$ cells | BioLegend | Flow cytometry |
| anti-mouse Cd11c (PE/Cy5-Cd11c) | 117316 | 0.25 μg/$10^6$ cells | BioLegend | Flow cytometry |
| anti-Cd68 | ab955 | 1:200 | Abcam | IHC |
| anti-mouse Cd69 (APC-Cd69) | 104513 | 0.2 μg/$10^6$ cells | BioLegend | Flow cytometry |
| anti-mouse Cxcr5 (FITC-Cxcr5) | 145519 | 0.25 μg/$10^6$ cells | BioLegend | Flow cytometry |
| anti-mouse Gl7 (FITC-Gl7) | 144603 | 0.25 μg/$10^6$ cells | BioLegend | Flow cytometry |
| Pna-HRP | L7759 | 1:100 | Sigma-Aldrich | IHC |
| IgD-BIOT | 1120-08 | 1:100 | SouthernBiotech, Birmingham, AL, USA | IHC |
| anti-mouse Pd1 (PE-Pd1) | 130205 | 0.2 μg/ $10^6$ cells | BioLegend | Flow cytometry |
| anti-mouse Tnfα (FITC- Tnfα) | 506304 | 0.25 μg/ $10^6$ cells | BioLegend | Flow cytometry |
| Isotype Control | 400405 | 0.25 μg/ $10^6$ | BioLegend | Flow cytometry |

| | | | | |
|---|---|---|---|---|
| antibody (FITC) | | cells | | |
| anti-human TLR4 | ab13556 | 1:100 | Abcam | IF |
| anti-hTLR7/mTlr7 | ab45371 | 1:100 | Abcam | IF |
| anti-hTLR8 | ab53630 | 1:100 | Abcam | IF |
| anti-rabbit IgG (Alexa Fluor 488) | A11055 | 1:100 | LifeTechnology, Carlsbad, CA, USA | IF |
| anti-goat IgG (Alexa Fluor 488) | A0423 | 1:100 | Beyotime, Shanghai, China | IF |
| Anti-rabbit IgG (H+L) | Daylight680 | | Company, city, state | IB |
| Anti-human β-actin | | 1:1000 | Proteintech, Wuhan, China | IB |
| Anti-hTLR7 | 24906ss | 1:700 | Novus, Littleton, CO, USA | IB |
| Anti-hTLR8 | ab24185 | 1:500 | Abcam China, Shanghai, China | IB |

***Table S9*** A list of the primers used for RT-PCR or qPCR analyses of mRNAs and miRNAs. LNA, locked nucleic acid.

| **Organism** | **Gene** | **Gene ID** | **Primer sequence (5' → 3')** | **Amplicon (bp)** |
|---|---|---|---|---|
| human | hsa-miR-574-5p | MI0003581 | Forward: GGGGTGAGTGTGTGTGTG (LNA) Reverse: TGCGTGTCGTGGAGTC | 65 |
| human | *18S rRNA* | NR_003286.2 | Forward: CGACGACCCATTCGAACGTCT Reverse: CTCTCCGGAATCGAACCCTGA | 103 |
| human | *BAFF-R* | NM_052945.3 | Forward: TGGTCCTGGTGGGTCTGGTGA Reverse: CAGGCAGGAGCTGTGGCATCA | 157 |
| human | *BCMA* | NM_001192.2 | Forward: ATCATGTTGCAGATGGCTGGG Reverse: CCAGAGAATCGCATTCGTTCC | 174 |
| human | *CCL2* | NM_002982.3 | Forward: ACCAGCAGCAAGTGTCCCAAA Reverse: CGGAGTTTGGGTTTGCTTGTC | 129 |
| human | *CD19* | NM_001770.5 | Forward: TATTGTCACCGTGGCAACCTG Reverse: CCAAAGTCACAGCTGAGACCT | 115 |
| human | *CD74* | NM_001025158.2 | Forward: AGCATCACTCCCCAAGGAAGA Reverse: TGTGAACCATGGCCCTGAAA | 113 |
| human | *GAPDH* | NM_001256799.2 | Forward: CCAGCAAGAGCACAAGAGGAA Reverse: ATGGTACATGACAAGGTGCGG | 157 |

| | | | | |
|---|---|---|---|---|
| human | *HLA-C* | M26431 | Forward: CCATGAGGTATTTGTGGACCG Reverse: TCTCGGACTCTCGTCGTCG | 122 |
| human | *HLA-DRA* | NM_019111.4 | Forward: ATACTCCGATCACCAATGTACCT Reverse: GACTGTCTCTGACACTCCTGT | 173 |
| human | *IL6* | NM_000600 | Forward: ACAAATTCGGTACATCCTCGAC Reverse: GAATCCAGATTGGAAGCATCC | 154 |
| human | *IL8* | NM_000584.3 | Forward: CCAGGAAGAAACCACCGGAAG Reverse: TGGTCCACTCTCAATCACTCTCAG | 221 |
| human | *IRF7* | NM_001572.3 | Forward: GTGATGCTGCGGGATAACTC Reverse: ATGTGTGTGTGCCAGGAATG | 174 |
| human | *IRF8* | NM_002163.2 | Forward: ACGCTGTGCTTTGAATAAGAGC Reverse: TCCTCAGGAACAATTCGGTAAAC | 102 |
| human | *NR4A1* | NM_001202233.1 | Forward: CCCTGAAGTTGTTCCCCTCAC Reverse: GCCCTCAAGGTGTGGAGAAG | 110 |
| human | *OLR1* | NM_001172632.1 | Forward: GCTATTCTTTGTCACTTGGG Reverse: AAATGTTGACATAAAGGTGC | 120 |
| human | *SFRS1* | NM_001078166.1 | Forward: TCTCTGGACTGCCTCCAAGT Reverse: GGCTTCTGCTACGACTACGG | 473 |

| | | | | |
|---|---|---|---|---|
| human | *TACI* | NM_012452.2 | Forward: TGCTGGGTACCTGCATGTCCT Reverse: ACAGCTGATGCAGTCCCTCAG | 140 |
| human | *TLR7* | NM_016562.3 | Forward: TGTGCATCAAGAGGCTGCAGA Reverse: GGGCACATGCTGAAGAGAGTT | 587 |
| human | *TLR8* | NM_138636.5 | Forward: CAGAGCATCAACCAAAGCAA; Reverse: GCTGCCGTAGCCTCAAATAC | 184 |
| human | *TSC1* | NM_000368.4 | Forward: CAACAAGCAAATGTCGGGGAG Reverse: CATAGGGCCACGGTCAGAA | 114 |
| mouse | mmu-miR-574-5p | MI0005518 | RT: GTCGTATCCAGTGCGTGTCGTGGAGTCGGCAATTGCACTGGATACGACTACACAC Forward: GGGGTGAGTGTGTGTGTG (LNA) Reverse: TGCGTGTCGTGGAGTC | 65 |
| mouse | *18S rRNA* | NR_003278.3 | Forward: CGACGACCCATTCGAACGTCT Reverse: CTCTCCGGAATCGAACCCTGA | 103 |
| mouse | *Adar* | NM_019655.3 | Forward: TGAGCATAGCAAGTGGAGATACC Reverse: GCCGCCCTTTGAGAAACTCT | 95 |

| | | | | |
|---|---|---|---|---|
| mouse | *Cd86* | NM_019388 | Forward: CTGGACTCTACGACTTCACAATG Reverse: AGTTGGCGATCACTGACAGTT | 131 |
| mouse | *Cd274* | NM_021893 | Forward: GCTCCAAAGGACTTGTACGTG Reverse: TGATCTGAAGGGCAGCATTTC | 238 |
| mouse | *Csprs* | NM_114564 | Forward: CCATCACCTGCTTTGTATCTCC Reverse: GCCCCAAATACCAGCCAAC | 111 |
| mouse | *Cxcl10* | NM_021274 | Forward: CCAAGTGCTGCCGTCATTTTC Reverse: GGCTCGCAGGGATGATTTCAA | 157 |
| mouse | *Ddx58* | NM_172689 | Forward: AAGAGCCAGAGTGTCAGAATCT Reverse: AGCTCCAGTTGGTAATTTCTTGG | 106 |
| mouse | *Dhx58* | NM_030150 | Forward: GGAAGTGATCTTACCTGCTCTGG Reverse: TTGCCTCTGTCTACCGTCTCT | 123 |
| mouse | *Eif2ak2* | NM_011163 | Forward: ATGCACGGAGTAGCCATTACG Reverse: TGACAATCCACCTTGTTTTCGT | 194 |
| mouse | *Gapdh* | NM_001289726.1 | Forward: CGGAGTCAACGGATTTGGTCGTAT Reverse: AGCCTTCTCCATGGTGGTGAAGA | 307 |

| | | | | |
|---|---|---|---|---|
| | | | C | |
| mouse | *GzmA* | NM_010370 | Forward: TGCTGCCCACTGTAACGTG Reverse: GGTAGGTGAAGGATAGCCACAT | 207 |
| mouse | *GzmB* | NM_013542 | Forward: CCACTCTCGACCCTACATGG Reverse: GGCCCCCAAAGTGACATTTATT | 142 |
| mouse | *Hnf1b* | NM_009330 | Forward: CACCAAGCCGGTTTTCCATAC Reverse: GGAGTGTCATAGTCGTCGCC | 96 |
| mouse | *Il-12 P40* | NM_001303244.1 | Forward: GACGTCAGCTGGGAGTACCCT Reverse: GCGCTGGATTCGAACAAAGA | 81 |
| mouse | *Ifi47* | NM_008330 | Forward: TCTCCAGAAACCCTCACTGGT Reverse: TCAGCGGATTCATCTGCTTCG | 200 |
| mouse | *Ifih1* | NM_001164477 | Forward: AGATCAACACCTGTGGTAACACC Reverse: CTCTAGGGCCTCCACGAACA | 107 |
| mouse | *Ifit1* | NM_008331 | Forward: CTGAGATGTCACTTCACATGGAA Reverse: GTGCATCCCCAATGGGTTCT | 117 |
| mouse | *Ifit3* | NM_010501 | Forward: CCTACATAAAGCACCTAGATGGC Reverse: ATGTGATAGTAGATCCAGGCGT | 148 |
| mouse | *Igtp* | NM_018738 | Forward: CTCATCAGCCCGTGGTCTAAA Reverse: | 102 |

| | | | | |
|---|---|---|---|---|
| | | | CACCGCCTTACCAATATCTTCAA | |
| mouse | *Il6* | NM_001314054.1 | Forward: TCCTCTGGTCTTCTGGAGTA Reverse: CTTAGCCACTCCTTCTGTGA | 160 |
| mouse | *Irf1* | NM_008390.2 | Forward: ATGCCAATCACTCGAATGCG Reverse: TTGTATCGGCCTGTGTGAATG | 197 |
| mouse | *Irf3* | NM_016849.4 | Forward: CACAGATGGCTGACTTTGGC Reverse: GGCCATCAAATAACTTCGGTAG | 300 |
| mouse | *Irf4* | NM_013674.2 | Forward: CTCTTCAAGGCTTGGGCATT Reverse: TGCTCCTTTTTTGGCTCCCT | 201 |
| mouse | *Irf7* | NM_001252600 | Forward: CCCCAGCCGGTGATCTTTC Reverse: CACAGTGACGGTCCTCGAAG | 125 |
| mouse | *Irf7* | NM_001252601.1 | Forward: TTGGGCAAGACTTGTCAGCA Reverse: GTCAAGGCCACTGACCCAG | 213 |
| mouse | *Klrd1* | NM_010654 | Forward: TCTAGGATCACTCGGTGGAGA Reverse: CACTTGTCCAGGCAAACACAG | 185 |
| mouse | *Ly6a* | NM_010738 | Forward: AGGAGGCAGCAGTTATTGTGG Reverse: CGTTGACCTTAGTACCCAGGA | 114 |
| mouse | *Mx1* | NM_010846 | Forward: GACCATAGGGGTCTTGACCAA Reverse: | 182 |

| | | | | |
|---|---|---|---|---|
| | | | AGACTTGCTCTTTCTGAAAAGCC | |
| mouse | *Nlrc3* | NM_001081280 | Forward: CAGATTGGTAACAAAGGAGCCA Reverse: CGTTCGGTTTATCTTCAGAGCA | 141 |
| mouse | *Oas1a* | NM_145211 | Forward: GCCTGATCCCAGAATCTATGC Reverse: GAGCAACTCTAGGGCGTACTG | 217 |
| mouse | *Papd7* | NM_001116913 | Forward: AGGTGGTGAAACGGATCGAAA Reverse: CCAGGTCTATGTCACTTGTTGGA | 114 |
| mouse | *Rfc3* | NM_027009 | Forward: CAGAGTGCAGCAATACCCTTT Reverse: ACAATAGCATTTGCGGTCTCC | 90 |
| mouse | *Sp100* | NM_013673 | Forward: AGCTACAACCACAGTCCCCT Reverse: TCCTGTCCTTTTCCGTCTTCTAA | 133 |
| mouse | *Sp110* | NM_030194 | Forward: ATGAAGGTGAACATCGCCTATG Reverse: GGACAGAGGGACCAGATTTTG | 132 |
| mouse | *Sp140* | NM_001013817.2 | Forward: GGTGGAGATCGCAAGTGCCAT Reverse: CTCCACTGGGACCAGGTTCTT | 127 |
| mouse | *Socs1* | NM_009896 | Forward: CTGCGGCTTCTATTGGGGAC Reverse: AAAAGGCAGTCGAAGGTCTCG | 216 |
| mouse | *Stat1* | NM_001205314 | Forward: GCTGCCTATGATGTCTCGTTT Reverse: | 124 |

| | | | | |
|---|---|---|---|---|
| | | | TGCTTTTCCGTATGTTGTGCT | |
| mouse | *Stat2* | NM_019963 | Forward: CTGAAGGACGAACAGGATGTC Reverse: CAGGGTGGTTAATCGGCCAA | 186 |
| mouse | *Themis* | NM_178666 | Forward: AGTCACCATGTAGACAGACCC Reverse: GTGGCCCATGCTTGCTCTT | 124 |
| mouse | *Tlr3* | NM_126166 | Forward: GTGAGATACAACGTAGCTGACTG Reverse: TCCTGCATCCAAGATAGCAAGT | 162 |
| mouse | *Tlr7* | NM_133211 | Forward: ATGTGGACACGGAAGAGACAA Reverse: GGTAAGGGTAAGATTGGTGGTG | 207 |
| mouse | *Tnfα* | NM_001278601.1 | Forward: CGTGGAACTGGCAGAAGAGG Reverse: CTGCCACAAGCAGGAATGAG | 101 |
| mouse | *Trim21/Ro52* | NM_009277 | Forward: GGGAGGAGGTCACCTGTTCTA Reverse: GGCACTCGGGACATGAACTG | 132 |
| mouse | *Trim25* | NM_009546 | Forward: ATGGCTCAGGTAACAAGGGAG Reverse: GGGAGCAACAGGGGTTTTCTT | 107 |
| mouse | *U6* | NR_004394.1 | RT: CGCTTCACGAATTTGCGTGTCAT Forward: GCTTCGGCAGCACATATACTAAAAT (LNA) Reverse: CGCTTCACGAATTTGCGTGTCAT | 89 |

| | | | | |
|---|---|---|---|---|
| mouse | *Ube216* | NM_019949 | Forward: GTGGCGAAAGAGCTGGAGAG Reverse: GGGGAAATCAATCCGCACTTG | 156 |
| mouse | *Usp18* | NM_011909 | Forward: AGAGTTAGCAAGCTCCGACAT Reverse: TGAGGTGAATGGTCAAGGTTTG | 107 |

***Table S10*** A list for the ELISA kits used.

| **ELISA kit** | **Cat#** | **Supplier** |
|---|---|---|
| Human IL6 | VAL102 | R&D Systems China Co., Ltd, Minneapolis, MN, USA |
| Human IFNα | BMS216INST | eBioscience Inc., San Diego, CA, USA |
| Human IFNγ | VAL104 | R&D |
| Human TNFα | VAL105 | R&D |
| Mouse IgG | 88-50400 | eBioscience |
| Mouse IgG1 | 88-50410 | eBioscience |
| Mouse IgG2b | 88-50430 | eBioscience |
| Mouse IgM | 88-50470 | eBioscience |
| Mouse Il6 | E09358-1646 | eBioscience |
| Mouse Ifnα | E09479-1647 | eBioscience |
| Mouse Ifnβ | 439407 | BioLegend, San Diego, CA, USA |
| Mouse Ifnγ | 430807 | BioLegend |
| Mouse Tnfα | E09479-1648 | eBioscience |
| Mouse Trim21/Ro52 | EA-5203 | Signosis Inc, Santa Clara, CA, USA |
| Mouse Sp100 | SU-B29795 | Shanghai Enzyme-linked Biotechnology Co., Ltd., Shanghai, China |
| Mouse anti-dsDNA antibody | 5110 | Alpha Diagnostic International, San Antonio, TX, USA |
| Mouse anti-dsDNA antibody | CK-E20444 | Shanghai Enzyme-linked Biotechnology Co., Ltd. |
| Mouse anti-RNA antibody | CK-E28508 | Shanghai Enzyme-linked Biotechnology Co., Ltd. |
